# Beyond the biosynthetic gene cluster paradigm: Genome-wide co-expression networks connect clustered and unclustered transcription factors to secondary metabolic pathways

**DOI:** 10.1101/2020.04.15.040477

**Authors:** Min Jin Kwon, Charlotte Steiniger, Timothy C. Cairns, Jennifer H. Wisecaver, Abigail Lind, Carsten Pohl, Carmen Regner, Antonis Rokas, Vera Meyer

**Affiliations:** Chair of Applied and Molecular Microbiology, Institute of Biotechnology, Technische Universität Berlin, Berlin, Germany; Department of Biochemistry, Center for Plant Biology, Purdue University, West Lafayette, Indiana, USA; Department of Biological Sciences, Vanderbilt University, Nashville, Tennessee, USA; Gladstone Institute for Data Science and Biotechnology, San Francisco, California, USA; Department of Biomedical Informatics, Vanderbilt University School of Medicine, Nashville, Tennessee, USA

**Keywords:** filamentous fungi, *Aspergillus niger*, secondary metabolite gene clusters, gene co-expression, correlation network, natural product, specialized metabolism, genetic network, gene regulation

## Abstract

Fungal secondary metabolites are widely used as therapeutics and are vital components of drug discovery programs. A major challenge hindering discovery of novel secondary metabolites is that the underlying pathways involved in their biosynthesis are transcriptionally silent in typical laboratory growth conditions, making it difficult to identify the transcriptional networks that they are embedded in. Furthermore, while the genes participating in secondary metabolic pathways are typically found in contiguous clusters on the genome, known as biosynthetic gene clusters (BGCs), this is not always the case, especially for global and pathway-specific regulators of pathways’ activities. To address these challenges, we used 283 genome-wide gene expression datasets of the ascomycete cell factory *Aspergillus niger* generated during growth under 155 different conditions to construct two gene co-expression networks based on Spearman’s correlation coefficients (SCC) and on mutual rank-transformed Pearson’s correlation coefficients (MR-PCC). By mining these networks, we predicted six transcription factors named MjkA – MjkF to concomitantly regulate secondary metabolism in *A. niger*. Over-expression of each transcription factor using the Tet-on cassette modulated production of multiple secondary metabolites. We found that the SCC and MR-PCC approaches complemented each other, enabling the delineation of global (SCC) and pathway-specific (MR-PCC) transcription factors, respectively. These results highlight the great potential of co-expression network approaches to identify and activate fungal secondary metabolic pathways and their products. More broadly, we argue that novel drug discovery programs in fungi should move beyond the BGC paradigm and focus on understanding the global regulatory networks in which secondary metabolic pathways are embedded.

**Importance:** There is an urgent need for novel bioactive molecules in both agriculture and medicine. The genomes of fungi are thought to contain vast numbers of metabolic pathways involved in the biosynthesis of secondary metabolites with diverse bioactivities. Because these metabolites are biosynthesized only under specific conditions, the vast majority of fungal pharmacopeia awaits discovery. To discover the genetic networks that regulate the activity of secondary metabolites, we examined the genome-wide profiles of gene activity of the cell factory *Aspergillus niger* across hundreds of conditions. By constructing global networks that link genes with similar activities across conditions, we identified six global and pathway-specific regulators of secondary metabolite biosynthesis. Our study shows that elucidating the behavior of the genetic networks of fungi under diverse conditions harbors enormous promise for understanding fungal secondary metabolism, which ultimately may lead to novel drug candidates.

## Introduction

Fungal secondary metabolites (SMs) are bioactive, usually small molecular weight compounds, which have restricted taxonomic distribution and are produced at specific stages of growth and development^1^. The most well-known clinical applications of these molecules include antibiotics, cholesterol-lowering agents, and immunosuppressants (e.g., penicillin, statins, and cyclosporins, respectively)^2^. However, they also play an important role in drug discovery programs, with recently marketed therapeutics consisting of either fungal SMs or their semi-synthetic derivatives^3^. In contrast to these contributions to human welfare, fungal SMs include potent carcinogenic crop contaminants^4^, and the mycotoxin-producing capacity of commonly used fungal cell factories in food or biotechnological processes is often either unknown^5^ or underestimated^6^. Moreover, plant-infecting fungi deploy numerous SMs as virulence factors that facilitate successful infection^7^, ultimately destroying enough food for 10% of the human population per year^8^. Improved understanding of the genetic, molecular, and biochemical aspects of fungal secondary metabolism thus promises to drive novel medical breakthroughs, while also insuring improvements in global food safety and security^9^.

A common feature of SM-producing fungi is that the genes required for producing a single secondary metabolite are often found in contiguous clusters on the genome, which may facilitate both horizontal gene transfer of SMs and enable epigenetic regulation via chromatin remodelling^1,10^. Biosynthetic gene clusters (BGCs) typically consist of a gene encoding a core biosynthetic enzyme, most commonly a non-ribosomal peptide synthetase (NRPS), polyketide synthase (PKS), or terpene cyclase, which is responsible for the first metabolic step in product synthesis^11^. Additionally, BGCs include genes encoding so called ‘tailoring’ enzymes, such as P450 monooxygenases or methyltransferases, which modify the molecule produced by the core enzyme^11,12^. Moreover, many BGCs contain either putative membrane transporter-encoding genes, which are required for metabolite efflux from the cell in some^13^, but not all^14^, cases, or additional so called ‘resistance’ genes, which are necessary for the detoxification/self-protection against the produced molecules^15^.

Most BGCs are transcriptionally silent under standard laboratory and industrial cultivation conditions, which is a major challenge to the discovery of their cognate molecules^16^. Interestingly, many BGCs also contain transcription factor (TF)-encoding genes that regulate their activity^11,12,17^. In several instances, these TF-encoding genes have been over-expressed to activate transcription of the respective BGC, ultimately leading to discovery of novel SMs^13,18–21^. However, this strategy cannot be used for the approximately 40% of fungal BGCs that a resident TF^17^.

An alternative approach to engineering SM over-producing isolates has been to identify and genetically target global regulators of multiple BGCs. These include epigenetic regulators, notably components of the heterotrimeric velvet complex, which links development, light responses, and SM production in ascomycetes^22^. Alternatively, globally acting TFs that coordinate SM biosynthesis with differentiation (e.g., BrlA/StuA) and responses to environmental stimuli, such as pH (PacC) or nitrogen availability (AreA), can be activated using molecular approaches for elevated natural product biosynthesis^1,17,23^. A limitation to these strategies, however, is that all global regulators discovered to date activate only a fraction of the predicted BGCs in a single genome. For example, deletion of genes predicted to encode the methyltransferase LaeA, which is thought to silence BGC expression by the formation of transcriptionally silent heterochromatin, increased expression of 7 out of 17 BGCs in the biomass-degrading fungus *Trichoderma reesei* and 13 out of 22 BGCs analysed in the human pathogen *Aspergillus fumigatus*^24,25^.

A final confounding factor in understanding and functionally analysing fungal BGCs and their products is that there is considerable variation to the degree to which core, tailoring, transport, and regulatory genes are contiguously clustered in fungal genomes^10^. This includes so called ‘partial’ clusters in which some genes encoding biosynthetic enzymes and transporters are not physically linked with other clustered genes^26,27^, ‘superclusters’ in which two or more NRPS/PKS encoding genes reside in close physical proximity^28,29^, and SM biosynthetic genes which are not contiguously clustered^30^.

Consequently, innovative strategies are required to both discover novel transcriptional activators of BGCs and to accurately delineate their boundaries. Over the past several years, an approach that has gained considerable interest has been the utilisation of co-expression networks to analyse BGCs, for example during laboratory culture of industrial isolates^29,31^ or during infectious growth of plant-infecting fungi^32^. A limitation to these studies, however, was the relatively small number of conditions tested (up to several dozen), which resulted in the inability to detect the transcriptional activity of numerous BGCs. To overcome this limitation, we recently conducted a meta-analysis of 283 microarray datasets covering 155 different cultivation conditions for the biotechnologically exploited cell factory *Aspergillus niger*. This data collection covers a diverse range of environmental conditions and genetic perturbations and was used to construct a global gene co-expression network based on Spearman’s correlation coefficient (SCC)^33^. We found that 53 out of the 81 predicted BGC core genes in *A. niger* are expressed in at least one out of the 155 conditions, and we were able to delineate the boundaries of numerous BGCs, including, for example, the partial cluster required for biosynthesis of the siderophore triacetyl fusarinine C.

Our analysis also suggested that only a minority of BGCs are co-expressed with their resident TF; specifically, from the 25 out of the 53 expressed BGCs that contained a TF, only 8 BGCs were co-expressed with their respective TF. However, we were able to use this network to successfully predict global TFs that, independent of their physical location on the genome, regulate multiple BGCs. This relied on the so-called ‘guilt-by-association’ principle, whereby genes that are part of similar (or the same) biosynthetic pathways or genetic networks tend to have highly comparable patterns of gene expression. We functionally analyzed two of these co-expressed TFs (MjkA, MjkB) by generating loss-of-function and gain-of-function *A. niger* mutants, and could indeed demonstrate that their overexpression modulated (either indirectly or directly) the transcriptional activity of 45 (MjkA) and 43 (MjkB) BGC core genes, respectively^33^.

Despite the utility of co-expression network analyses, there are several possible limitations to the construction of transcriptional networks based on correlation coefficients such as Spearman or Pearson. In these networks, correlation coefficients are used as weighted edges to connect genes (nodes). One major challenge when constructing these networks is determining the edge weight threshold below which correlation coefficients are excluded from the network, with the goal being to remove non-biologically relevant gene associations. We have previously used *in silico* data randomization experiments to test the likely threshold of biologically meaningful co-expression based on Spearman^33^, however, it is still likely that for many BGCs, the correlation coefficient cut-off chosen (ρ ≥ |0.5|) may be unnecessarily stringent, resulting in false negative co-expression relationships for BGCs. Additionally, average correlation coefficients can vary by gene function and input data^34^. Importantly, in the case of BGC genes that are only expressed under few or only one specific environmental condition, it is likely that the expression vector for a given BGC gene will be sparse, and therefore more likely to artificially correlate with other rarely-expressed genes rather than with genes with a functional link.

To overcome these challenges, in this study we reanalyzed the existing *A. niger* transcriptome dataset with a specific focus on *A. niger* BGCs. Firstly, we generated gene expression modules based on a mutual rank approach, which can capture functional relationships for rarely-expressed secondary metabolism genes^34,35^, as we have previously shown in analyses of secondary metabolism in plants^36^. We compared this mutual rank strategy with our existing Spearman co-expression datasets, and by integrating both approaches generated a shortlist of six TF-encoding genes (including *mjkA* and *mjkB*), which we hypothesized may regulate multiple BGCs. Functional analyses of these genes by overexpression using the Tet-on gene switch revealed they play multiple roles in growth, development and pigment formation of *A. niger* as assayed by standard growth tests on medium agar plates and in shake flasks. Moreover, metabolomic profiling revealed a change in metabolite patterns of analyzed overexpression strains. Finally, by *in silico* analysis we generated a list of predicted molecules and associated them with putative BGCs. The methods and resources developed in this study will thus enable the efficient activation of fungal SMs for novel drug discovery programs and other studies. More broadly, our general approach holds potential for deciphering the global regulatory network governing BGCs and secondary metabolic pathways in fungi.

## Results

### Mining co-expression networks to identify biosynthetic and regulatory modules

Using the SCC approach, we previously estimated the global transcriptional activity of *A. niger* BGCs amongst the 283 microarray experiments by assessing gene expression of the predicted core enzyme^33^. These data highlighted that BGC expression varies considerably, with some core enzymes transcriptionally deployed during several dozen experiments, others expressed in >5, and 28 not expressed under any condition^33^. We reasoned that this microarray meta-analysis was also a promising resource for further interrogation of BGCs using the MR-PCC approach. In doing so, modules of co-expressed genes were determined using three different exponential decay rates (see Materials and Methods). Each different exponential decay rate produces modules with different qualities; NET25, the most relaxed threshold, has the largest modules, while NET05, the most stringent threshold, has the smallest modules. In addition, the NET10 exponential decay rate produces modules smaller than the NET25 modules and larger than the NET05 modules.

In total, there were 2,041 modules recovered from the NET25 network, 2,944 modules recovered from the NET10 network, and 2,999 modules recovered from the NET05 network (Supplemental Table 1). The median module size for the NET25, NET10, and NET05 networks was 11, 7, and 5 genes, respectively. Of the 78 predicted BGCs comprising in total 81 core genes in the *Aspergillus niger* genome, 43 predicted BGCs had one or more genes recovered within a single module (Supplemental Table 2). These 43 BGCs had varying levels of co-expression. For some BGCs, such as the fumonisin-producing BGC, most genes in the gene cluster are co-expressed at high levels (**Figure 1**A). For others, either a small subset of the genes in the BGC were not co-expressed (e.g., BGC 34; **Figure 1**B) or only a small fraction of genes was co-expressed (e.g., BGC 38, where only 6 / 22 genes in the BGC were co-expressed; **Figure 1**C). Notably, 7 genes in BGC 38 were co-expressed with 10 genes from BGC 34, thus forming a metamodule (**Figure 1**D). This metamodule consisted in total of 50 genes, including one core gene (FAS) and two TFs from BGC 34 and two core genes (PKS, NRPS) from BGC 38. Concordantly, we could also identify co-expression between BGC 34 and BGC 38 cluster members via the SCC approach. Notably, Multigene BLAST showed that BGC 34 and 38 are conserved in black *Aspergilli* (Supplemental Figure 1). Both clusters belong to a large SCC sub-network comprised of 1,804 genes (**Figure 2**), which is the largest gene co-expression sub-network with BGC genes based on the Spearman rank coefficient ρ ≥ |0.5|. This sub-network included many TFs that are not physically located inside BGCs or are co-expressed with non-resident BGC genes.

**Figure 1:**
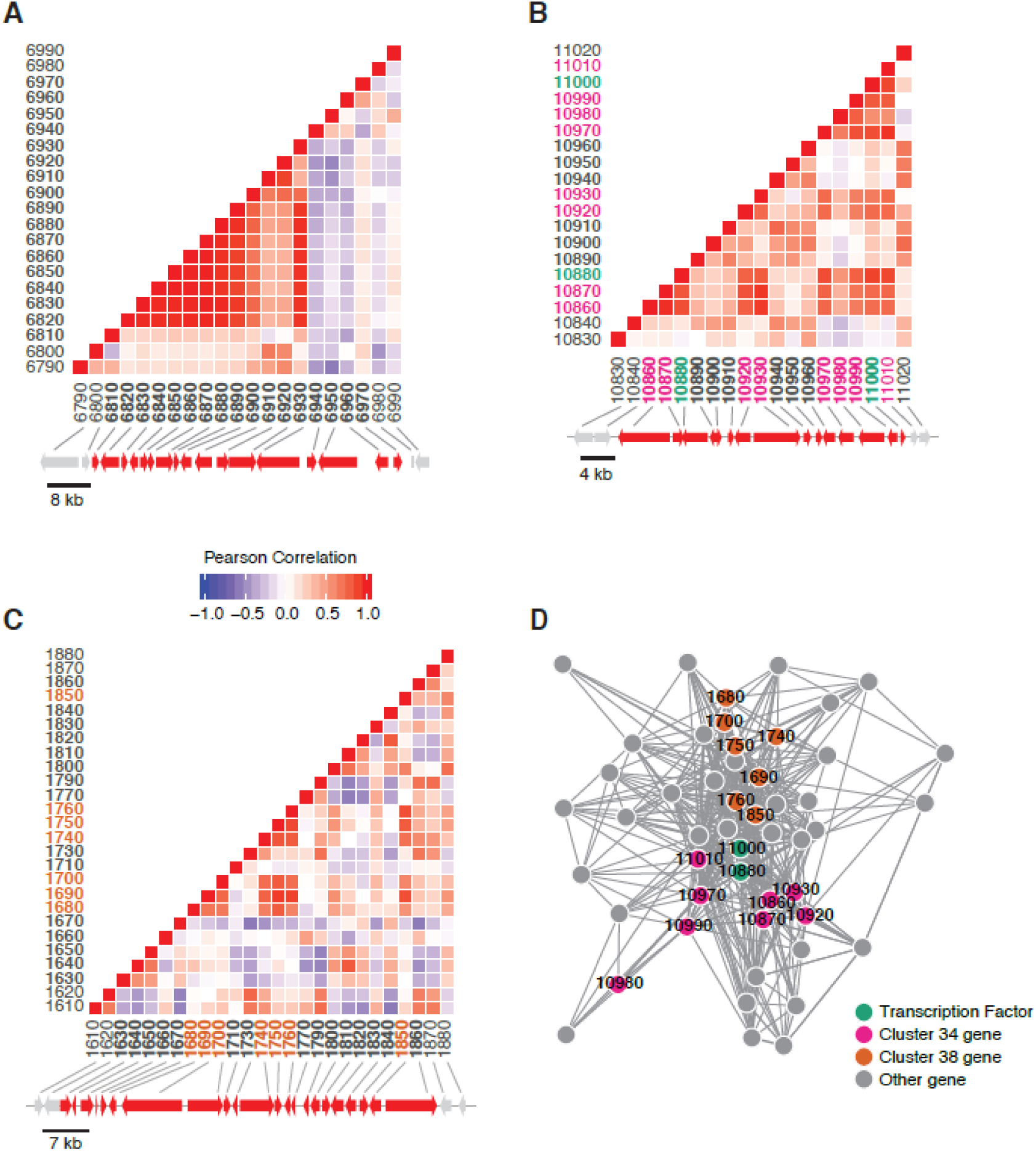
Heatmap depicting the Pearson’s correlation of co-expression of genes within three canonical BGCs. Across all panels, genes within the canonical cluster are bolded in the heatmap and colored red in the accompanying chromosome segment. Two flanking genes are included on either side and colored grey. Gene names have been abbreviated. (A) A significant fraction of genes within the fumonisin metabolic gene cluster are co-expressed. (B) Co-expression of predicted BGC 34, which contains two transcription factors. Both are colored green in the heatmap, and other clustered genes recovered in the metamodule are colored pink. (C) A small fraction of genes within predicted BGC 38 are co-expressed. Genes are color coded in the heatmap as in (A); genes recovered in a metamodule are colored orange. (D) Network map of transcription factor metamodule containing all genes co-expressed with both transcription factors across all three network analyses. Nodes in the map represent genes, and edges connecting two genes represent the weight (transformed MR score) for the association. Transcription factors are colored green. Other genes present in BGC 34 are colored pink. Genes present in BGC 38 are colored orange.

**Figure 2:**
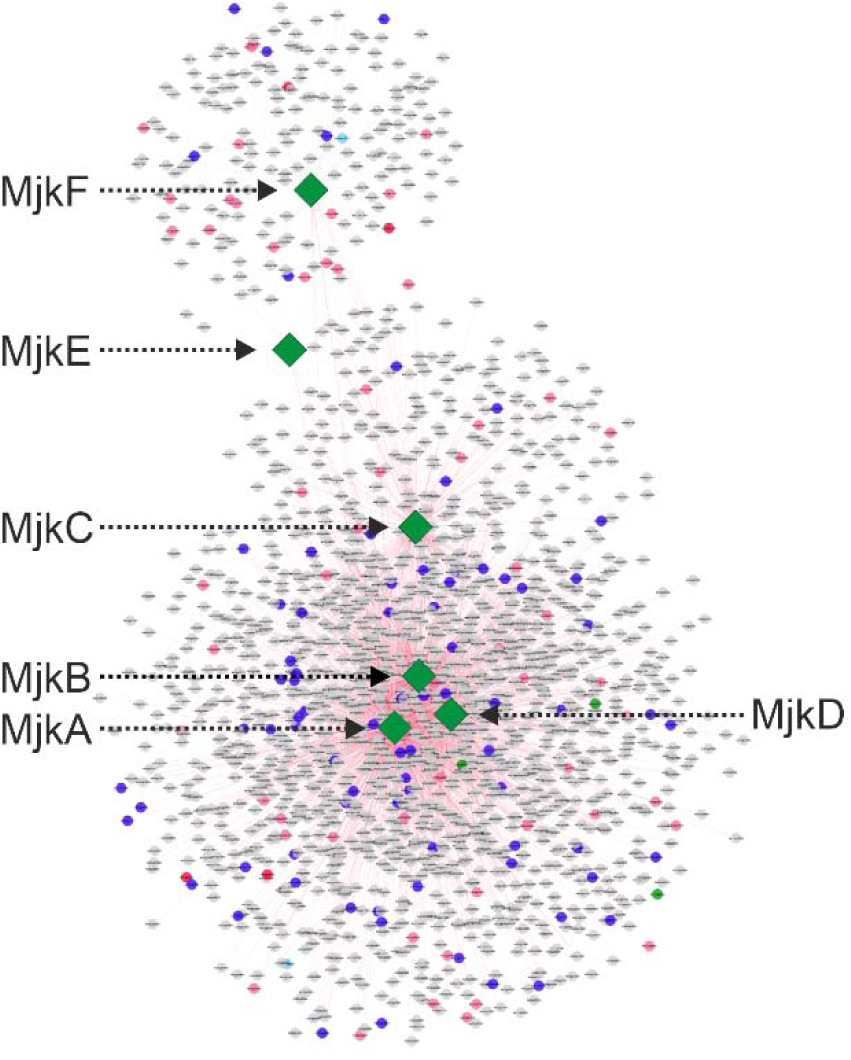
The largest Spearman sub-network containing predicted BGC core and tailoring genes (highlighted in pink) as well as transcription factors (highlighted in blue). The six transcription factors studied by molecular analyses in this study (MjkA-F) are indicated in green.

It has been speculated over the last decades that BGC resident TFs may co-regulate gene expression at more than one BGC^1,17^. Both co-expression network approaches supported this hypothesis for *A. niger*, as evidenced by the co-expression of two TFs residing in BGC 34 (An08g11000 and An08g10880, chromosome 1) with multiple genes at BGC 38 (chromosome 8), including the predicted NRPS (**Figure 3**). This was especially interesting given that (i) BGC 38 does not contain a predicted TF; (ii) both these BGCs are present in 22 (BGC 34) or 24 (BGC 38) of 83 analyzed genomes of the genus *Aspergillus*, and (iii) BGC 38 is in close proximity to the functionally characterized BGC 39 necessary for azanigerone production^37^.

**Figure 3:**
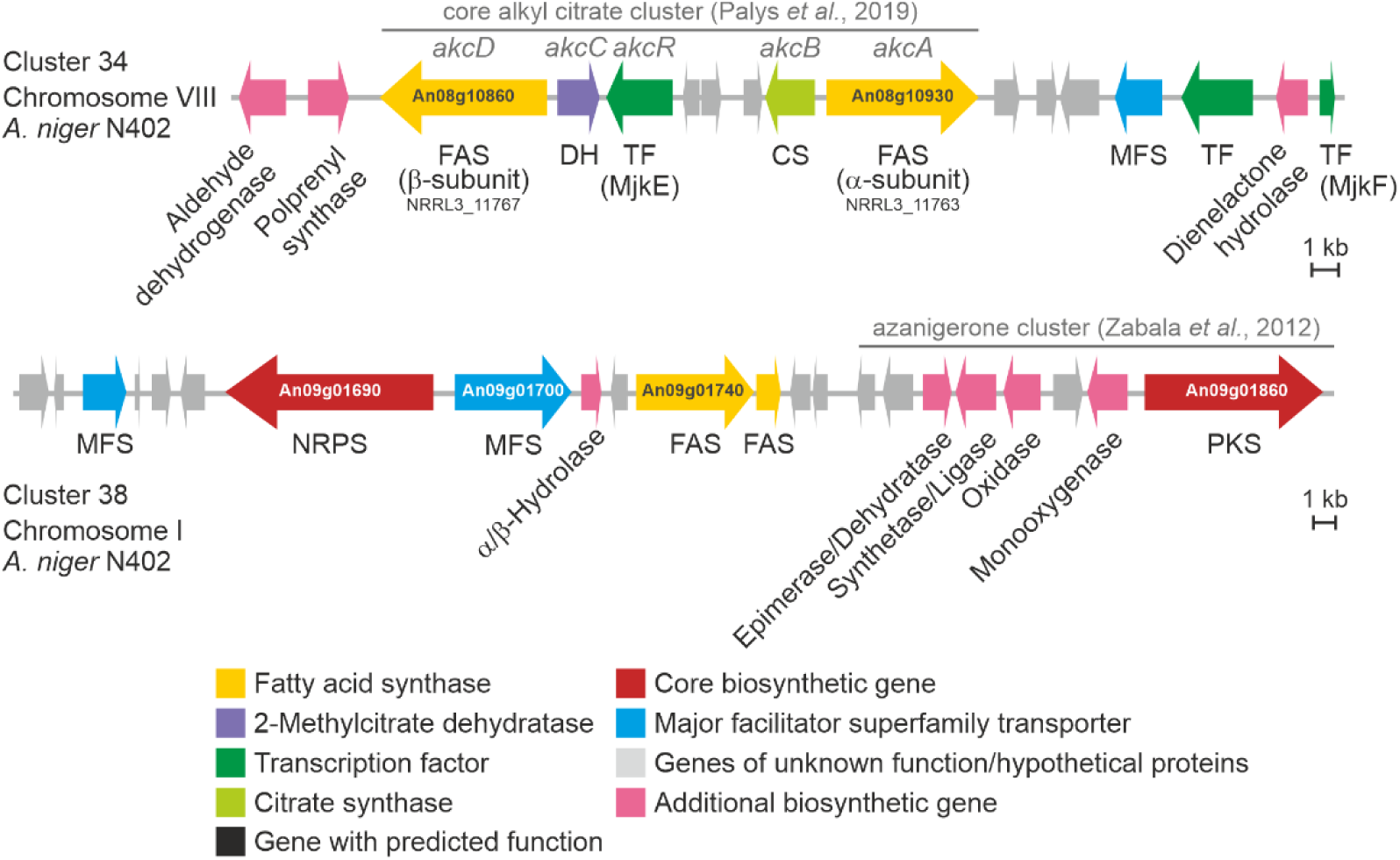
Schematic representation of BGC 34 and BGC 38 as predicted by antiSMASH. Based on sequence similarity and gene functional prediction, BGC 34 corresponds to the alkyl citrate-producing cluster identified in parallel to this study in *A. niger* NRRL3^38^. BGC 38 is positioned next to the azanigerone cluster.

Interestingly, our analysis demonstrates that the SCC approach primarily carves out co-expression of frequently expressed genes, whereas the strength of the MR-PCC approach is the identification co-expression relationships amongst rarely expressed genes. We thus decided to study the impact of six putative TF-encoding genes on *A. niger* secondary metabolism in more depth. Four were predicted by the SCC approach to be co-expressed with at least 10 BGC core genes and are unclustered (MjkA-MjkD), whereas the remaining two were predicted by the MR-PCC approach to be co-expressed with both BGC 34 and 38 and are clustered with BGC 34 (MjkE, MjkF; **Table 1**, **Figure 3**).

**Table 1:** Selected list of transcription factors analyzed in this study, which are co-expressed with BGCs in *A. niger*.

| Name | ORF code | No. of co-expressed BGC core genes (SCC) | No. of co-expressed BGC core genes (MR-PCC) | Clustered in a BGC | Tet-on based overexpression phenotype on solid growth medium |
| --- | --- | --- | --- | --- | --- |
| MjkA | An07g07370 | 14 | - | no | Red pigment formation, reduced growth, sclerotia formation |
| MjkB | An12g07690 | 13 | - | no | Red pigment formation |
| MjkC | An01g14020 | 17 | - | no | Yellow pigment formation, reduced growth |
| MjkD | An07g02880 | 10 | - | no | Yellow pigment formation |
| MjkE | An08g11000 | 13 | 1 | yes (BGC 34) | Brown pigment formation |
| MjkF | An08g10880 | 15 | 1 | yes (BGC 34) | Reduced growth, frequent reversions |

### Overexpression of predicted transcription factors MjkA-F modulate *A. niger* pigmentation and development

Prior to conducting gene functional analysis experiments, we assessed gene expression profiles for *mjkA* - *mjkF* across our 155 cultivation conditions. While both *mjkE* and *mjkF*, which reside in BGC 34, were rarely expressed, the four *mjkA* – *mjkD* genes encoding unclustered TFs were transcribed under numerous conditions, with *mjkA* notably expressed to 90% the level of *A. niger* actin under several conditions (**Figure 4**).

**Figure 4:**
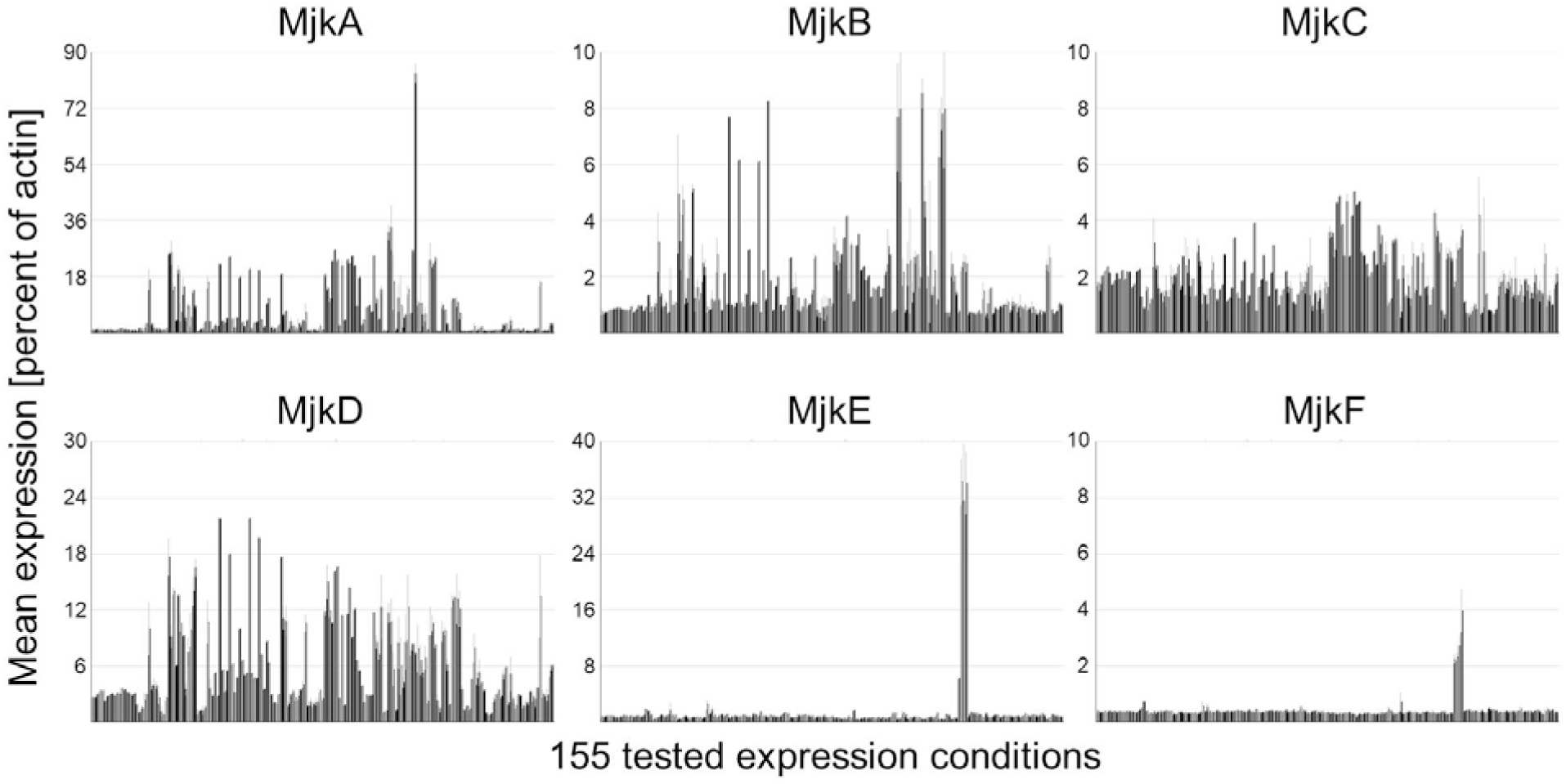
Expression levels for all 6 TFs in 155 expression conditions. Note the different scale bars. MjkE (An08g11000) and MjkF (An08g10880) are only expressed during maltose-limited bioreactor in developmental mutant deleted in the *flbA* gene^41^

To assess the role of these TFs in modulating BGC expression, we generated conditional expression isolates in which a Tet-on gene switch was placed upstream of the open reading frame as previously described for the genes *mjkA* and *mjkB*^33^. This gene switch has undetectable levels of basal expression in the absence of induction, and addition of 10 µg/ml Dox enables expression above that of the *A. niger* glucoamylase gene, whose promoter is often used for overexpression studies33,39,40. Conditional expression isolates previously constructed for genes *mjkA* and *mjkB* were also analyzed in this study to further assess their role in *A. niger* secondary metabolism and development (Supplemental Table 3).

Standard growth assays on solid and in liquid media clearly identified differences in media pigmentation in overexpression isolates when compared to the progenitor control (**Figure 5**, Supplemental Figure 2), suggesting a role of these genes in *A. niger* development and/or secondary metabolism. The conditional expression strains MjkA, MjkD, and MjkF also displayed reduced growth on solid agar under overexpression conditions (Supplemental Figure 3). Intriguingly, *mjkA* overexpression also resulted in the formation of sclerotia (**Figure 5**), which are an important prerequisite for sexual development in Aspergillus^42^. However, *A. niger sensu stricto* has not been reported to have a sexual cycle. Still, *A. niger* rarely produces sclerotia under specific growth conditions, which are paralleled by the production of many secondary metabolites including indolterpenes of the aflavinine type^42^. We thus re-analyzed transcriptomic data which were available for this isolate and for the MjkB overexpression strain from bioreactor cultivation^33^ to screen for differential expression of developmental regulators following conditional MjkA and/or MjkB expression. Strikingly, the expression of 36 and 27 regulators and TFs were affected when *mjkA* or *mjkB* were up- or downregulated, respectively (**Figure 6**). Notably, the overexpression of MjkA resulted in downregulation of genes encoding transcription factors known to control primary metabolism (*creA, areB, xlnR, amyR, prtT, pacC, crzA, hapX, farA, farB, acuB*^43^) and asexual development (*brlA, abaA, stuA, flbA, flbB, flbC*^43^) as well as chromatin structure (*laeA, velB, vipC, mtfA, hdaA*^43^) in *Aspergillus* (**Figure 6**). Deletion of *mjkA* caused strong upregulation of the regulator-encoding genes *areA, cpcA, msnA, csnE, flbD* and *vosA* (Supplemental Table 4) with functions in primary metabolism and development^43^, implying that MjkA is a global regulator of *A. niger* metabolism, differentiation and development and hierarchically placed on a higher level than so far known global regulators in *Aspergillus* mentioned above. Note that the MjkA encoding gene can be found in 61 / 83 sequenced *Aspergillus* genomes as identified by BLAST analyses (Supplemental Table 5).

**Figure 5:**
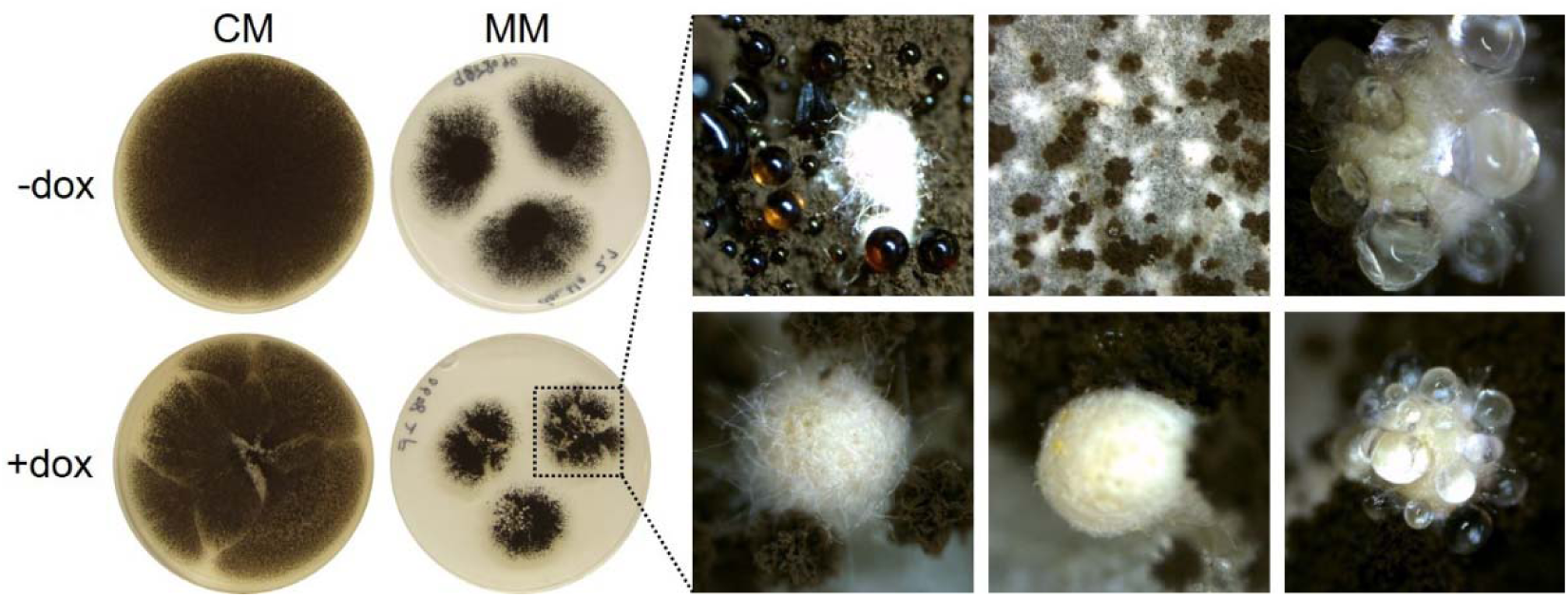
Tet-on-based overexpression of *mjkA* modifies *A. niger* development. Overexpression of *mjkA* induced by the addition of 10 µg/ ml doxycycline leads to sclerotia formation on agar plates, especially when cultivated on minimal medium (MM).

**Figure 6:**
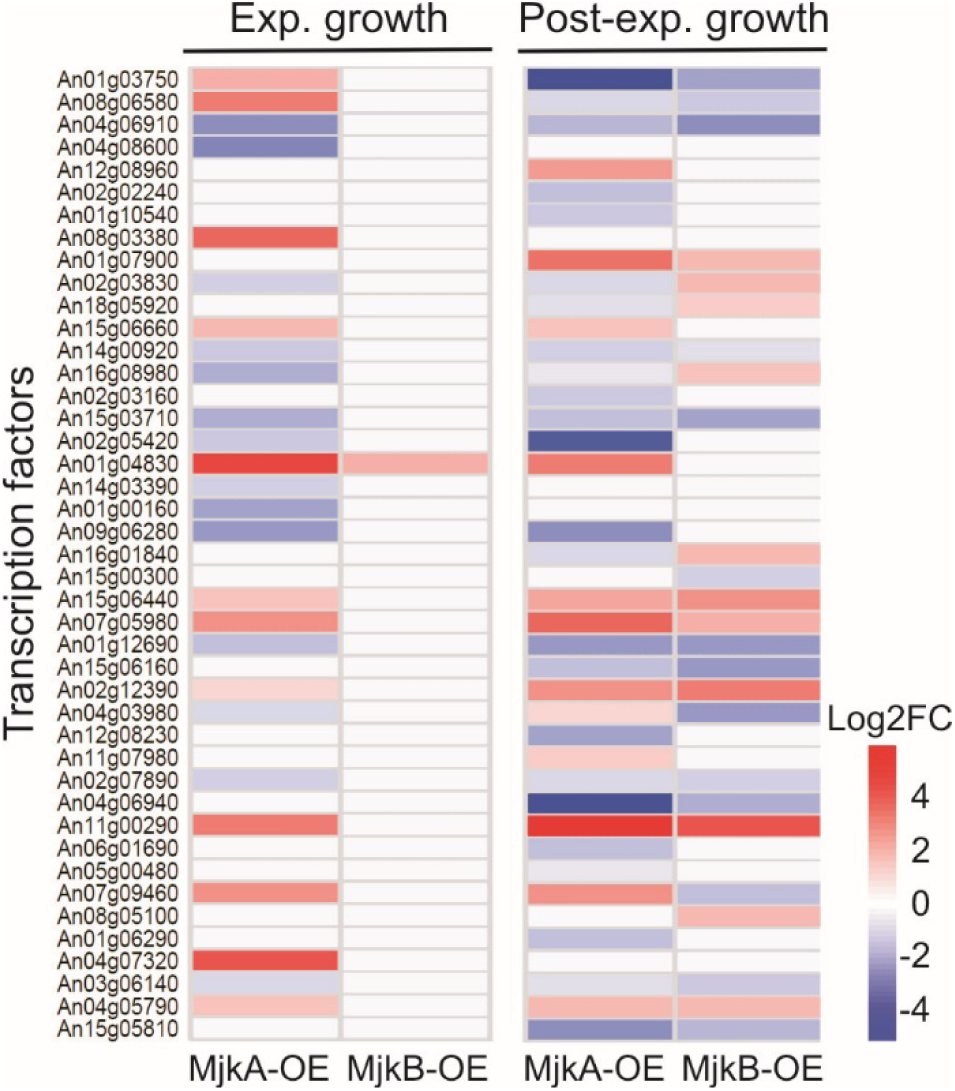
Differential gene expression of transcription factors following overexpression of *mjkA* and *mjkB* genes during controlled bioreactor batch cultivations of *A. niger* performed in our previous study^33^. Note that overexpression of MjkA strongly affects expression of predicted regulators during both growth phases, whereas the effect of MjkB is limited to the post-exponential growth phase. ORF codes are given.

### Overexpression of predicted transcription factors MjkA-F modulates the secondary metabolite profile of *A. niger*

To understand the effect of the MjkA-F TFs on the secondary metabolite profile of *A. niger*, we next conducted untargeted metabolome analysis of the progenitor strain and *mjkA*-*mjkF* conditional expression strains after 2, 4 or 10 days of incubation on minimal agar plates supplemented with 10 µg/ml Dox. For each overexpression strain, one single time point was selected for metabolome analysis. Time points were chosen when the greatest deviation in either media pigmentation or growth relative to the control strain was observed (Supplemental Figure 3). Since culture samples were harvested at the center as well as the outer edges of the growing colonies and pooled for analysis, the obtained results comprise metabolites from both old and young mycelia. This analysis detected a total of 2,063 compounds, from which 1,835 were annotated. Metabolic pathway visualization of the identified metabolites using iPATH showed that intermediates from various biosynthetic routes towards SMs (Supplemental Figures 4 and 5) were covered. Statistical analysis (*t*-test) identified numerous metabolites that were significantly different (*p* ≤ 0.05 and log_2_ ratio > 1 or -1) for the compared genotypes and time points (**Figure 7**A).

**Figure 7:**
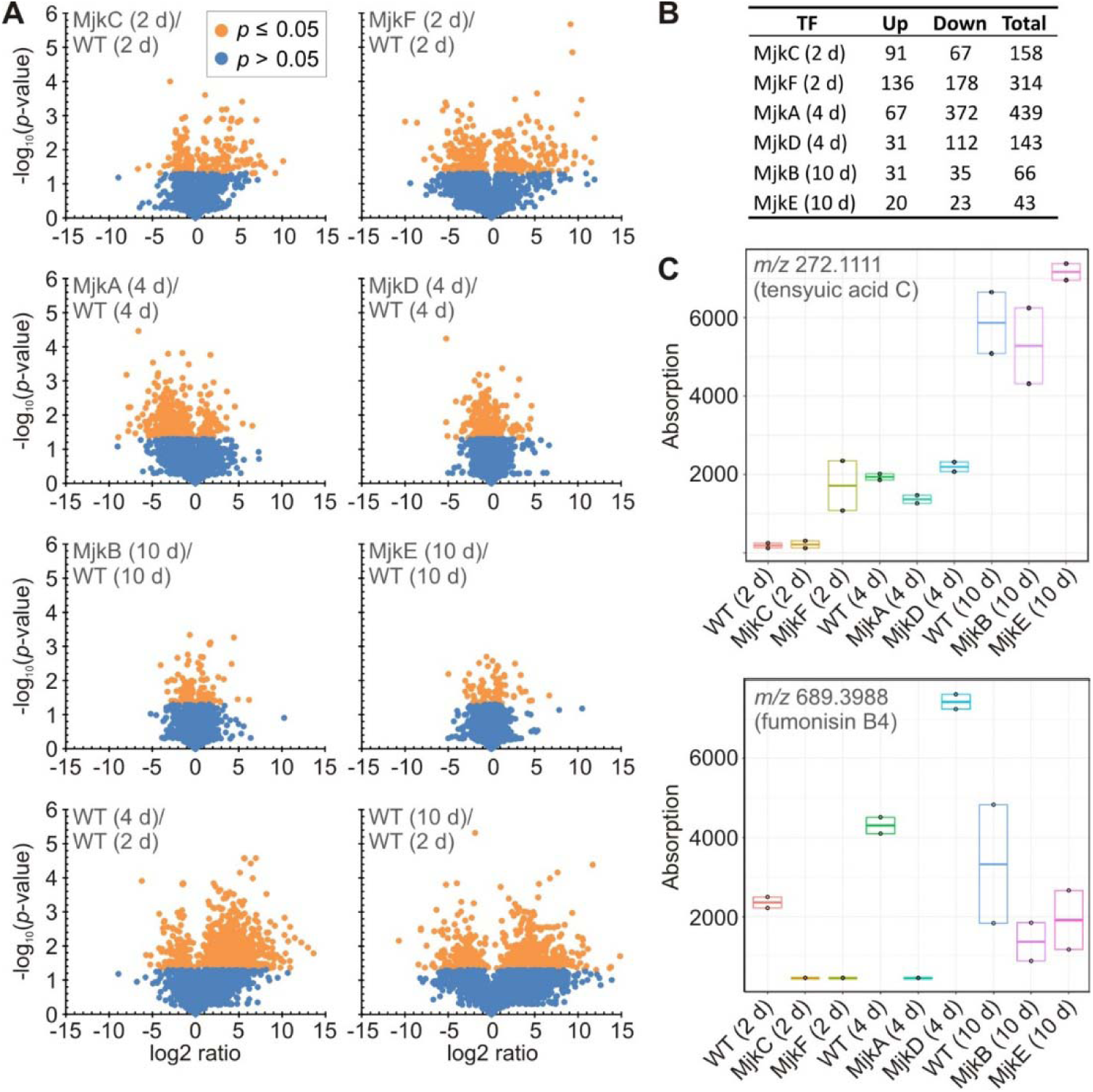
Overexpression of *mjkA* - *mjkF* genes affects numerous metabolites in *A. niger*. (A) Annotated metabolites were plotted by significance (*p*-value) versus fold-change (log2 ratio). Metabolites reaching a *p*-value < 0.05 are marked orange. Metabolites with a *p*-value < 0.05 and a log_2_ ratio > 1 or -1 were considered significant. (B) Number of significantly affected metabolites (*p*-value < 0.05 and log_2_ ratio > 1 or -1) in comparison to the control strain. (C) Exemplary visualization of tensyuic acid C (alkyl citrate) and fumonisin B4 abundances during cultivation of overexpression and control strains of *A. niger* on agar plates at different time points (biological duplicates).

Generally, overexpression of *mjkC* and *mjkF* (2 days) as well as overexpression of *mjkA* and *mjkD* (4 days) each affected more than 140 metabolites (**Figure 7**B). Interestingly, only overexpression of *mjkC* led to an upregulation of more than half of the affected metabolites, whereas overexpression of *mjkA, mjkD* and *mjkF* led to down-regulation (**Figure 7**B). In comparison, overexpression of *mjkB* and *mjkE* (10 days) apparently affected fewer metabolites (66 and 43, respectively), which might also be due to a reduced overall metabolic activity of the cultures after prolonged cultivation.

Amongst the significantly affected metabolites, several known SMs of *A. niger* and related species^44^ could be putatively identified by means of LC-QTOF-HRMS based on mass and retention time (**Figure 7**C, Supplemental Figures 6 and 7). These compounds comprise naphto-γ-pyrones (aurasperones, isonigerone, fonsecin, carbonarins), bicoumarins (bicoumanigrin, kotanin, desmethylkotanin, funalenone), and fumonisins. Moreover, overexpression of the putative TFs affected meroterpenoids (1-hydroxyyanuthone A) and benzoquinone-type pigments (atromentin, cycloleucomelone), as well as different types of alkaloids such as pyranonigrins, pyrophens (aspernigrin A, carbonarone A, nygerone A), nigragillins (nigragillin, nigerazine B), and tensidols. Not found amongst the significantly affected compounds were some known SMs of *A. niger*, which have already been linked to their corresponding BGCs, such as azanigerone^37^, TAN-1612^45^, and ochratoxin^46^.

Notably, the list of previously identified SMs of *A. niger* almost exclusively comprised of polyketide products (Supplemental Figure 6). Thus, even though the peptide-forming NRPS from BGC 38 (An09g01690) is present in a mutual rank metamodule with MjkE and MjkF, the biosynthetic product of BGC 38 is unlikely to be one of the compounds identified in the current study. Based on an *in silico* assembly line prediction using antiSMASH, An09g01690 encodes a bimodular NRPS, which cannot be classified yet into a linear or iterative assembly type and its product is thus not predictable. Since it is co-expressed with two putative fatty acid synthase encoding genes (An09g01740, An09g01750) in BGC 38 (**Figure 1** and **Figure 3**), the encoded peptide presumably features a fatty acid moiety of varying length based on the available fatty acid pool of *A. niger*. Similar patterns have been observed for other nonribosomally synthesized lipopeptides such as daptomycin^47^.

In parallel to this study, BGC 34 (**Figure 3**) was recently demonstrated to be responsible for alkyl citrate production in *A. niger* NRRL3^38^. For this SM class, a range of bioactivities has been reported, including antiparasitic^48^, antifungal^49^ antibacterial^50^, and plant root growth promotion effects^51^. Other complex alkyl citrates (zaragozic acids, also called squalestatins) have been shown to be amongst the most potent natural squalene synthase inhibitors^52,53^. Notably, the metabolome analysis in this study showed that several alkyl citrates, such as hexylaconitic acid A, hexylitaconic acid J, tensyuic acid C and E, were also differentially produced upon TF overexpression at different time points (**Figure 7**C, Supplemental Figure 7).

## Discussion

This study has demonstrated that gene co-expression analysis enables the identification of fungal transcriptional networks in which secondary metabolite genes are embedded. By comparing mutual rank and Spearman derived co-expression networks, we have respectively identified both BGC resident and, additionally, unclustered TFs, a finding broadly consistent with the existence of SM regulatory genes that reside outside predicted BGC loci^17^. However, there is a growing body of evidence to suggest that, at least in some instances, there has been an over-reliance on physical clustering for the prediction of SM pathway genes and their cognate transporters/regulators. Indeed, with several notable exceptions^54,55^, it is still relatively rare that genes required for the biosynthesis of an entire fungal SM are firstly experimentally verified, and secondly, fully contiguously clustered. Thus, the true extent of SM pathway gene clustering in fungi remains unclear. This is further complicated by divergence in the degree in which the BGCs are ‘intact’ across fungal genomes, which is even true for ‘gold standard’ BGCs, such as those necessary for epipolythiodioxopiperazine synthesis (e.g., gliotoxin/sirodesmin)^54^. Hence, experimental approaches to activate and functionally analyze the full fungal SM repertoire cannot exclusively rely on *in silico* genomics approaches.

Given that co-expression approaches have only recently been applied to define fungal BGC boundaries and their transcriptional networks^29,31,33,56^, in this study we examined the potential utility of two different approaches for constructing co-expression networks, namely mutual rank and Spearman approaches. Our results suggest that both approaches enable delineation and refinement of contiguous BGC boundaries. However, whereas the Spearman approach was better suited for the identification of global TFs, the mutual rank approach was better suited for the identification of pathway-specific TFs. This work should therefore guide future co-expression analyses of other fungal transcriptional datasets based on the requirements of the end user (i.e., global or pathway-specific studies).

Overexpression of six TF-encoding genes (*mjkA-F*) predicted from co-expression networks to be involved in *A. niger* SM regulation enabled the modification of *A. niger* secondary metabolite profiles, which included the production of SMs that were not detected in the progenitor control (Supplemental Figure 7). Thus, wholesale modulation of fungal SMs in standard lab culture is possible using hypotheses derived from both Spearman/mutual rank network approaches. The simplicity of the culture conditions is an attractive aspect of the discovery pipeline in this work, which may be preferable when compared to more complex experimental setups, such as co-cultivation experiments, or isolation of novel metabolites from the complex fungal niche (e.g., soil) or marine environments^57^.

From a methodological perspective, our data support the notion that TF overexpression using an inducible gene switch is an effective strategy for SM activation, and probably is preferable to conventional gene deletion approaches^33^. It should be noted, however, that this study was clearly not able to activate all *A. niger* SMs, as we only analyzed SM profiles from a single growth stage/time point for each mutant. Therefore, we speculate that activation of other metabolites will be observed at different culture conditions/growth phases. Consequently, the full exploration of the SM repertoire of *A. niger* isolates MjkA-F will be conducted in follow up studies. Where conservation of MjkA-F is observed in other fungal genomes, the functional analysis (i.e., overexpression) of such orthologues to activate and discover other SM molecules appears feasible.

An exciting observation during this study was that MjkA seemed to function at a hierarchy above major transcriptional regulators, such as CreA, AreA, PacC, BrlA, CrzA, and LeaA, to name but a few (**Figure 6**). Additionally, the formation of sclerotia due to overexpression of MjkA can be viewed as a preliminary (and tentative) step towards laboratory-controlled sex, opening up the possibility of classical genetics in this species^58^. Such developmental jackpots may be viewed as an additional benefit to wholesale analysis of fungal SMs using co-expression networks.

In this work, we also conducted significant *in silico* and mass spectrometry-based characterization of differential SM production profiles and attempted to link empirically observed SMs to specific BGCs. Despite recent advances in publicly available tools for such experiments, including the prediction of putative SM structures based on the analysis of PKS/NRPS domains^59^, coupling BGCs to their products is still challenging. In this respect, linking BGCs amongst multiple differentially produced SMs between control and experimental cohorts remains a significant bottleneck in discovery pipelines and requires experimental validation of putative BGC-metabolite candidates, e.g., by means of core gene knockout or overexpression.

In summary, this study has generated novel co-expression resources and methods for the microbial cell factory *A. niger*. Strains MjkA-F are promising tools for metabolite discovery and will be used in future to reverse engineer the transcriptional networks to which they belong. Our data clearly support the well-established prevalence of BGCs in filamentous fungal genomes, but suggest a refinement to this paradigm — whereby for activation and functional analysis experiments of SMs, it may be safer to consider that the necessary genes for a fungal SM of interest (including core genes, tailoring genes, transporters, detoxifiers, and regulators) may be unclustered, but can be identified by means of SCC as well as MR-PCC co-expression analyses. Such shifts in experimental thinking may help facilitate the full exploitation and comprehensive understanding of SMs amongst the fungal kingdom.

## Materials and Methods

### Calculating mutual rank for microarray experiments

*A. niger* microarrays across a range of experimental conditions and genetic backgrounds^33^ were analyzed in R using the affy, simpleaffy, and makecdfenv packages^60–62^. Raw data from each of the 283 individual microarrays were normalized using RMA as implemented in the affy package^60^. To enable cross-experiment comparisons, expression values were normalized by scaling to the cross-experiment trimmed mean (excluding the top and bottom 5% of expression values). Pearson’s correlation coefficient was calculated between every pair of genes across all conditions. An ordered list of all genes from most to least correlated was generated for each gene. For every pair of genes, the mutual rank was calculated by taking the geometric mean of the rank of each gene in the other gene’s ordered list. The mutual rank (MR) of two genes A and B is the geometric mean of each gene’s correlation rank, and is given by the formula:

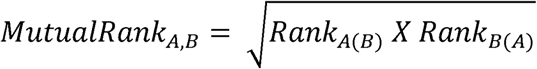

where *Rank*_*A(B)*_ is the rank of gene B in an ordered list of the correlation coefficients of all genes with respect to gene A ranked from most to least correlated^34^. MR scores were transformed to network edge weights using the exponential decay function *e*^-(*MR*-1/*x*)^; three different networks were constructed with *x* set to 5, 10, and 25, respectively. Edges with a Pearson’s correlation coefficient < 0.3 or an edge weight < 0.1 were excluded from the global network, which was then visualized in Cytoscape^63^. Modules of co-expressed genes were inferred using ClusterONE with default parameters^64^. Modules were analyzed for the presence of transcription factors and for SM backbone genes based on protein domains found within these genes and from gene annotations predicted by antiSMASH^65^. For two transcription factors (MjkE and MjkF), results from all co-expression networks were combined by collapsing all modules containing these genes of interest into a meta-module of non-overlapping gene sets. For identification of shared clusters in Aspergillus species (Supplemental Table 6), MultiGeneBlast^66^ was used with 83 available representative genome assemblies available on NCBI Assembly as search database.

### Strains and molecular techniques

*A. niger* strains used in this study are summarized in Supplemental Table 3. Media compositions, transformation of *A. niger*, strain purification and fungal chromosomal DNA isolation were as previously described^67^. Standard PCR and cloning procedures were used for the generation of all constructs^68^ and all cloned fragments were confirmed by DNA sequencing. Correct integrations of constructs in *A. niger* were verified by Southern analysis^68^. For overexpressing *mjkC, mjkD, MjkE* and *mjkF*, the respective open reading frames were cloned into the Tet-on vector pVG2.2^39^ and the resulting plasmids integrated as single or multiple copies at the *pyrG* locus of strain MA169.4. Details on cloning protocols, primers used and Southern blot results are available upon request from the authors.

### Growth assays

Strains were grown at 30°C in minimal medium (MM) or complete medium (CM), consisting of MM supplemented with 1% yeast extract and 0.5% casamino acids as described previously^69^. When indicated, solid or liquid media were supplemented with doxycycline (DOX) to a final concentration of 10 µg/ml. For the growth assay on plates, 10^5^ spores were inoculated on CM or MM +/- DOX and grown for up to 6 days. For shake flask cultivations, freshly harvested spores were inoculated into 50 ml of MM (10^6^/ml) and grown at 30°C, 200 rpm. DOX was added after 16 hr of inoculation (∼exponential phase) and afterwards every 24 hr until 92 hr. Strain MJK17.25 served as control strain.

### Metabolome profiling

Metabolites were extracted from colonies of *A. niger* MJK17.25 grown on agar plates (independent biological duplicates) by METABOLON (Potsdam, Germany). In brief, three agar plugs (outer edge to plate, centre of colony, outer edge adjacent to next colony) were collected at different time points from a colony cultivated for 2 – 10 days on minimal agar medium and pooled in one reaction tube. Each sample was extracted in a concentration of 0.5 g/ml with isopropanol:ethyl acetate (1:3, v/v) by ultrasound for 60 min and centrifuged at 4°C at 13,500 rpm for 20 min. The supernatant was sterile filtrated (Carl Roth, 0.22µm) and transferred in a new eppendorf tube. All subsequent steps were carried out at METABOLON (Potsdam, Germany). Metabolites were identified in comparison to METABOLON’s database entries of authentic standards. The LC separation was performed using hydrophilic interaction chromatography with a iHILIC®-Fusion, 150×2.1 mm, 5µm, 200 Å column (HILICON, Umeå Sweden), operated by an Agilent 1290 UPLC system (Agilent, Santa Clara, USA).

The LC mobile phase was A) 10 mM Ammonium acetate (Sigma-Aldrich, USA) in water (Thermo, USA) with 95% acetonitrile (Thermo, USA; pH 6) and B) acetonitrile with 5% 10 mM Ammonium acetate in 95% water. The LC mobile phase was a linear gradient from 95% to 65% acetonitrile over 8.5 min, followed by linear gradient from 65% to 5% acetonitrile over 1 min, 2.5 min wash with 5% and 3 min re-equilibration with 95% acetonitrile (flow rate 400 μl/min). Mass spectrometry was performed using a high-resolution 6540 QTOF/MS Detector (Agilent, Santa Clara, USA). Spectra were recorded in a mass range from 50 *m/z* to 1700 *m/z* in positive and negative ionization mode. The measured metabolite concentration was normalized to the internal standard. Significant concentration changes of metabolites in different samples were analyzed by appropriate statistical test procedures (Students test, Welch test, Mann-Whitney test). A *p*-value < 0.05 was considered as significant.

## Acknowledgements

We would like to thank the European Commission for funding (Marie Curie International Training Network QuantFung, FP7-People-2013-ITN, grant no. 607332) and the National Science Foundation (http://www.nsf.gov) for funding grants IOS-1401682 and DEB-1831493 to J.H.W. This work was conducted in part using computational resources provided by the Advanced Computing Center for Research and Education at Vanderbilt University and Information Technology at Purdue. We acknowledge support by the German Research Foundation and the Open Access Publication Funds of TU Berlin.

