## Supplemental Table and Files for "Beyond the biosynthetic gene cluster paradigm: Genome-wide co-expression networks connect clustered and unclustered transcription factors to secondary metabolic pathways"

Author ORCIDs:

Min Jin Kwon: [https://orcid.org/0000-0002-0171-7092](https://orcid.org/0000-0002-0171-7092" \t "_blank)

Charlotte Steiniger: [https://orcid.org/0000-0001-5329-2391](https://orcid.org/0000-0001-5329-2391" \t "_blank)

Timothy C. Cairns: <https://orcid.org/0000-0001-7106-224X>

Jennifer H. Wisecaver: <https://orcid.org/0000-0001-6843-5906>

Abigail Lind: [https://orcid.org/0000-0002-9579-4178](https://orcid.org/0000-0002-9579-4178" \t "_blank)

Carmen Regner: <https://orcid.org/0000-0002-4083-6916>

Carsten Pohl: [https://orcid.org/0000-0002-3402-9595](https://orcid.org/0000-0002-3402-9595" \t "_blank)

Antonis Rokas: <https://orcid.org/0000-0002-7248-6551>

Vera Meyer: <https://orcid.org/0000-0002-2298-2258>

Author email addresses:

Min Jin Kwon:

Charlotte Steiniger:

Timothy C. Cairns:

Jennifer H. Wisecaver:

Abigail Lind:

Carsten Pohl:

Carmen Regner:

Antonis Rokas:

Vera Meyer:

1. Supplemental Figures

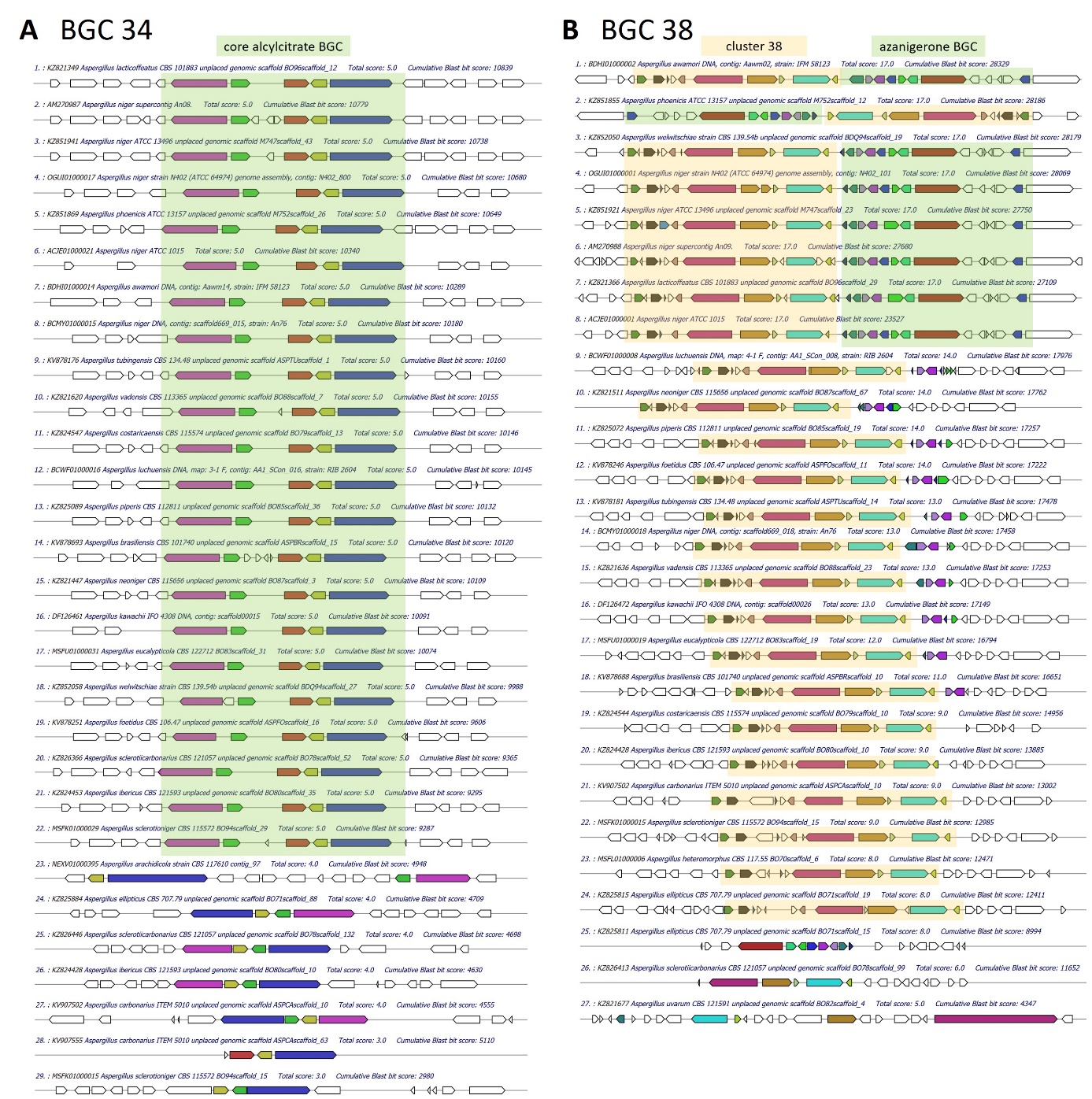

**Supplemental Figure 1:** MultiGeneBLAST of BGC 34 and 38 members reveals several orthologous clusters in related Aspergilli. Cluster members for the query were chosen based on the co-expression heatmap of BGC 34 (An08g10830 - An08g11020) and BGC 38 (An09g01610 - An09g01880), respectively.

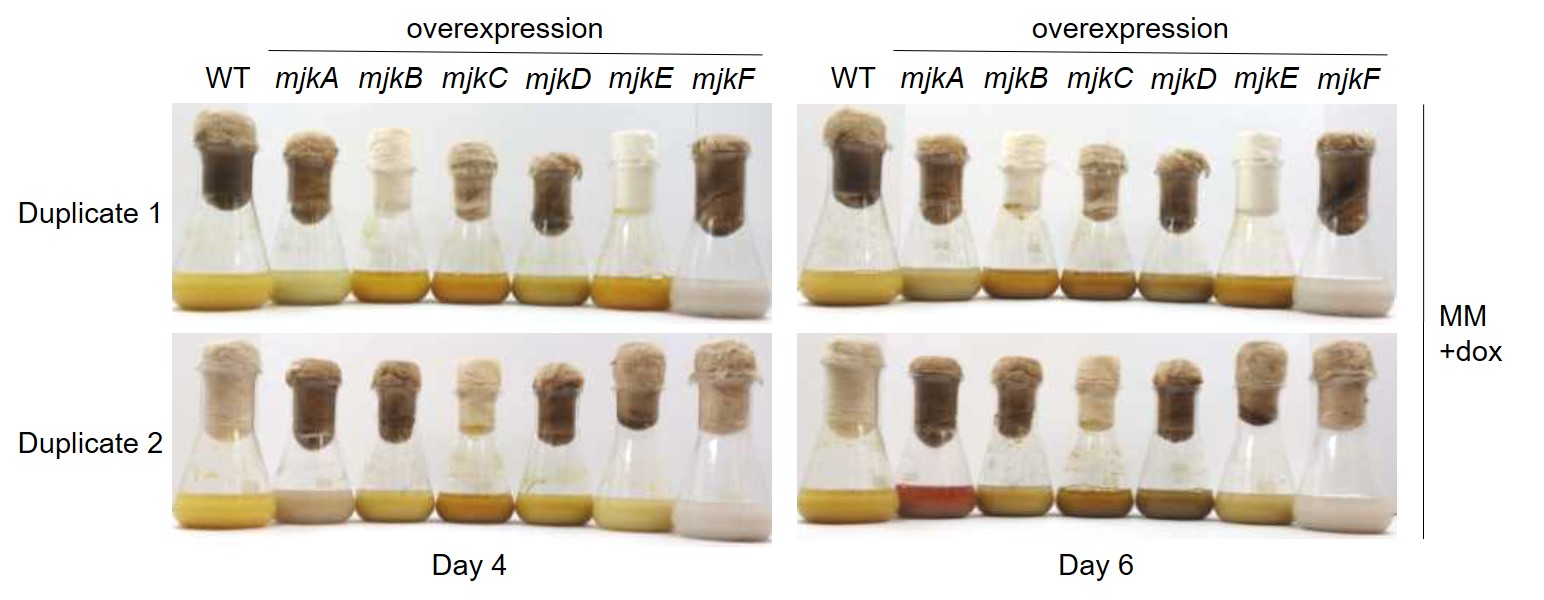

**Supplemental Figure 2:** Phenotypes of A. niger overexpression mutants MjkA – MjkF during shake flask cultivation.

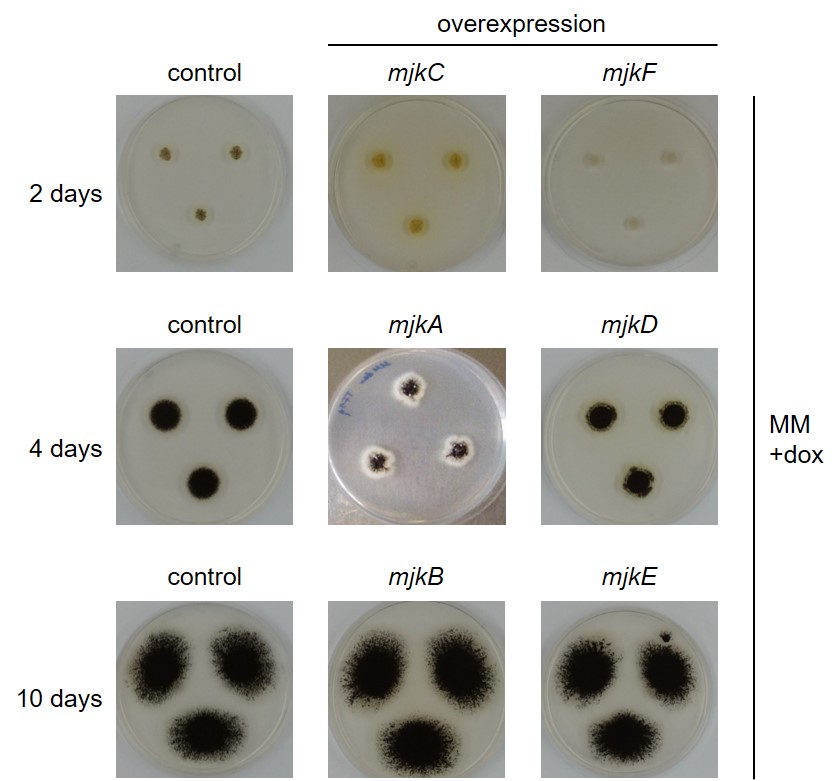

**Supplemental Figure 3:** Phenotypes and sampling time points for metabolome analysis.

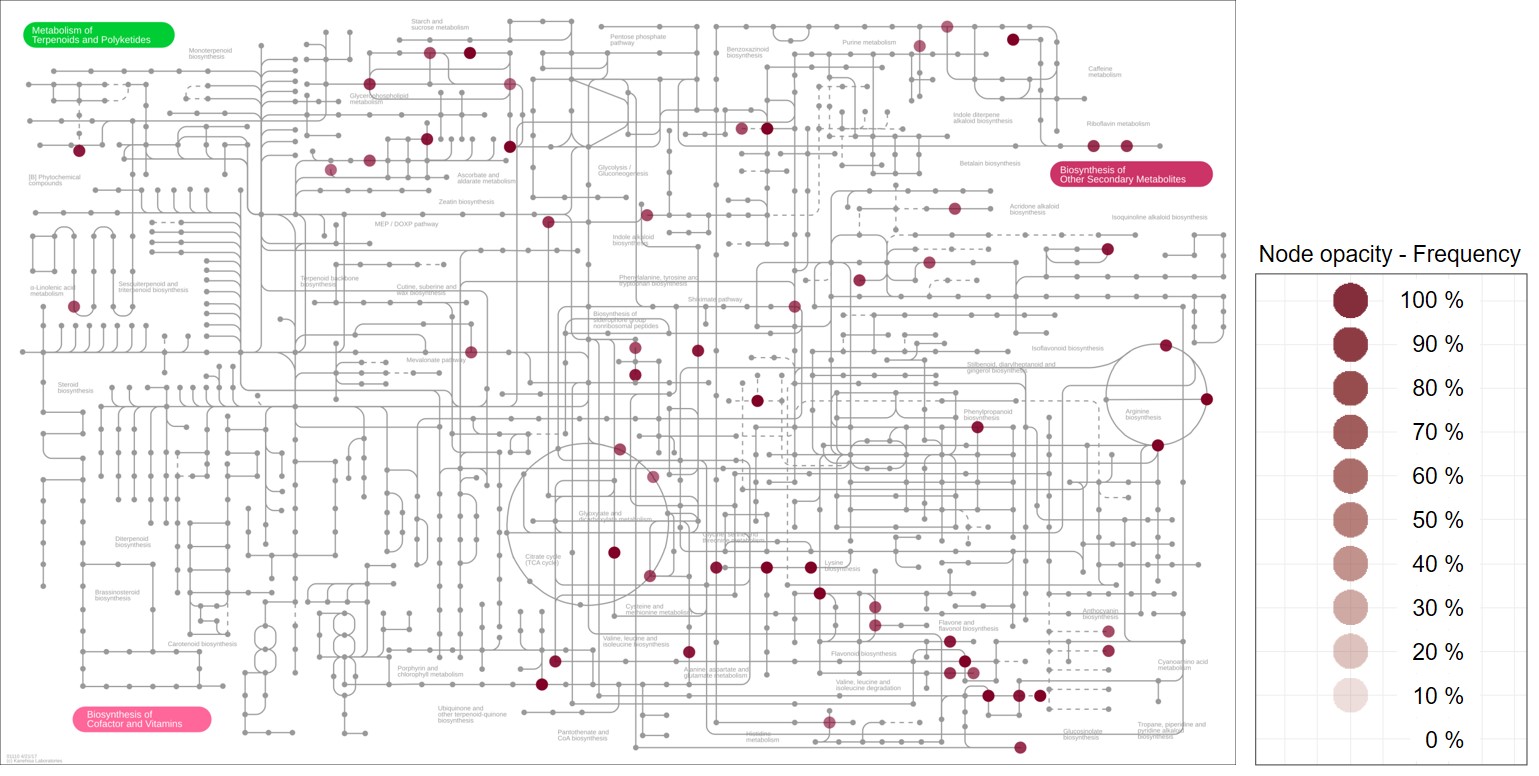

**Supplemental Figure 4:** Identified metabolites involved in SM biosynthesis. Identified metabolites from the untargeted metabolome studies of A. niger were visualized on a metabolic pathway map depicting SM biosynthesis routes using iPATH.

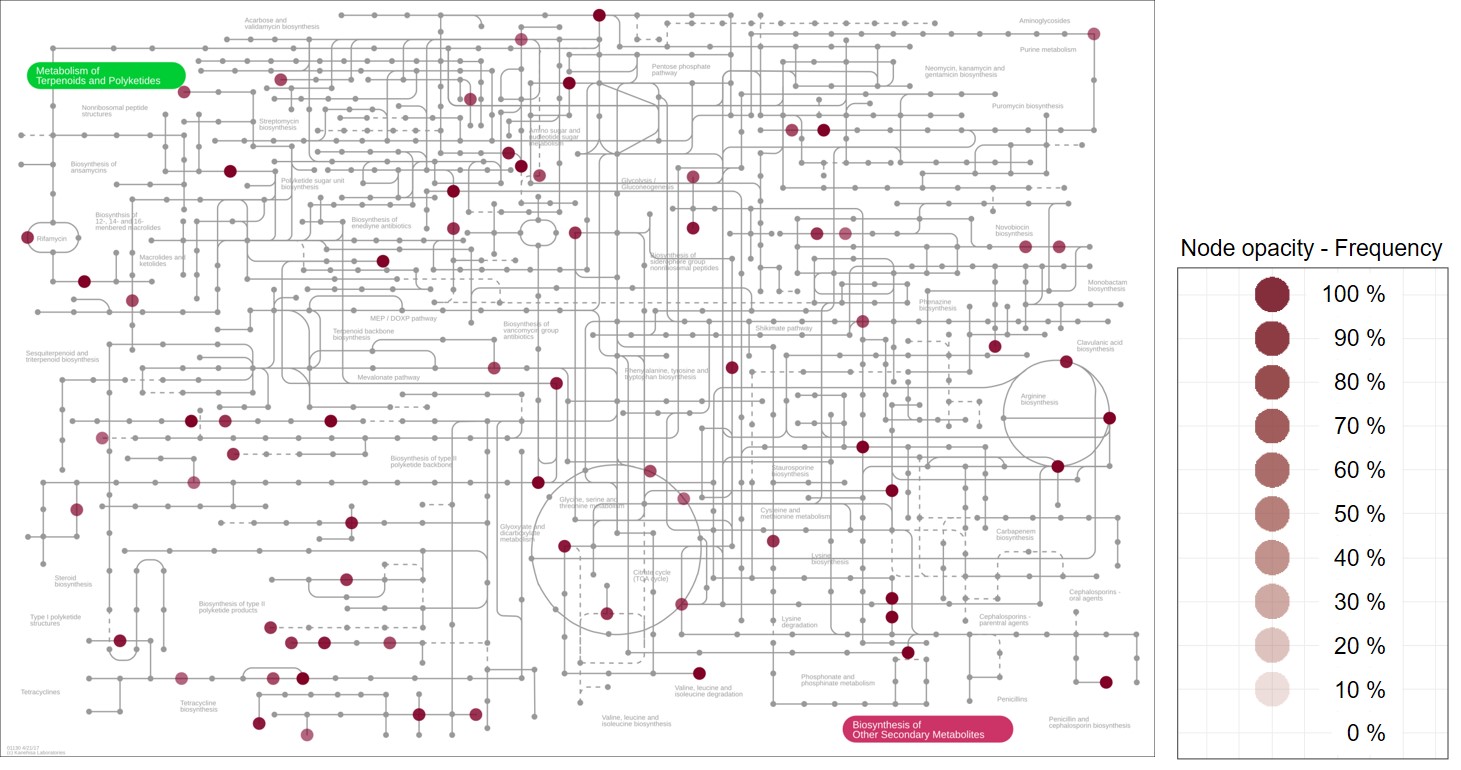

**Supplemental Figure 5:** Identified metabolites involved in antibiotics biosynthesis. Identified metabolites from the untargeted metabolome studies of A. niger were visualized on a metabolic pathway map depicting biosynthesis routes leading to antibiotics using iPATH.

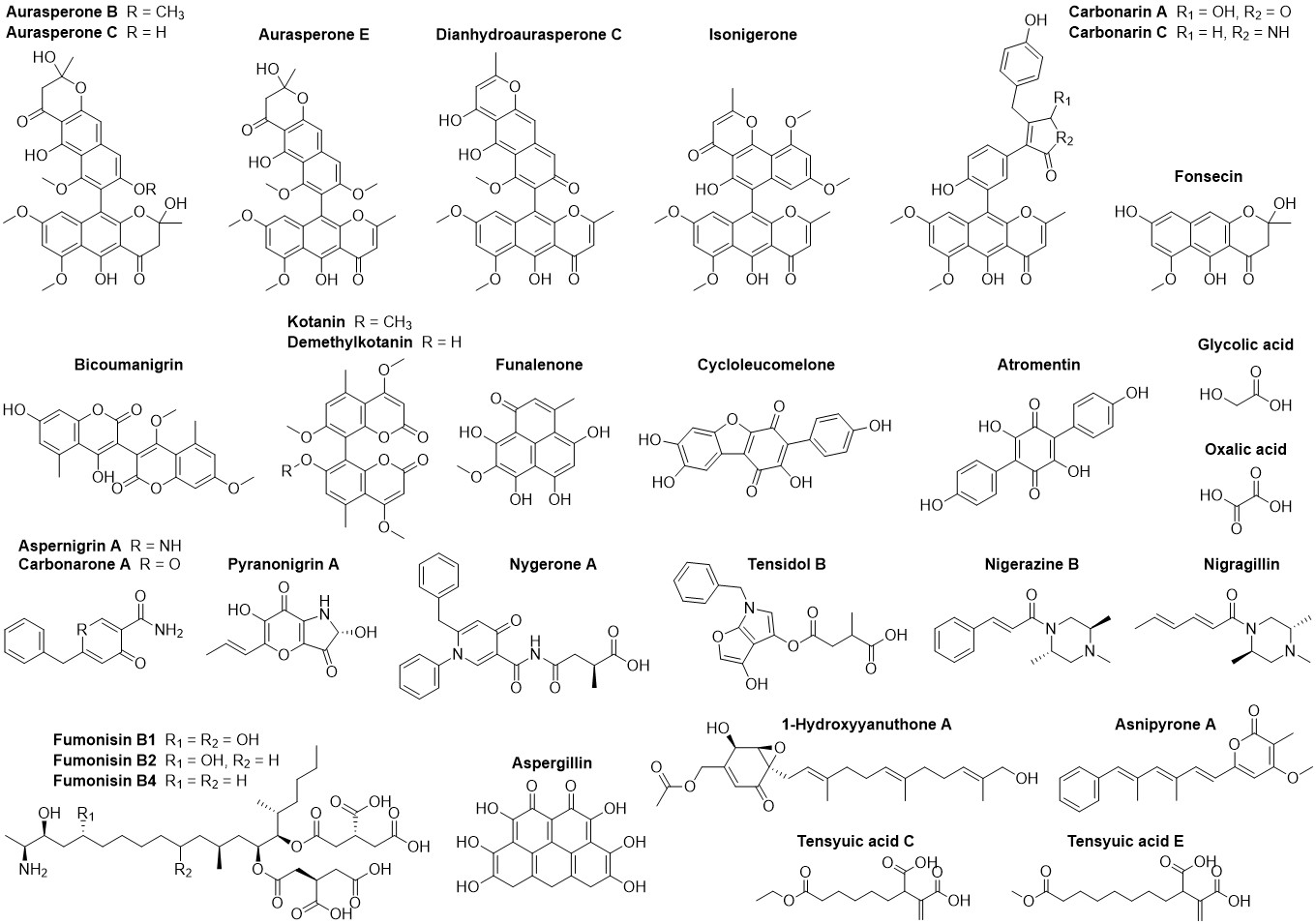

**Supplemental Figure 6:** Known SMs of A. niger and related species affected by overexpression of mjkA - mjkF. Structures of putatively identified SMs from metabolome analysis, which were significantly affected and have been previously described for A. niger and related species (according to Nielsen et al., 2009).

See pdf spreadsheet

**Supplemental Figure 7:** Boxplot visualization of known SMs of A. niger identified from metabolome analysis. SM abundances were monitored during cultivation of overexpression and control strains of A. niger on agar plates at different time points (biological duplicates).

2. Supplemental Tables

**Supplemental Table 1:** All co-expression modules called using ClusterONE.

See excel spreadsheet

**Supplemental Table 2:** List of all co-expression modules containing BGC core genes that overlap with two or more clustered genes.

See excel spreadsheet

**Supplemental Table 3:** Strains used in this study.

| Strain | Relevant genotype/description | Reference |
| --- | --- | --- |
| N402 | *cspA^-^* (derivative of ATCC9029) | Bos *et al.* 1988 |
| AB4.1 | *pyrG^-^* derivative of N402 (*pyrG378*) | Van Hartingsveldt  *et al.* 1987 |
| MA169.4 | Δ*kusA*::DR-*amdS*-DR, *pyrG^-^* (derivative of AB4.1) | Carvalho *et al.* 2010 |
| MJK17.25 | Δ*kusA*::DR-*amdS*-DR, *pyrG^+^* (derivative of  MA169.4 used as control strain) | Schäpe *et al.* 2019 |
| MJK10.12 | Δ*kusA*::DR-*amdS*-DR, Tet-on-*mjkA*-T_trpC_, pyrG+,  multiple copy at *pyrG* locus (derivative of MA169.4) | Schäpe *et al.* 2019 |
| MJK11.17 | Δ*kusA*::DR-*amdS*-DR, Tet-on-*mjkB*-T_trpC_, pyrG+,  single copy at *pyrG* locus (derivative of MA169.4) | Schäpe *et al.* 2019 |
| CR1.5 | Δ*kusA*::DR-*amdS*-DR, Tet-on- *mjkC* -T_trpC_, pyrG+,  single copy at *pyrG* locus (derivative of MA169.4) | This study |
| CR2.2 | Δ*kusA*::DR-*amdS*-DR, Tet-on- *mjkD*-T_trpC_, pyrG+,  single copy at *pyrG* locus (derivative of MA169.4) | This study |
| CR3.12 | Δ*kusA*::DR-*amdS*-DR, Tet-on- *mjkE* -T_trpC_, pyrG+,  single copy at *pyrG* locus (derivative of MA169.4) | This study |
| CR4.15 | Δ*kusA*::DR-*amdS*-DR, Tet-on- *mjkF* -T_trpC_, pyrG+,  single copy at *pyrG* locus (derivative of MA169.4) | This study |

**Supplemental Table 4:** Differential expression of selected genes amongst conditional expression isolates described in this study.

See excel spreadsheet

**Supplemental Table 5:** MjkA homologs amongst Aspergilli identified by BLAST analysis.

See excel spreadsheet

**Supplemental Table 6:** Aspergillus species used for MultiGeneBlast database generation.

| GenBank assembly accession | Organism | Strain |
| --- | --- | --- |
| GCA_000002655.1 | *Aspergillus fumigatus* | Af293 |
| GCA_000002715.1 | *Aspergillus clavatus* | NRRL 1 |
| GCA_000002855.2 | *Aspergillus niger* | CBS 513.88 |
| GCA_000006275.2 | *Aspergillus flavus* | NRRL3357 |
| GCA_000011425.1 | *Aspergillus nidulans* | FGSC A4 |
| GCA_000149205.2 | *Aspergillus nidulans* | FGSC A4 |
| GCA_000149615.1 | *Aspergillus terreus* | NIH2624 |
| GCA_000149645.2 | *Aspergillus fischeri* | NRRL 181 |
| GCA_000150145.1 | *Aspergillus fumigatus* | A1163 |
| GCA_000184455.3 | *Aspergillus oryzae* | RIB40 |
| GCA_000230395.2 | *Aspergillus niger* | ATCC 1015 |
| GCA_000239835.2 | *Aspergillus kawachii* | IFO 4308 |
| GCA_000269785.2 | *Aspergillus oryzae* | 3.042 |
| GCA_000600275.1 | *Aspergillus ruber* | CBS 135680 |
| GCA_000691885.1 | *Aspergillus oryzae* | 100-8 |
| GCA_000731615.1 | *Aspergillus fumigatus* | var. RP-2014 |
| GCA_000812125.1 | *Aspergillus ustus* | 3.3904 |
| GCA_000952835.1 | *Aspergillus flavus* | AF70 |
| GCA_000956085.1 | *Aspergillus parasiticus* | SU-1 |
| GCA_000986645.1 | *Aspergillus rambellii* | SRRC1468 |
| GCA_000986665.1 | *Aspergillus ochraceoroseus* | SRRC1432 |
| GCA_001029325.1 | *Aspergillus fumigatus* | Z5 |
| GCA_001078395.1 | *Aspergillus udagawae* | IFM 46973 |
| GCA_001204775.1 | *Aspergillus nomius* | NRRL 13137 |
| GCA_001445615.1 | *Aspergillus lentulus* | IFM 54703 |
| GCA_001511075.1 | *Aspergillus calidoustus* | HKI Jena |
| GCA_001515345.1 | *Aspergillus niger* | An76 |
| GCA_001602395.1 | *Aspergillus luchuensis* | RIB 2604 |
| GCA_001717485.1 | *Aspergillus cristatus* | GZAAS20.1005 |
| GCA_001792695.1 | *Aspergillus bombycis* | NRRL 26010 |
| GCA_001889945.1 | *Aspergillus brasiliensis* | CBS 101740 |
| GCA_001890125.1 | *Aspergillus versicolor* | CBS 583.65 |
| GCA_001890685.1 | *Aspergillus luchuensis* | CBS 106.47 |
| GCA_001890705.1 | *Aspergillus sydowii* | CBS 593.65 |
| GCA_001890725.1 | *Aspergillus wentii* | DTO 134E9 |
| GCA_001890745.1 | *Aspergillus tubingensis* | CBS 134.48 |
| GCA_001890805.1 | *Aspergillus glaucus* | CBS 516.65 |
| GCA_001890905.1 | *Aspergillus aculeatus* | ATCC16872 |
| GCA_001990825.1 | *Aspergillus carbonarius* | ITEM 5010 |
| GCA_002007945.1 | *Aspergillus oryzae* | BCC7051 |
| GCA_002234955.1 | *Aspergillus fumigatus* | HMR AF 270 |
| GCA_002234965.2 | *Aspergillus turcosus* | HMR AF 23 |
| GCA_002234975.2 | *Aspergillus turcosus* | HMR AF 1038 |
| GCA_002234985.1 | *Aspergillus fumigatus* | HMR AF 706 |
| GCA_002237265.2 | *Aspergillus thermomutatus* | HMR AF 39 |
| GCA_002443195.2 | *Aspergillus flavus* | NRRL 21882 |
| GCA_002443215.2 | *Aspergillus flavus* | NRRL 30797 |
| GCA_002456175.2 | *Aspergillus flavus* | NRRL 118543 |
| GCA_002749805.1 | *Aspergillus arachidicola* | CBS 117610 |
| GCA_002846915.2 | *Aspergillus ochraceoroseus* | IBT 24754 |
| GCA_002847045.1 | *Aspergillus candidus* | CBS 102.13 |
| GCA_002847465.1 | *Aspergillus novofumigatus* | IBT 16806 |
| GCA_002847485.1 | *Aspergillus campestris* | IBT 28561 |
| GCA_002849105.1 | *Aspergillus steynii* | IBT 23096 |
| GCA_002850765.1 | *Aspergillus taichungensis* | IBT 19404 |
| GCA_003184525.1 | *Aspergillus sclerotioniger* | CBS 115572 |
| GCA_003184535.1 | *Aspergillus eucalypticola* | CBS 122712 |
| GCA_003184545.1 | *Aspergillus heteromorphus* | CBS 117.55 |
| GCA_003184585.1 | *Aspergillus saccharolyticus* | JOP 1030-1 |
| GCA_003184595.1 | *Aspergillus lacticoffeatus* | CBS 101883 |
| GCA_003184625.1 | *Aspergillus neoniger* | CBS 115656 |
| GCA_003184635.1 | *Aspergillus sclerotiicarbonarius* | CBS 121057 |
| GCA_003184645.1 | *Aspergillus ellipticus* | CBS 707.79 |
| GCA_003184685.1 | *Aspergillus indologenus* | CBS 114.80 |
| GCA_003184695.1 | *Aspergillus brunneoviolaceus* | CBS 621.78 |
| GCA_003184705.1 | *Aspergillus violaceofuscus* | CBS 115571 |
| GCA_003184745.1 | *Aspergillus uvarum* | CBS 121591 |
| GCA_003184755.1 | *Aspergillus piperis* | CBS 112811 |
| GCA_003184765.1 | *Aspergillus aculeatinus* | CBS 121060 |
| GCA_003184785.1 | *Aspergillus japonicus* | CBS 114.51 |
| GCA_003184825.1 | *Aspergillus fijiensis* | CBS 313.89 |
| GCA_003184835.1 | *Aspergillus costaricaensis* | CBS 115574 |
| GCA_003184845.1 | *Aspergillus ibericus* | CBS 121593 |
| GCA_003184865.1 | *Aspergillus homomorphus* | CBS 101889 |
| GCA_003184925.1 | *Aspergillus vadensis* | CBS 113365 |
| GCA_003344505.1 | *Aspergillus phoenicis* | ATCC 13157 |
| GCA_003344705.1 | *Aspergillus niger* | ATCC 13496 |
| GCA_003344945.1 | *Aspergillus welwitschiae* | CBS139.54b |
| GCA_003369625.1 | *Aspergillus mulundensis* | DSM 5745 |
| GCA_003589665.1 | *Aspergillus sclerotialis* | CBS 366.77 |
| GCA_003709025.1 | *Aspergillus flavus* | CA14 |
| GCA_003850985.1 | *Aspergillus awamori* | IFM 58123 |
| GCA_900248155.1 | *Aspergillus niger* | N402 (ATCC 64974) |
