## Supplemental Figure 7 for "Beyond the biosynthetic gene cluster paradigm: Genome-wide co-expression networks connect clustered and unclustered transcription factors to secondary metabolic pathways"

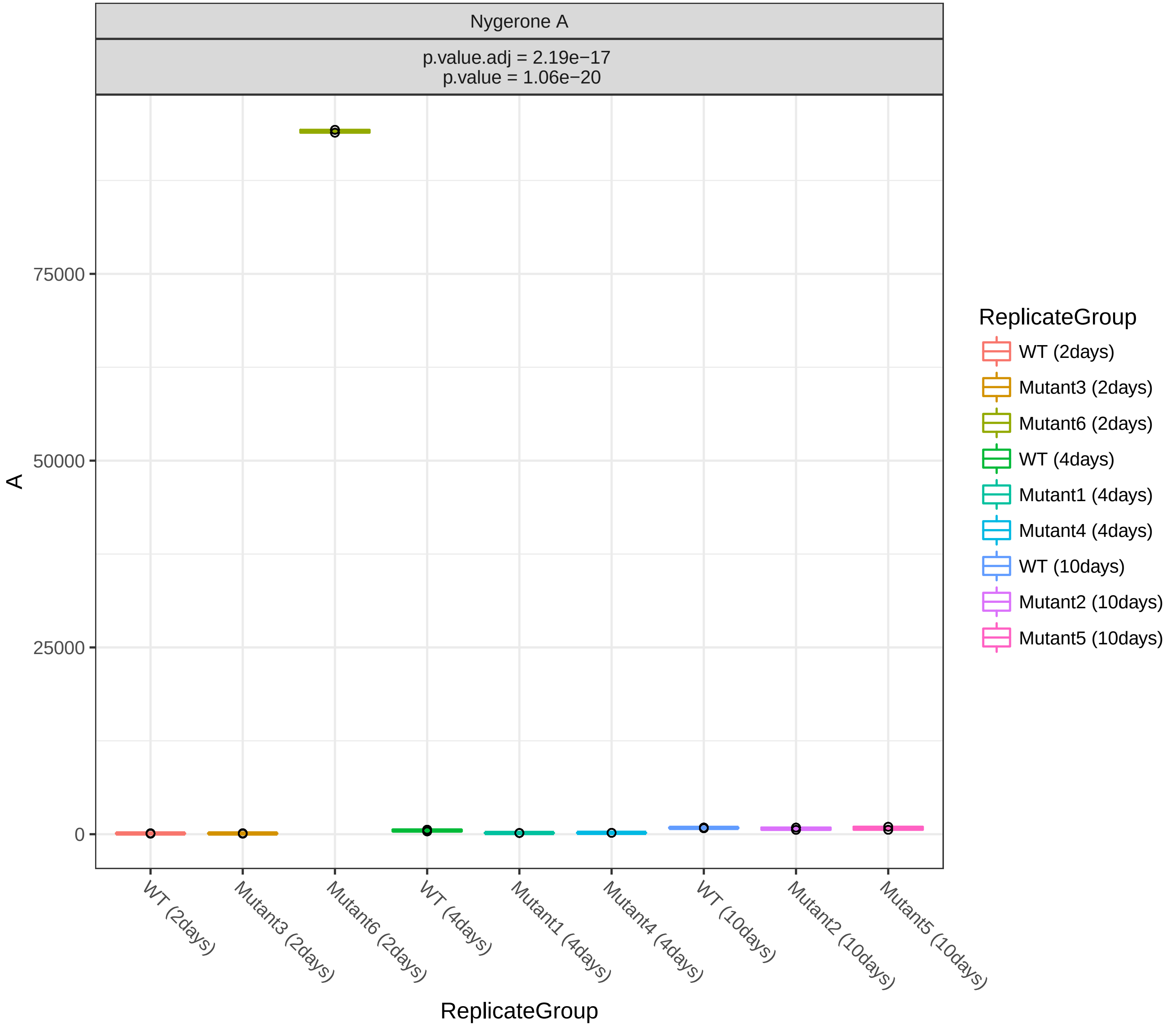
Nygerone A
p.value.adj = 2.19e−17
p.value = 1.06e−20
A
75000
50000
25000
0
WT (2days)
Mutant3 (2days)
Mutant6 (2days)
WT (4days)
Mutant1 (4days)
Mutant4 (4days)
WT (10days)
Mutant2 (10days)
Mutant5 (10days)
ReplicateGroup
ReplicateGroup
WT (2days)
Mutant3 (2days)
Mutant6 (2days)
WT (4days)
Mutant1 (4days)
Mutant4 (4days)
WT (10days)
Mutant2 (10days)
Mutant5 (10days)

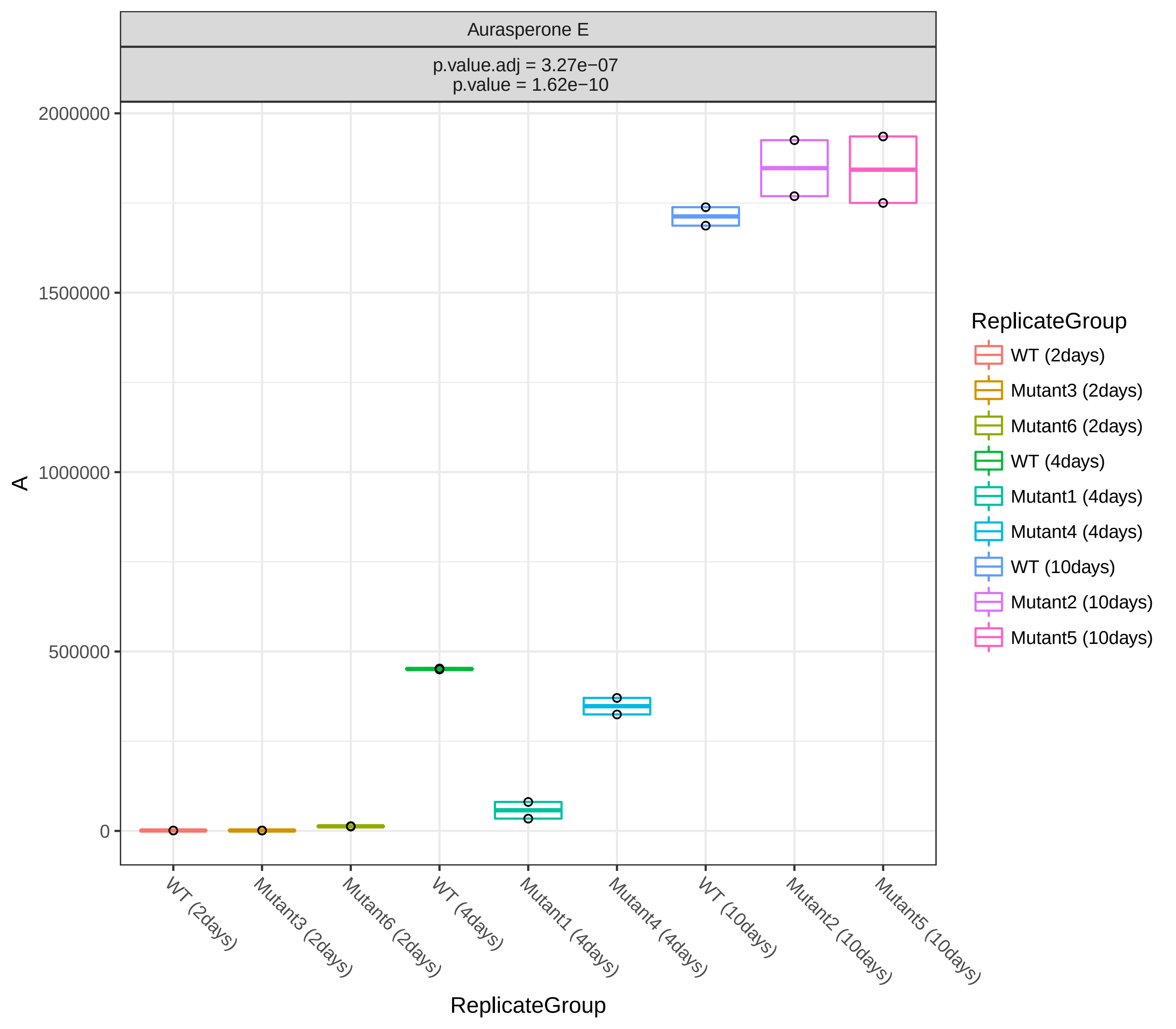
Aurasperone E
p.value.adj = 3.27e−07
p.value = 1.62e−10
2000000
1500000
1000000
500000
0
A
WT (2days)
Mutant3 (2days)
Mutant6 (2days)
WT (4days)
Mutant1 (4days)
Mutant4 (4days)
WT (10days)
Mutant2 (10days)
Mutant5 (10days)
ReplicateGroup
ReplicateGroup
WT (2days)
Mutant3 (2days)
Mutant6 (2days)
WT (4days)
Mutant1 (4days)
Mutant4 (4days)
WT (10days)
Mutant2 (10days)
Mutant5 (10days)

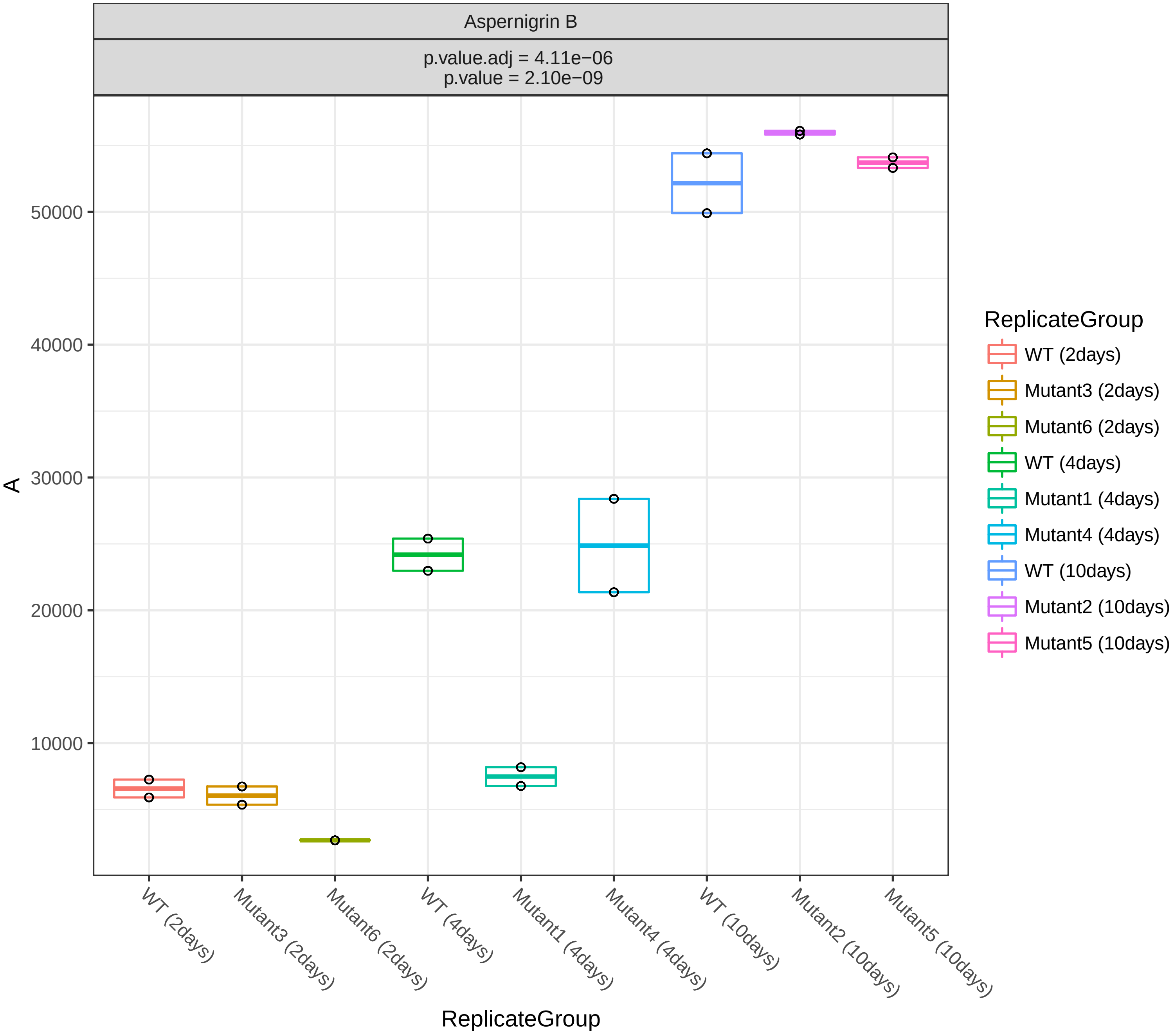
Aspernigrin B
p.value.adj = 4.11e−06
p.value = 2.10e−09
A
50000
40000
30000
20000
10000
WT (2days)
Mutant3 (2days)
Mutant6 (2days)
WT (4days)
Mutant1 (4days)
Mutant4 (4days)
WT (10days)
Mutant2 (10days)
Mutant5 (10days)
ReplicateGroup
ReplicateGroup
WT (2days)
Mutant3 (2days)
Mutant6 (2days)
WT (4days)
Mutant1 (4days)
Mutant4 (4days)
WT (10days)
Mutant2 (10days)
Mutant5 (10days)

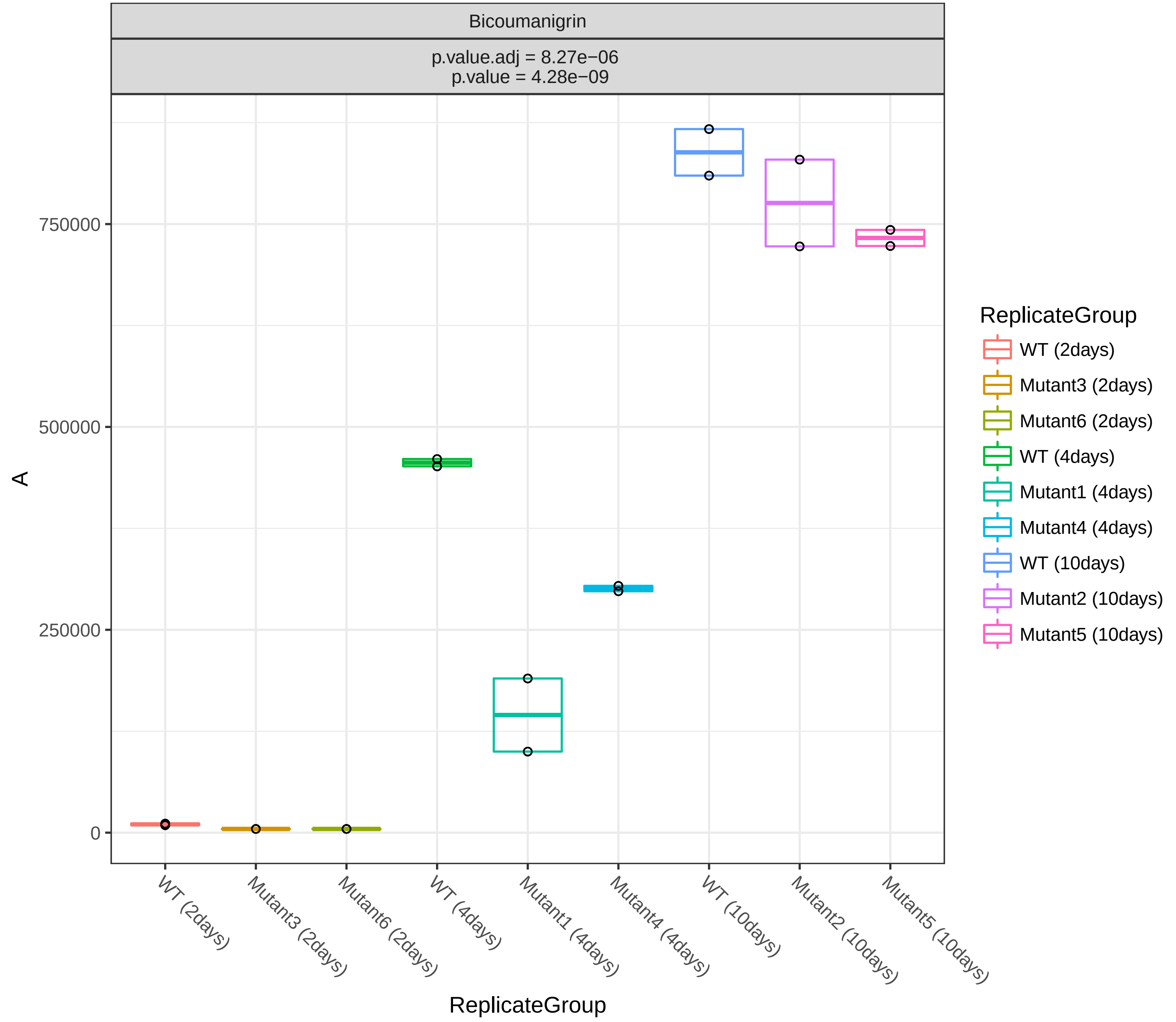

Bicoumanigrin
p.value.adj = 8.27e−06
p.value = 4.28e−09
A
750000
500000
250000
0
ReplicateGroup
WT (2days)
Mutant3 (2days)
Mutant6 (2days)
WT (4days)
Mutant1 (4days)
Mutant4 (4days)
WT (10days)
Mutant2 (10days)
Mutant5 (10days)
ReplicateGroup
WT (2days)
Mutant3 (2days)
Mutant6 (2days)
WT (4days)
Mutant1 (4days)
Mutant4 (4days)
WT (10days)
Mutant2 (10days)
Mutant5 (10days)

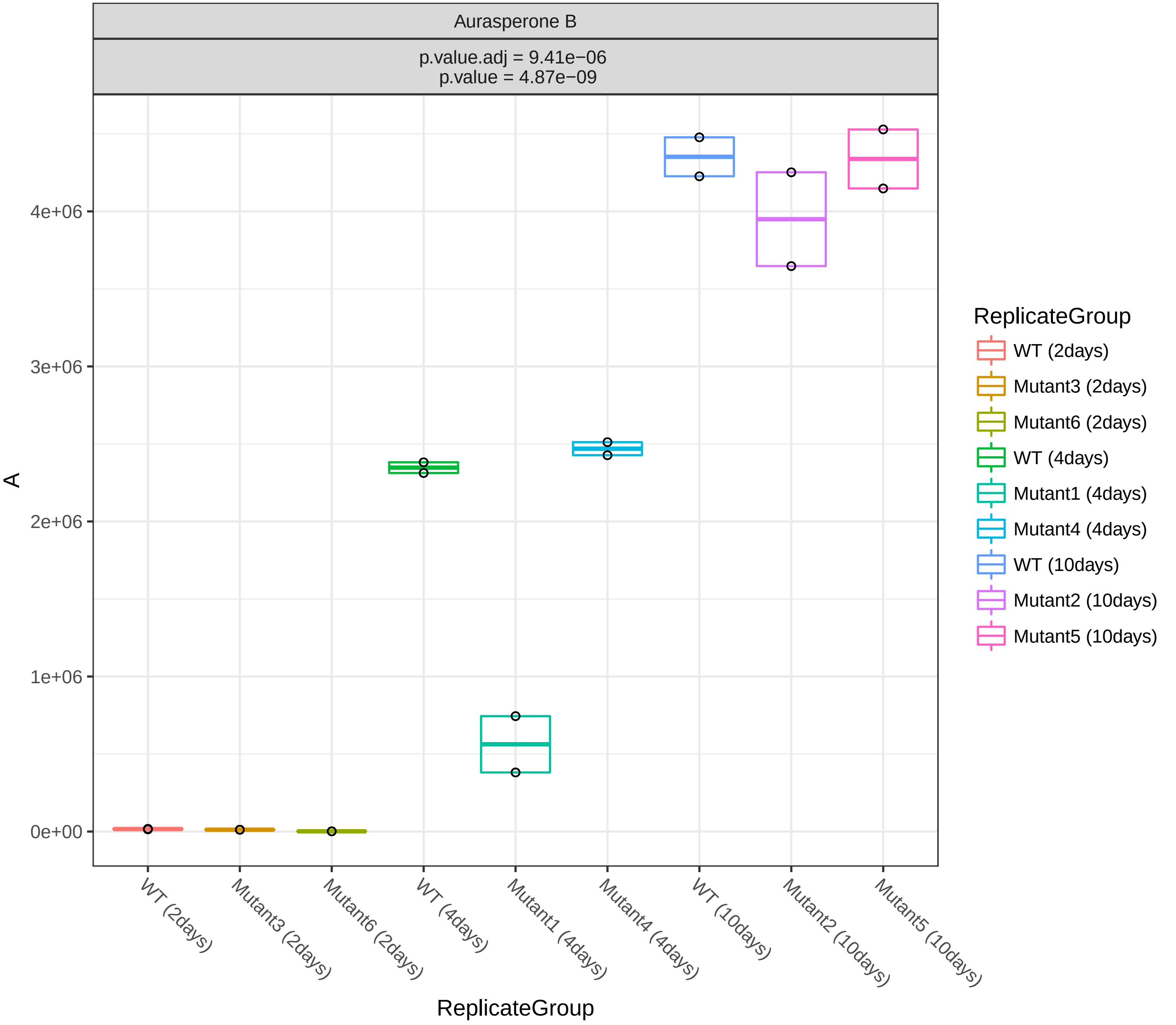
Aurasperone B
p.value.adj = 9.41e−06
p.value = 4.87e−09
A
4e+06
3e+06
2e+06
1e+06
0e+00
WT (2days)
Mutant3 (2days)
Mutant6 (2days)
WT (4days)
Mutant1 (4days)
Mutant4 (4days)
WT (10days)
Mutant2 (10days)
Mutant5 (10days)
ReplicateGroup
ReplicateGroup
WT (2days)
Mutant3 (2days)
Mutant6 (2days)
WT (4days)
Mutant1 (4days)
Mutant4 (4days)
WT (10days)
Mutant2 (10days)
Mutant5 (10days)

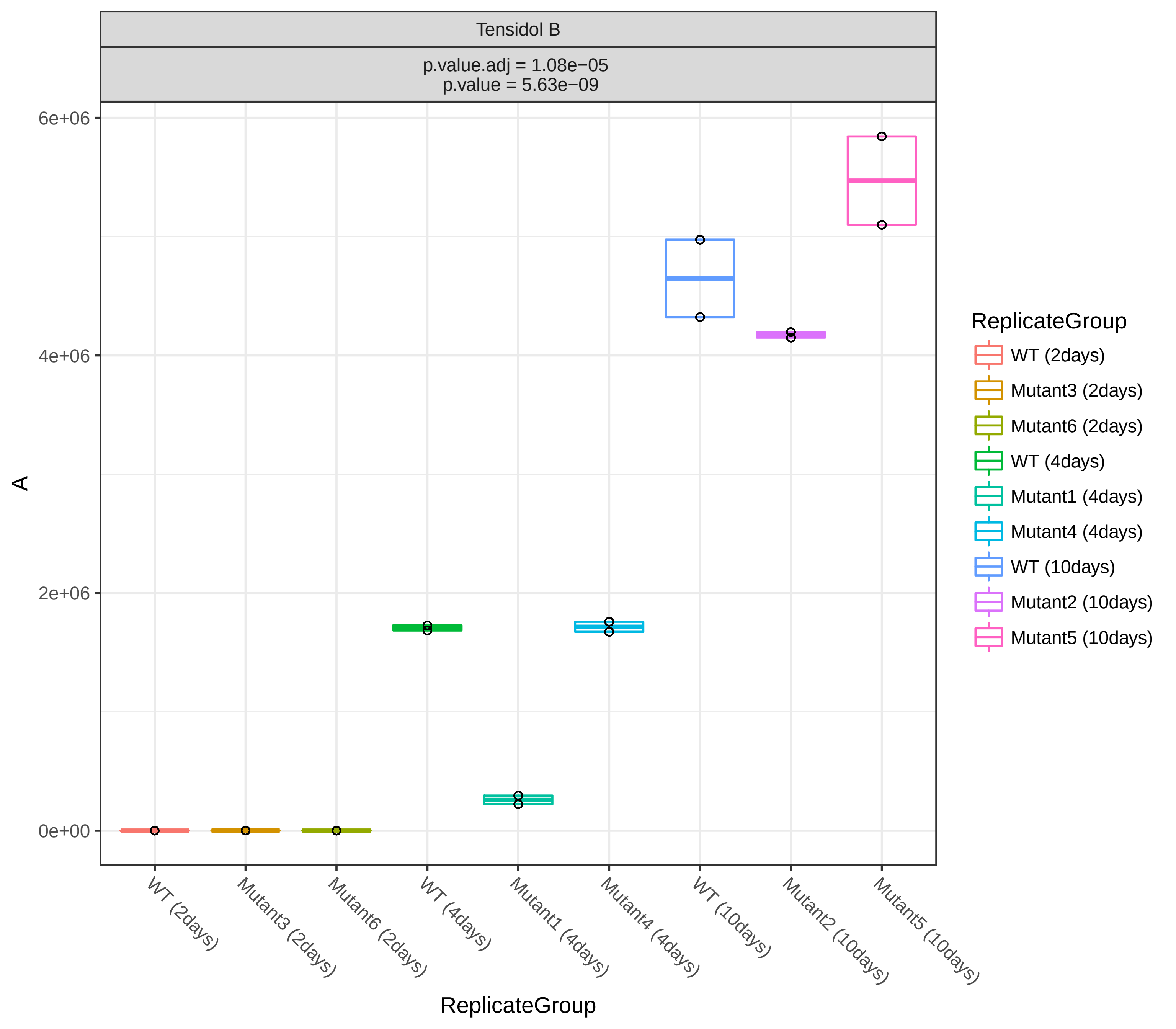

Tensidol B
p.value.adj = 1.08e−05
p.value = 5.63e−09
6e+06
4e+06
2e+06
0e+00
A
ReplicateGroup
WT (2days)
Mutant3 (2days)
Mutant6 (2days)
WT (4days)
Mutant1 (4days)
Mutant4 (4days)
WT (10days)
Mutant2 (10days)
Mutant5 (10days)

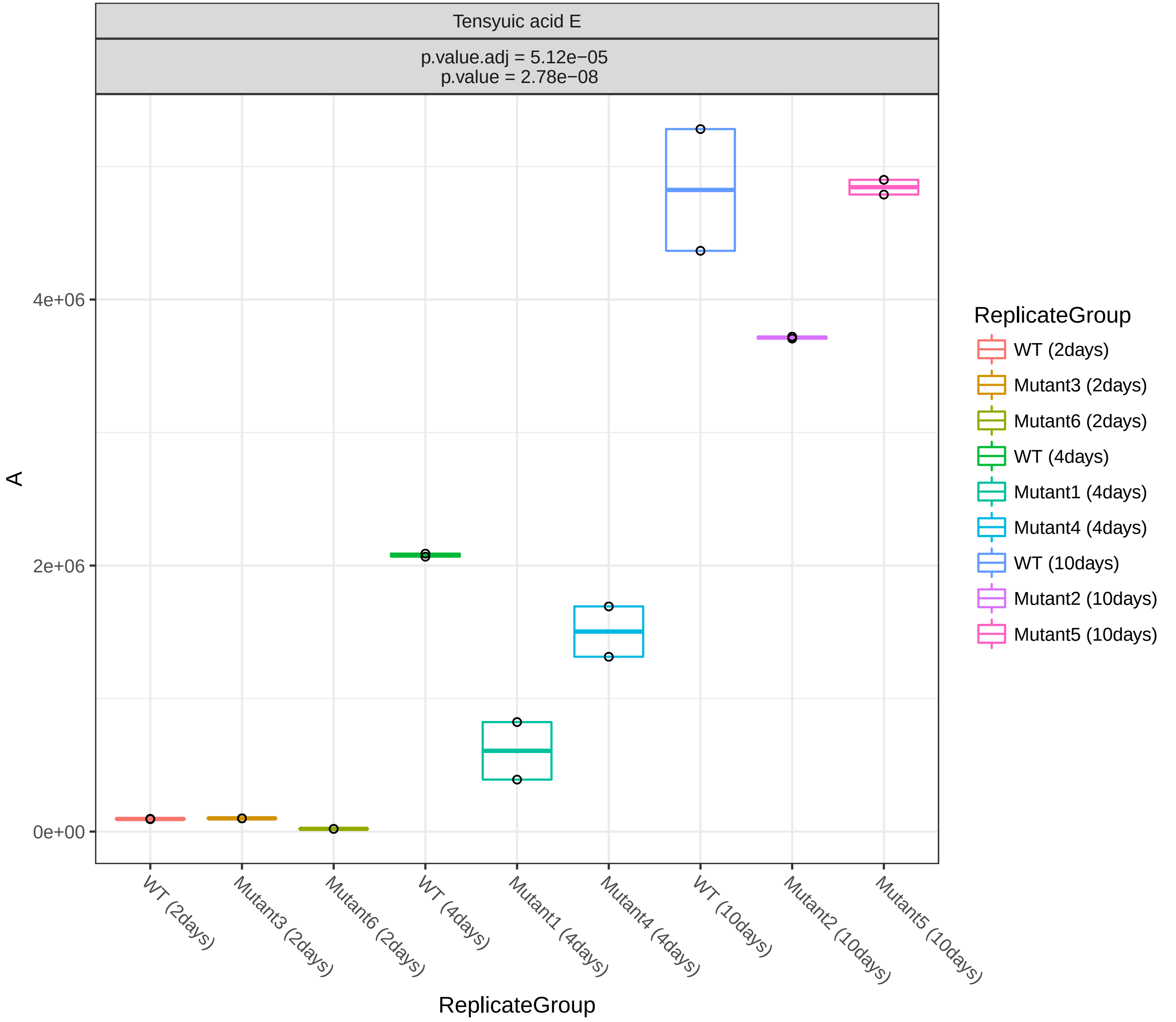

Tensyuic acid E
p.value.adj = 5.12e−05
p.value = 2.78e−08
A
4e+06
2e+06
0e+00
WT (2days)
Mutant3 (2days)
Mutant6 (2days)
WT (4days)
Mutant1 (4days)
Mutant4 (4days)
WT (10days)
Mutant2 (10days)
Mutant5 (10days)
ReplicateGroup
ReplicateGroup
WT (2days)
Mutant3 (2days)
Mutant6 (2days)
WT (4days)
Mutant1 (4days)
Mutant4 (4days)
WT (10days)
Mutant2 (10days)
Mutant5 (10days)

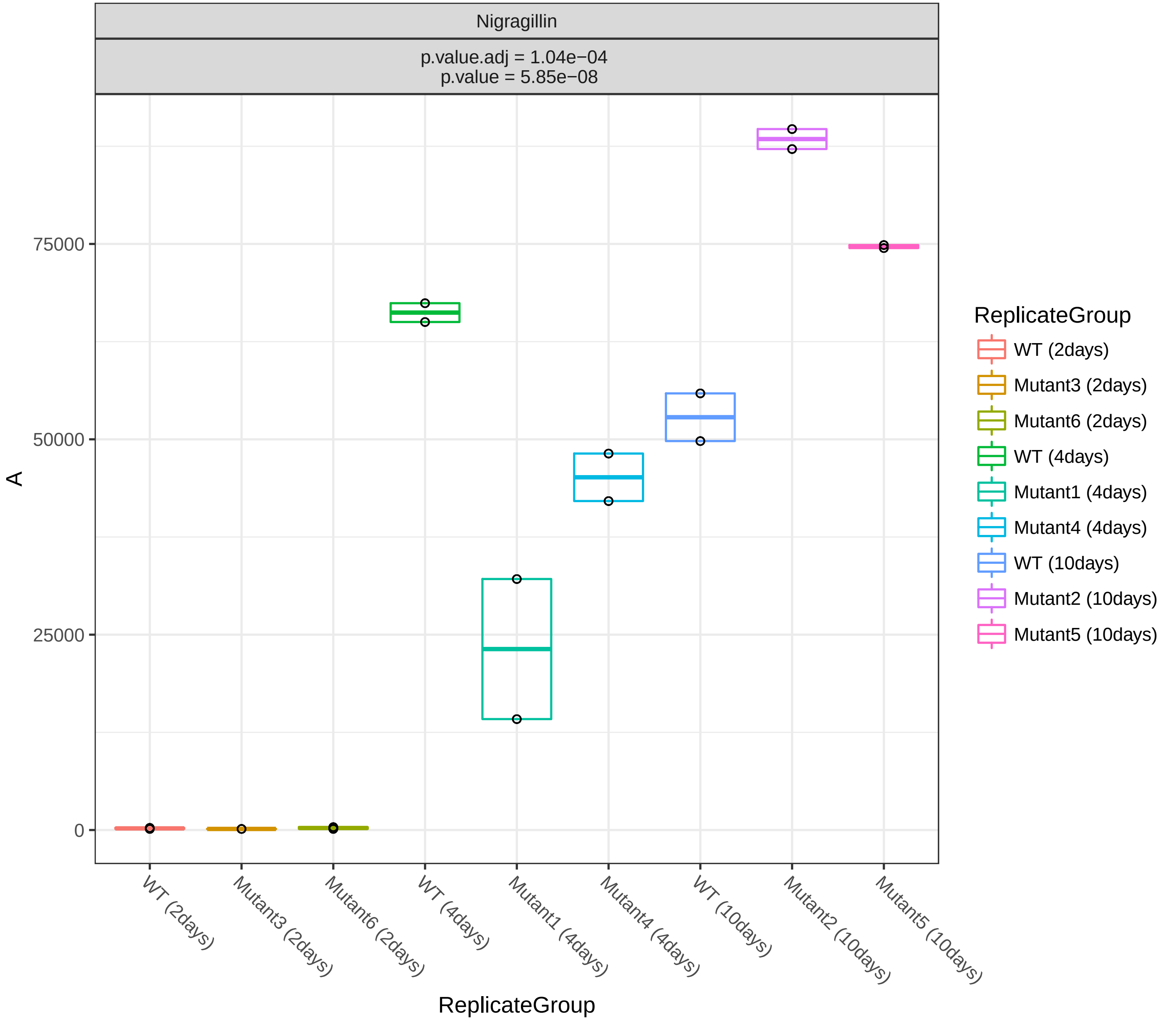

Nigragillin
p.value.adj = 1.04e−04
p.value = 5.85e−08
A
0
25000
50000
75000
ReplicateGroup
WT (2days)
Mutant3 (2days)
Mutant6 (2days)
WT (4days)
Mutant1 (4days)
Mutant4 (4days)
WT (10days)
Mutant2 (10days)
Mutant5 (10days)

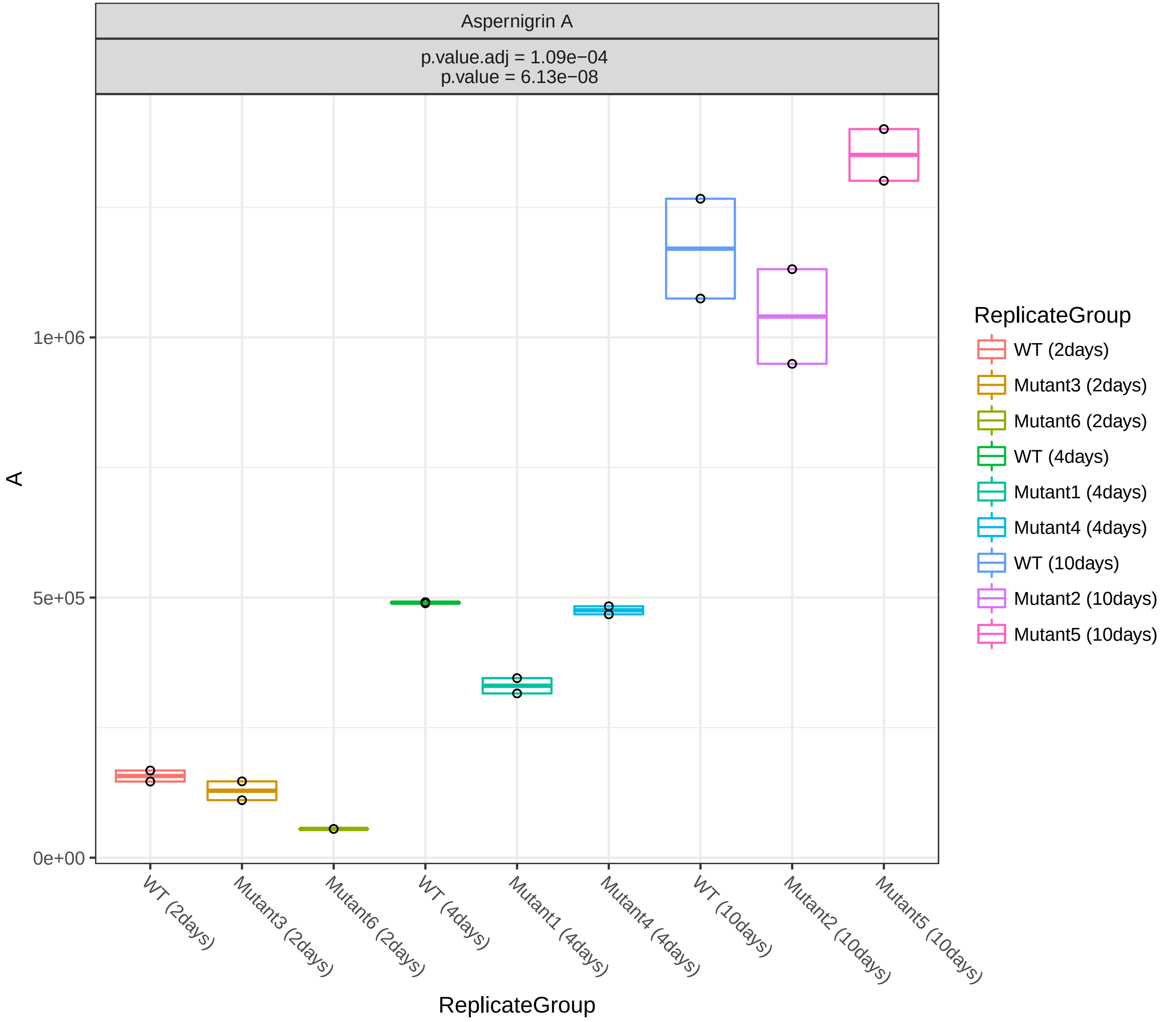
Aspernigrin A
p.value.adj = 1.09e−04
p.value = 6.13e−08
A
1e+06
5e+05
0e+00
WT (2days)
Mutant3 (2days)
Mutant6 (2days)
WT (4days)
Mutant1 (4days)
Mutant4 (4days)
WT (10days)
Mutant2 (10days)
Mutant5 (10days)
ReplicateGroup
ReplicateGroup
WT (2days)
Mutant3 (2days)
Mutant6 (2days)
WT (4days)
Mutant1 (4days)
Mutant4 (4days)
WT (10days)
Mutant2 (10days)
Mutant5 (10days)

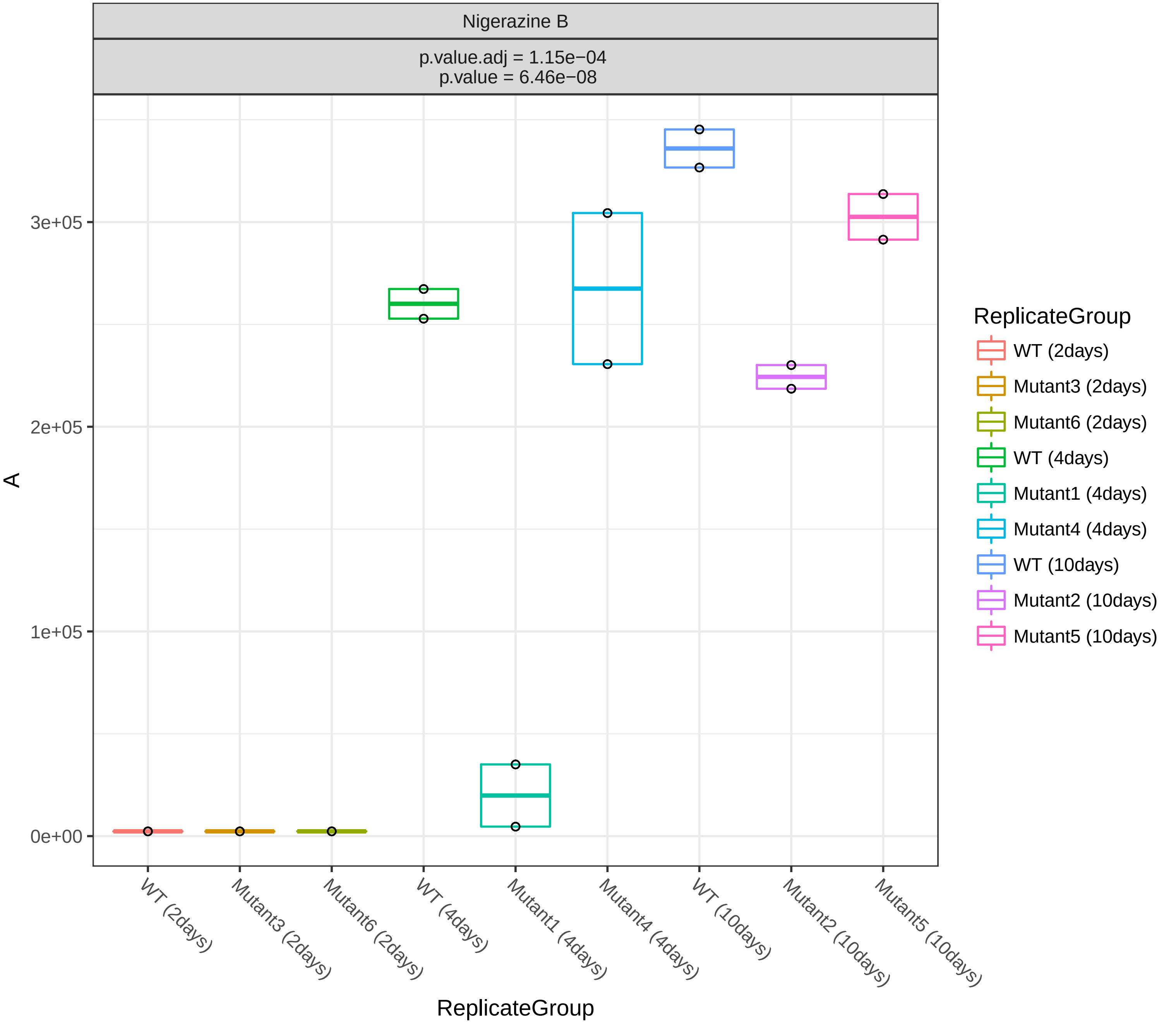
Nigerazine B
p.value.adj = 1.15e−04
p.value = 6.46e−08
A
3e+05
2e+05
1e+05
0e+00
WT (2days)
Mutant3 (2days)
Mutant6 (2days)
WT (4days)
Mutant1 (4days)
Mutant4 (4days)
WT (10days)
Mutant2 (10days)
Mutant5 (10days)
ReplicateGroup
ReplicateGroup
WT (2days)
Mutant3 (2days)
Mutant6 (2days)
WT (4days)
Mutant1 (4days)
Mutant4 (4days)
WT (10days)
Mutant2 (10days)
Mutant5 (10days)

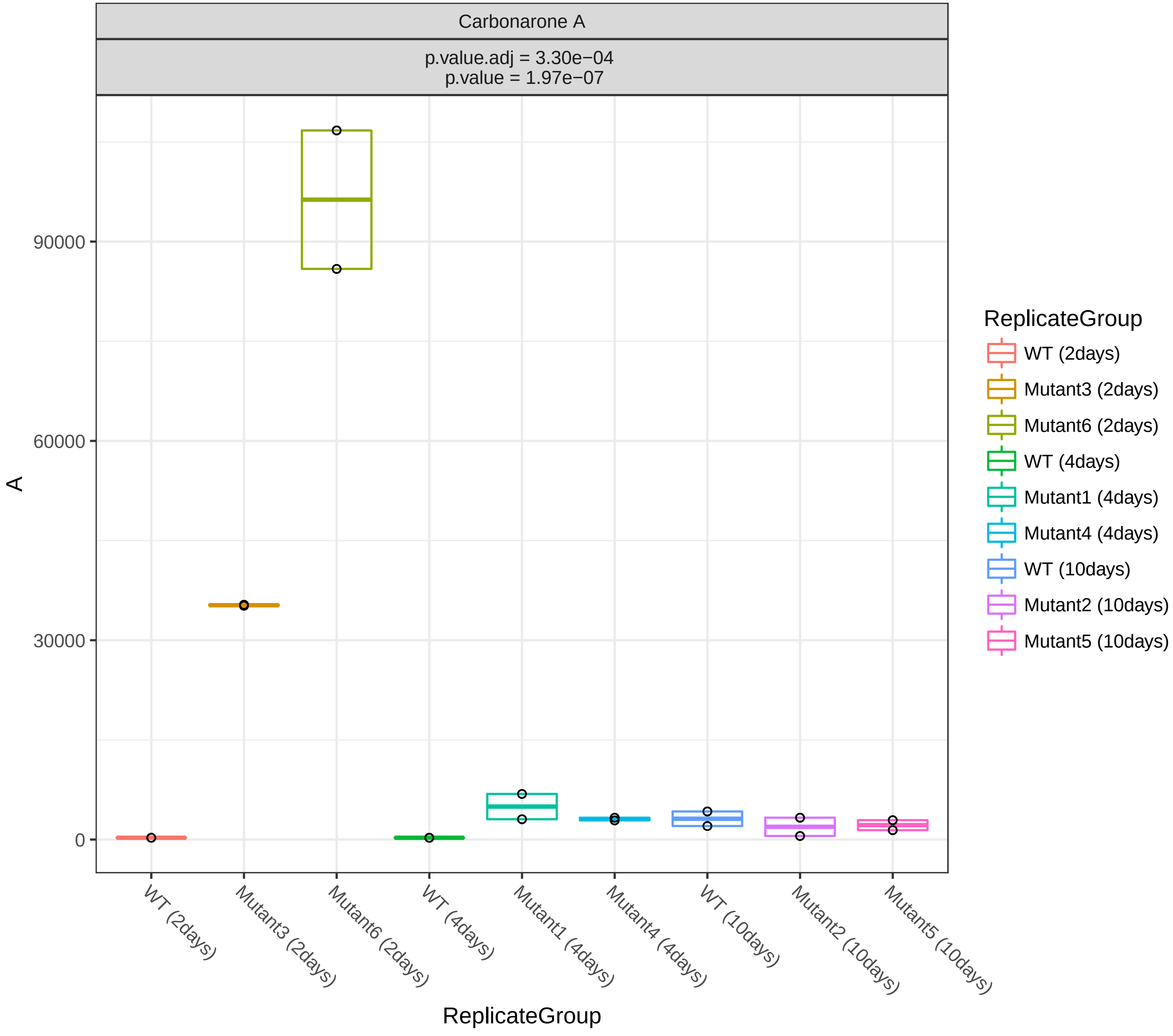

Carbonarone A
p.value.adj = 3.30e−04
p.value = 1.97e−07
A
90000
60000
30000
0
ReplicateGroup
WT (2days)
Mutant3 (2days)
Mutant6 (2days)
WT (4days)
Mutant1 (4days)
Mutant4 (4days)
WT (10days)
Mutant2 (10days)
Mutant5 (10days)

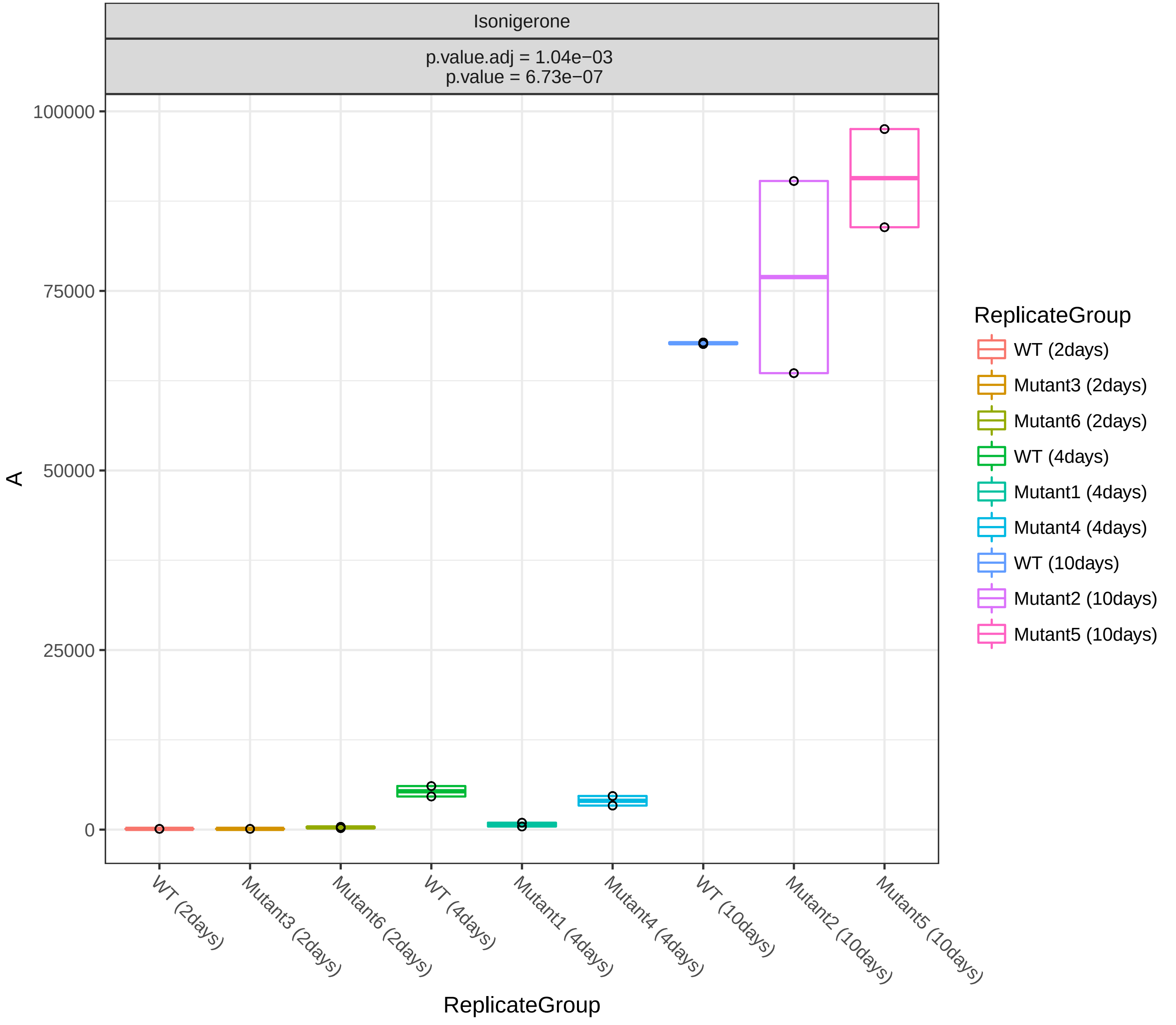

Isonigerone
p.value.adj = 1.04e−03
p.value = 6.73e−07
A
100000
75000
50000
25000
0
WT (2days)
Mutant3 (2days)
Mutant6 (2days)
WT (4days)
Mutant1 (4days)
Mutant4 (4days)
WT (10days)
Mutant2 (10days)
Mutant5 (10days)
ReplicateGroup
ReplicateGroup
WT (2days)
Mutant3 (2days)
Mutant6 (2days)
WT (4days)
Mutant1 (4days)
Mutant4 (4days)
WT (10days)
Mutant2 (10days)
Mutant5 (10days)

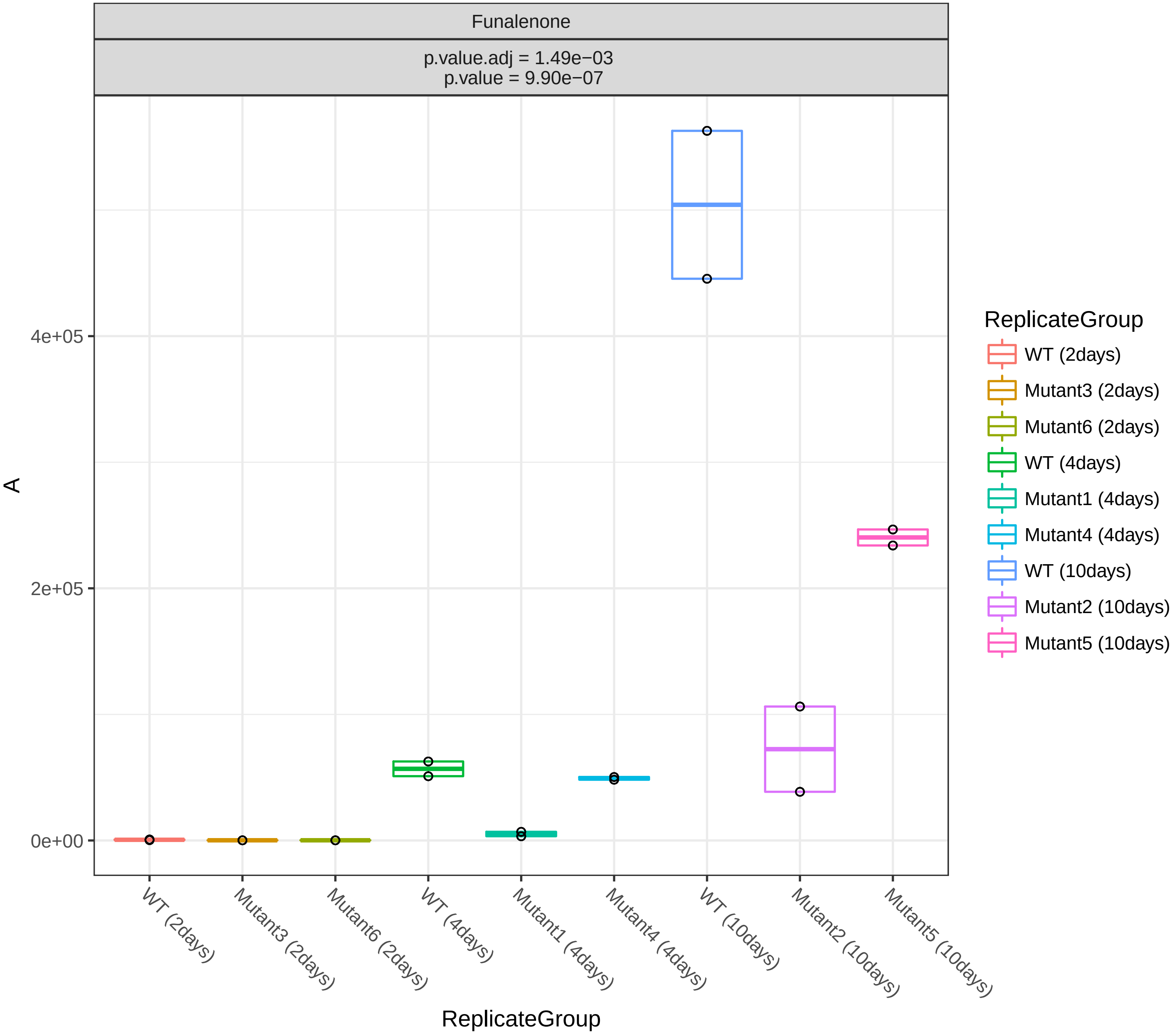

Funalenone
p.value.adj = 1.49e−03
p.value = 9.90e−07
A
4e+05
2e+05
0e+00
WT (2days)
Mutant3 (2days)
Mutant6 (2days)
WT (4days)
Mutant1 (4days)
Mutant4 (4days)
WT (10days)
Mutant2 (10days)
Mutant5 (10days)
ReplicateGroup
ReplicateGroup
WT (2days)
Mutant3 (2days)
Mutant6 (2days)
WT (4days)
Mutant1 (4days)
Mutant4 (4days)
WT (10days)
Mutant2 (10days)
Mutant5 (10days)

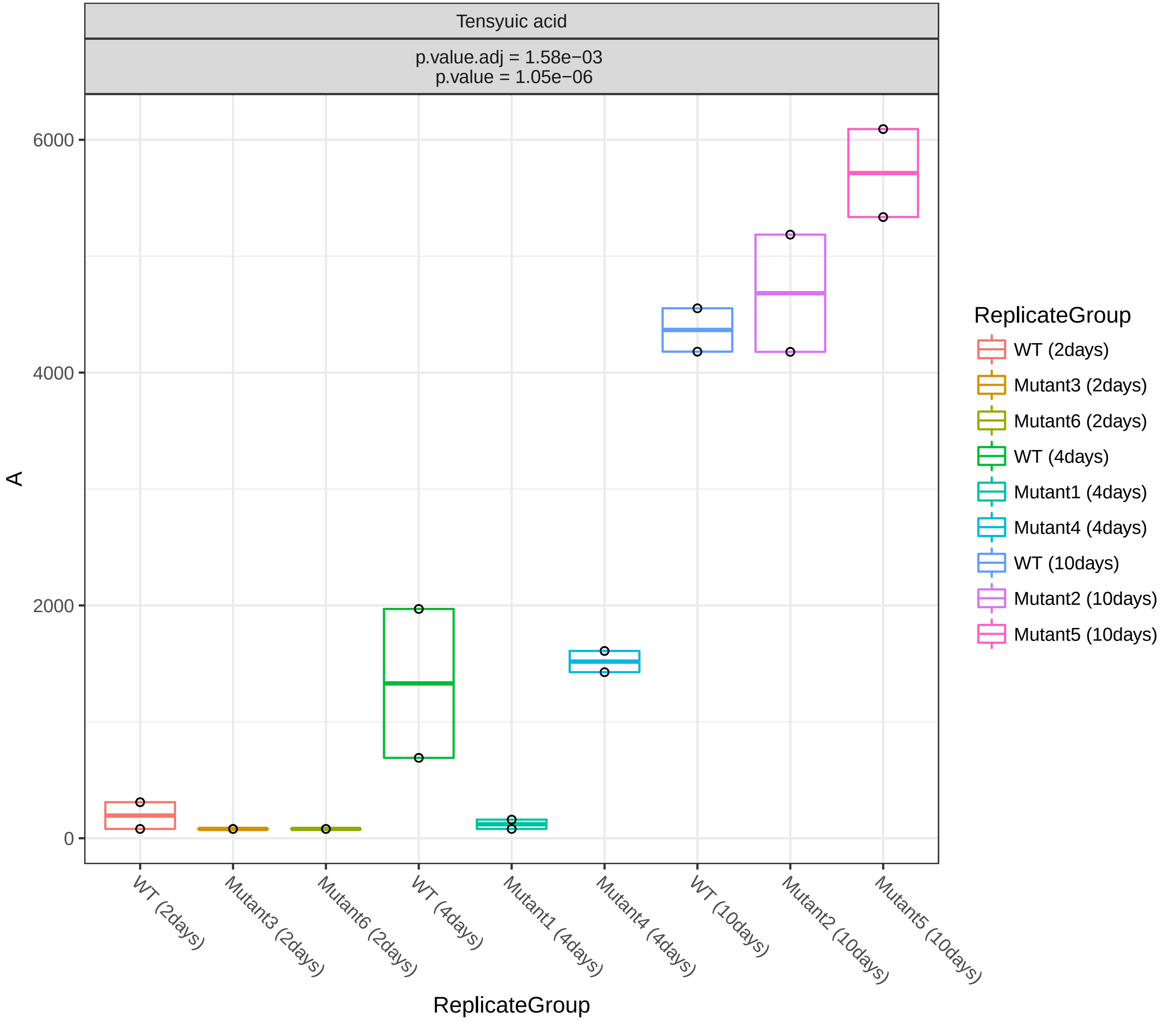
Tensyuic acid
p.value.adj = 1.58e−03
p.value = 1.05e−06
6000
4000
2000
0
A
WT (2days)
Mutant3 (2days)
Mutant6 (2days)
WT (4days)
Mutant1 (4days)
Mutant4 (4days)
WT (10days)
Mutant2 (10days)
Mutant5 (10days)
ReplicateGroup
ReplicateGroup
WT (2days)
Mutant3 (2days)
Mutant6 (2days)
WT (4days)
Mutant1 (4days)
Mutant4 (4days)
WT (10days)
Mutant2 (10days)
Mutant5 (10days)

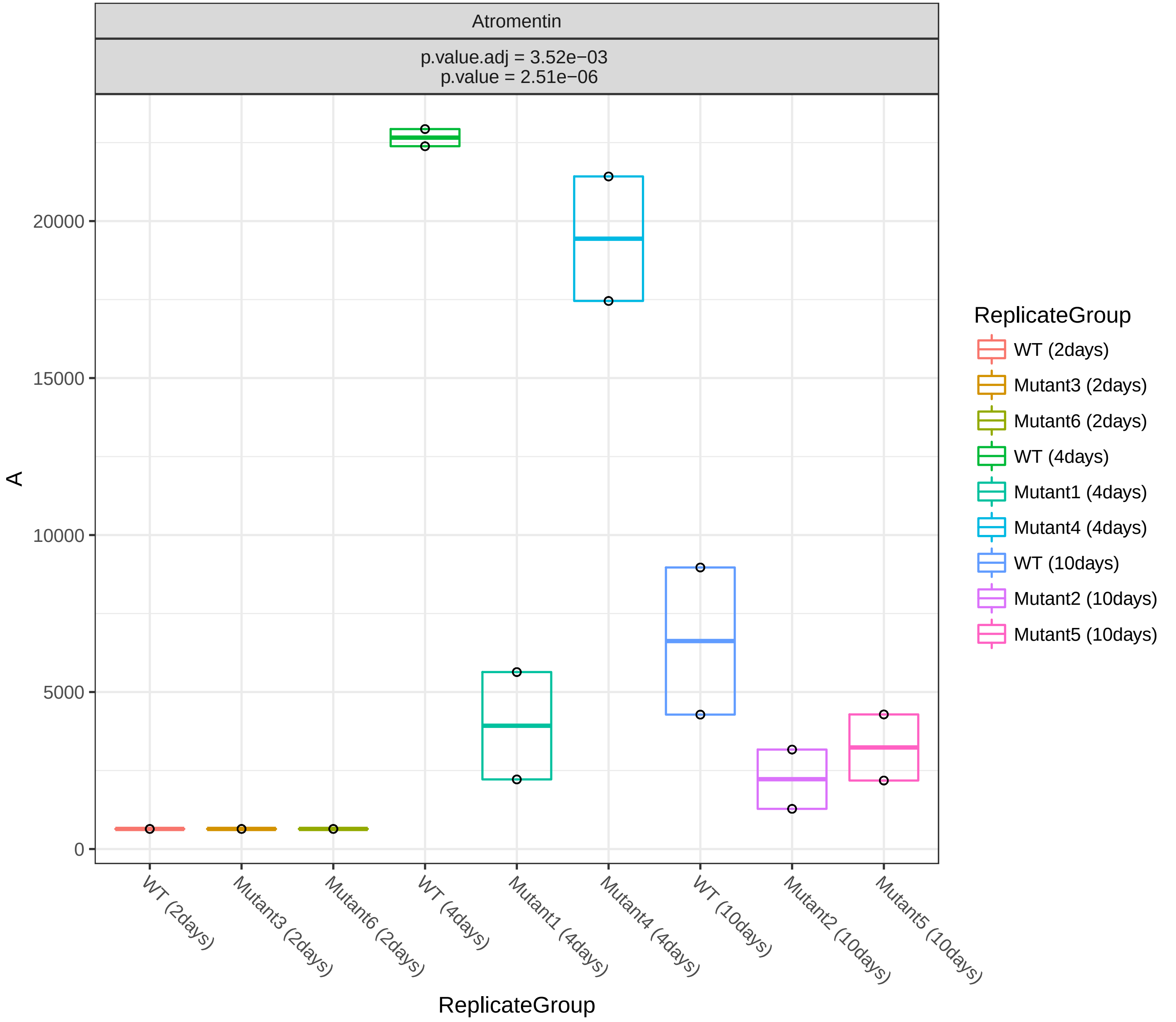
Atromentin
p.value.adj = 3.52e−03
p.value = 2.51e−06
A
20000
15000
10000
5000
0
WT (2days)
Mutant3 (2days)
Mutant6 (2days)
WT (4days)
Mutant1 (4days)
Mutant4 (4days)
WT (10days)
Mutant2 (10days)
Mutant5 (10days)
ReplicateGroup
ReplicateGroup
WT (2days)
Mutant3 (2days)
Mutant6 (2days)
WT (4days)
Mutant1 (4days)
Mutant4 (4days)
WT (10days)
Mutant2 (10days)
Mutant5 (10days)

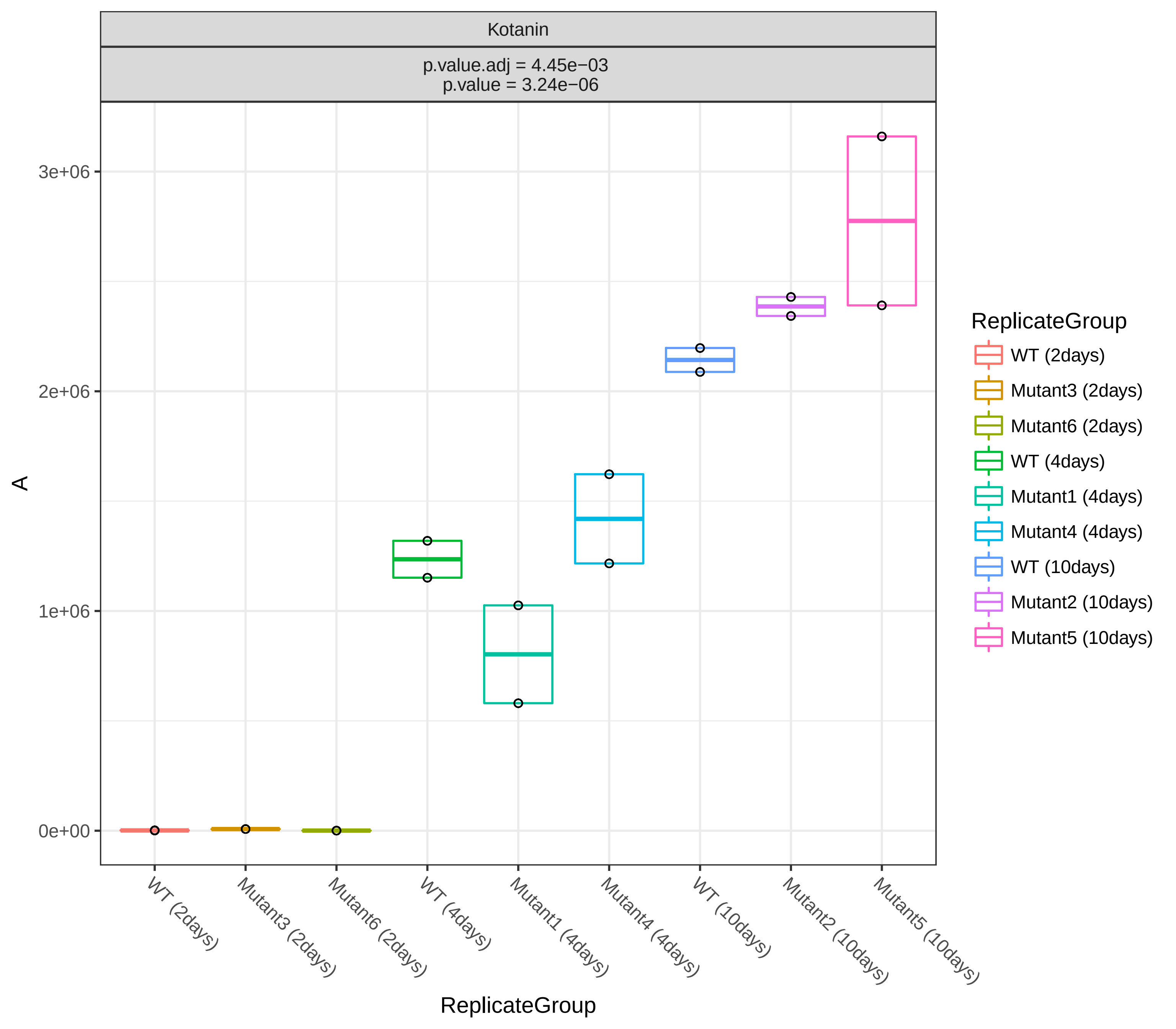
Kotanin
p.value.adj = 4.45e−03
p.value = 3.24e−06
3e+06
2e+06
1e+06
0e+00
A
WT (2days)
Mutant3 (2days)
Mutant6 (2days)
WT (4days)
Mutant1 (4days)
Mutant4 (4days)
WT (10days)
Mutant2 (10days)
Mutant5 (10days)
ReplicateGroup
ReplicateGroup
WT (2days)
Mutant3 (2days)
Mutant6 (2days)
WT (4days)
Mutant1 (4days)
Mutant4 (4days)
WT (10days)
Mutant2 (10days)
Mutant5 (10days)

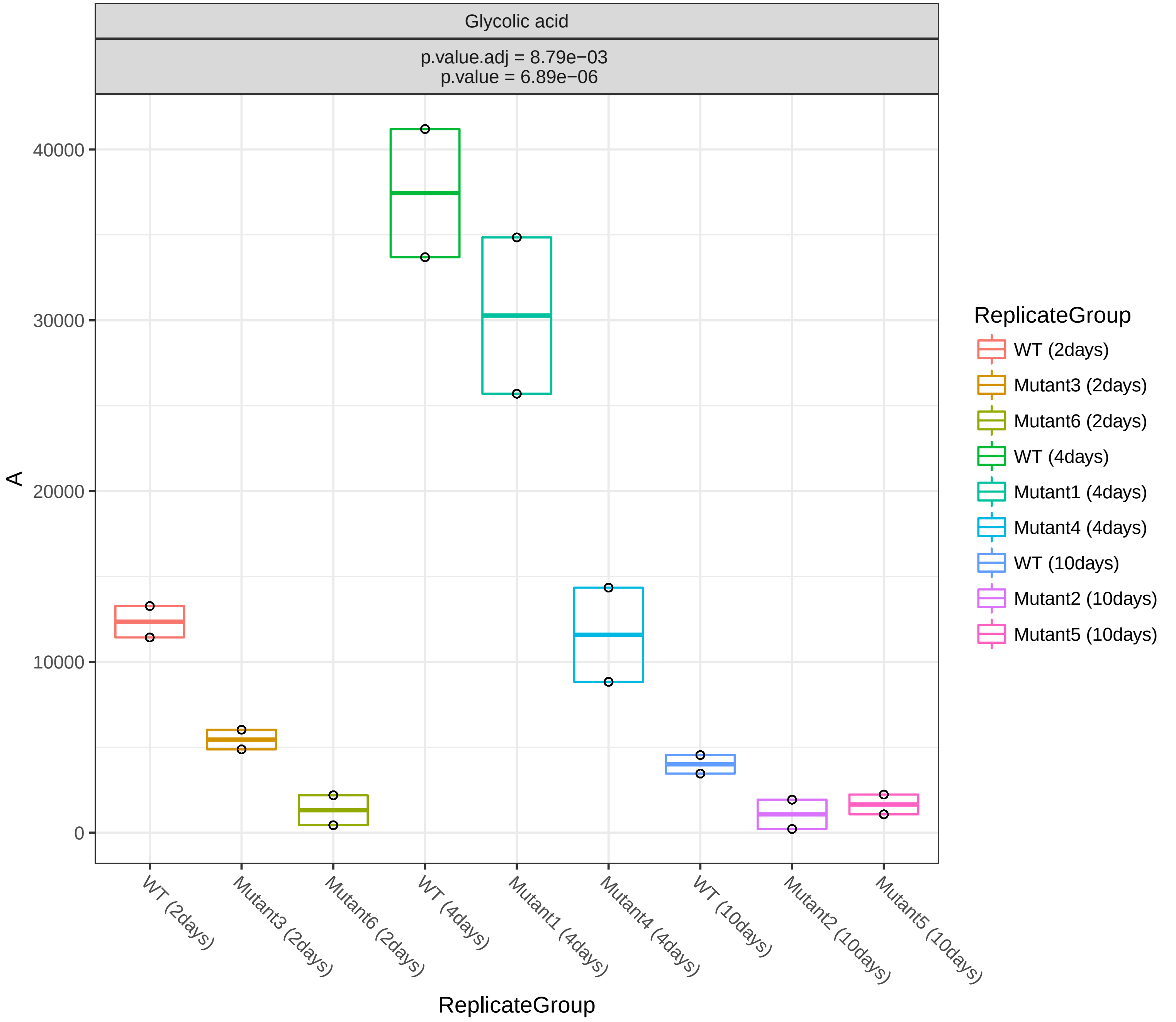
Glycolic acid
p.value.adj = 8.79e−03
p.value = 6.89e−06
A
40000
30000
20000
10000
0
WT (2days)
Mutant3 (2days)
Mutant6 (2days)
WT (4days)
Mutant1 (4days)
Mutant4 (4days)
WT (10days)
Mutant2 (10days)
Mutant5 (10days)
ReplicateGroup
ReplicateGroup
WT (2days)
Mutant3 (2days)
Mutant6 (2days)
WT (4days)
Mutant1 (4days)
Mutant4 (4days)
WT (10days)
Mutant2 (10days)
Mutant5 (10days)

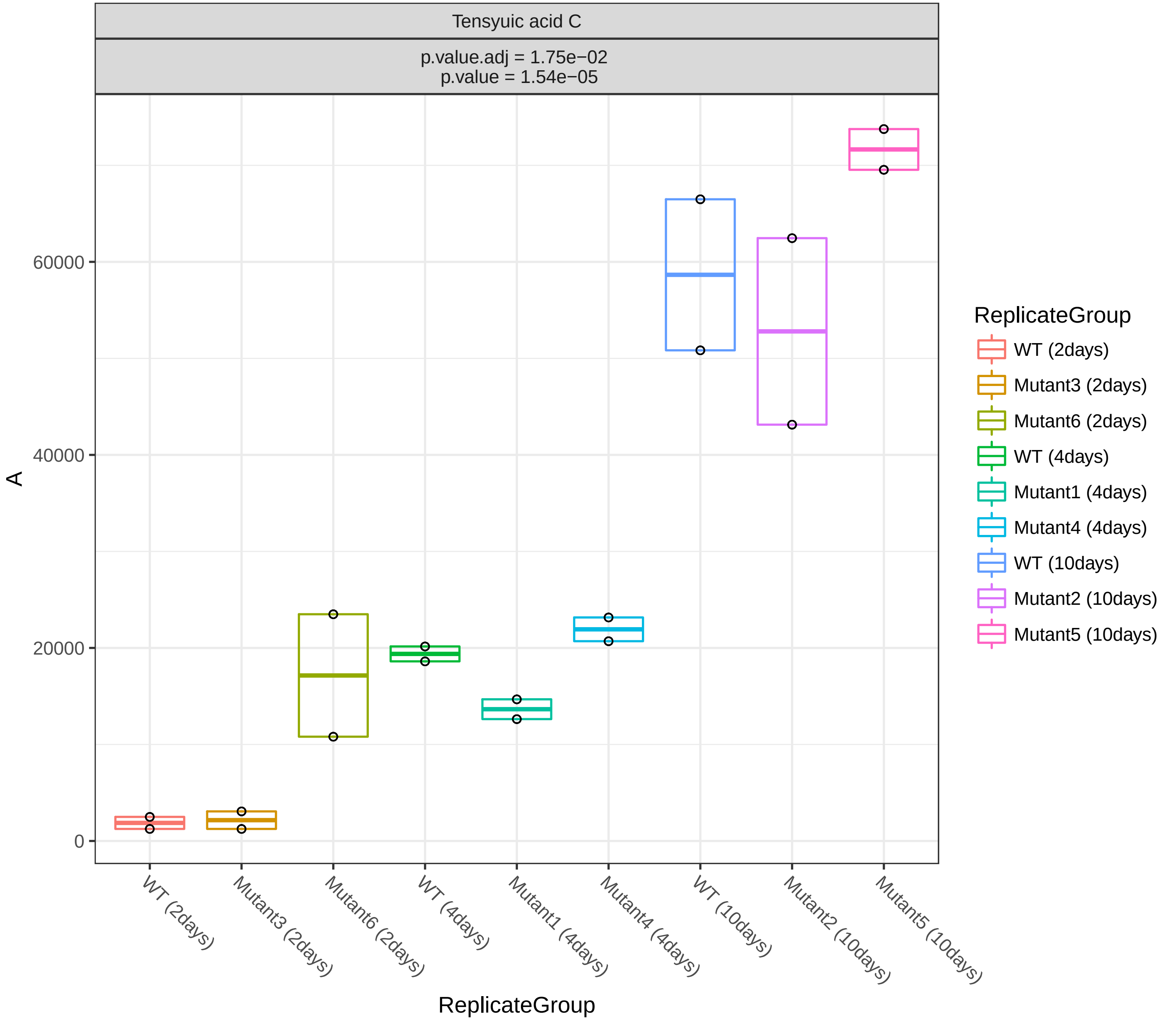
Tensyuic acid C
p.value.adj = 1.75e−02
p.value = 1.54e−05
A
0
20000
40000
60000
ReplicateGroup
WT (2days)
Mutant3 (2days)
Mutant6 (2days)
WT (4days)
Mutant1 (4days)
Mutant4 (4days)
WT (10days)
Mutant2 (10days)
Mutant5 (10days)

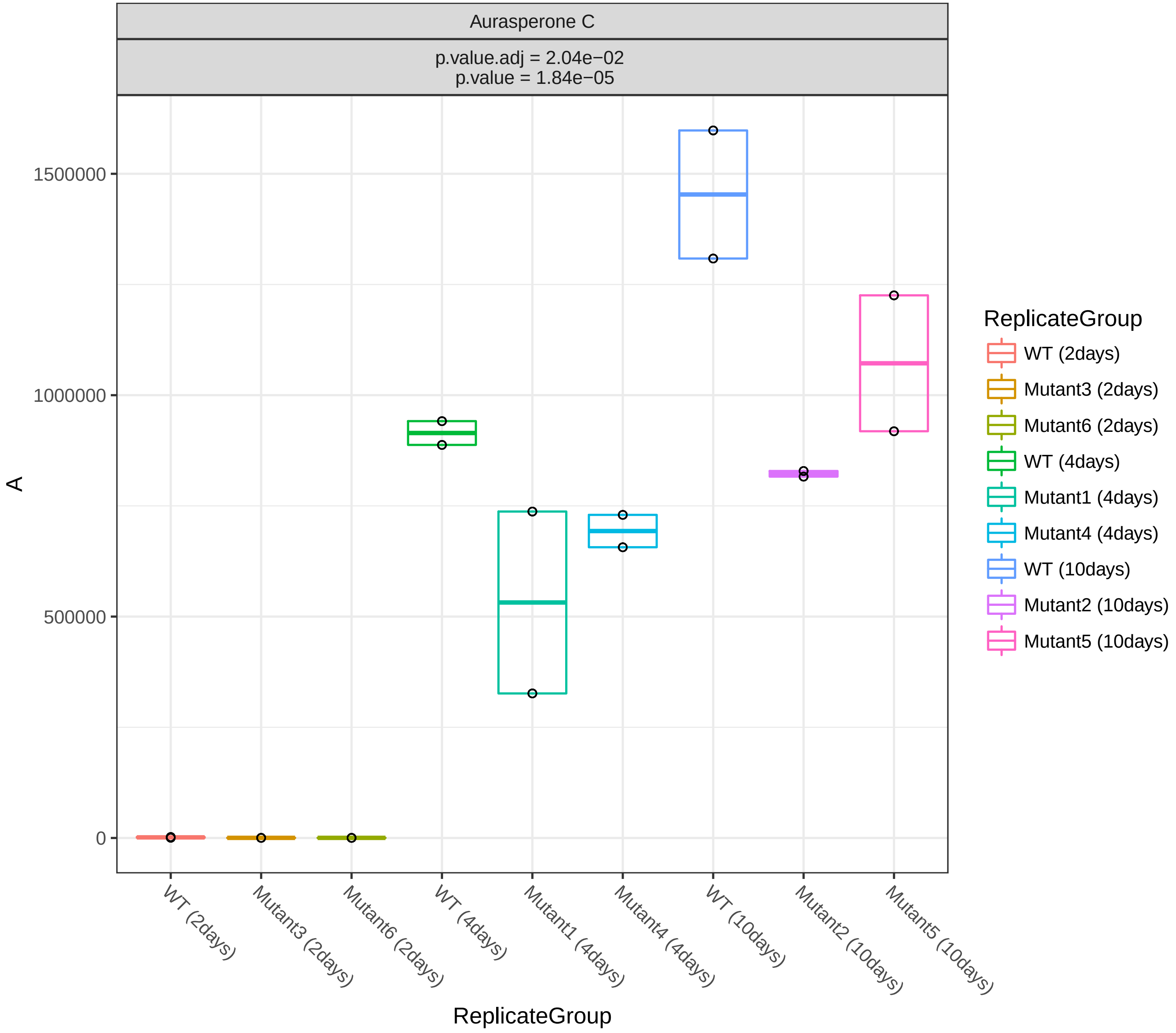
Aurasperone C
p.value.adj = 2.04e−02
p.value = 1.84e−05
1500000
1000000
500000
0
A
WT (2days)
Mutant3 (2days)
Mutant6 (2days)
WT (4days)
Mutant1 (4days)
Mutant4 (4days)
WT (10days)
Mutant2 (10days)
Mutant5 (10days)
ReplicateGroup
ReplicateGroup
WT (2days)
Mutant3 (2days)
Mutant6 (2days)
WT (4days)
Mutant1 (4days)
Mutant4 (4days)
WT (10days)
Mutant2 (10days)
Mutant5 (10days)

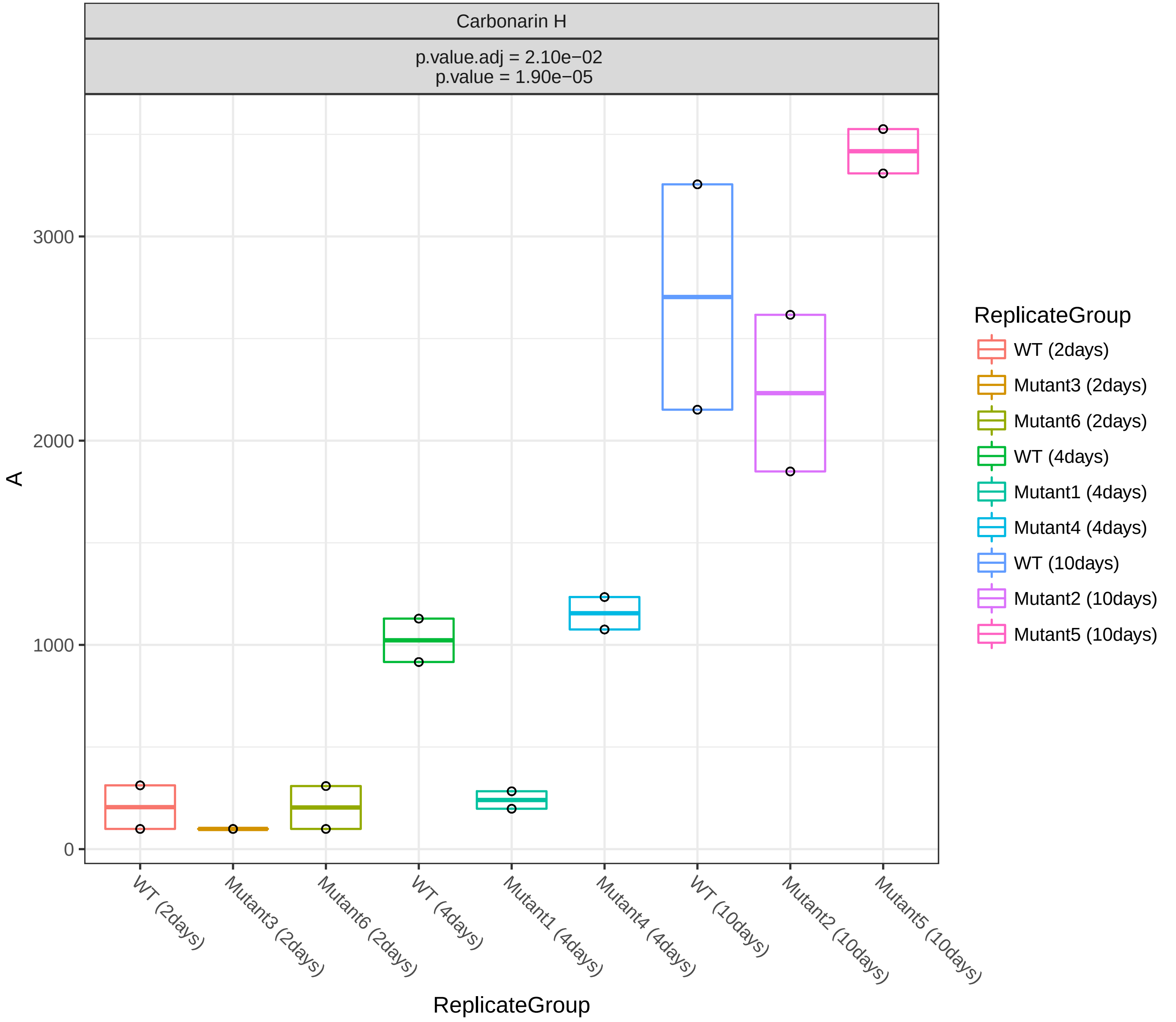

Carbonarin H
p.value.adj = 2.10e−02
p.value = 1.90e−05
A
3000
2000
1000
0
WT (2days)
Mutant3 (2days)
Mutant6 (2days)
WT (4days)
Mutant1 (4days)
Mutant4 (4days)
WT (10days)
Mutant2 (10days)
Mutant5 (10days)
ReplicateGroup
ReplicateGroup
WT (2days)
Mutant3 (2days)
Mutant6 (2days)
WT (4days)
Mutant1 (4days)
Mutant4 (4days)
WT (10days)
Mutant2 (10days)
Mutant5 (10days)

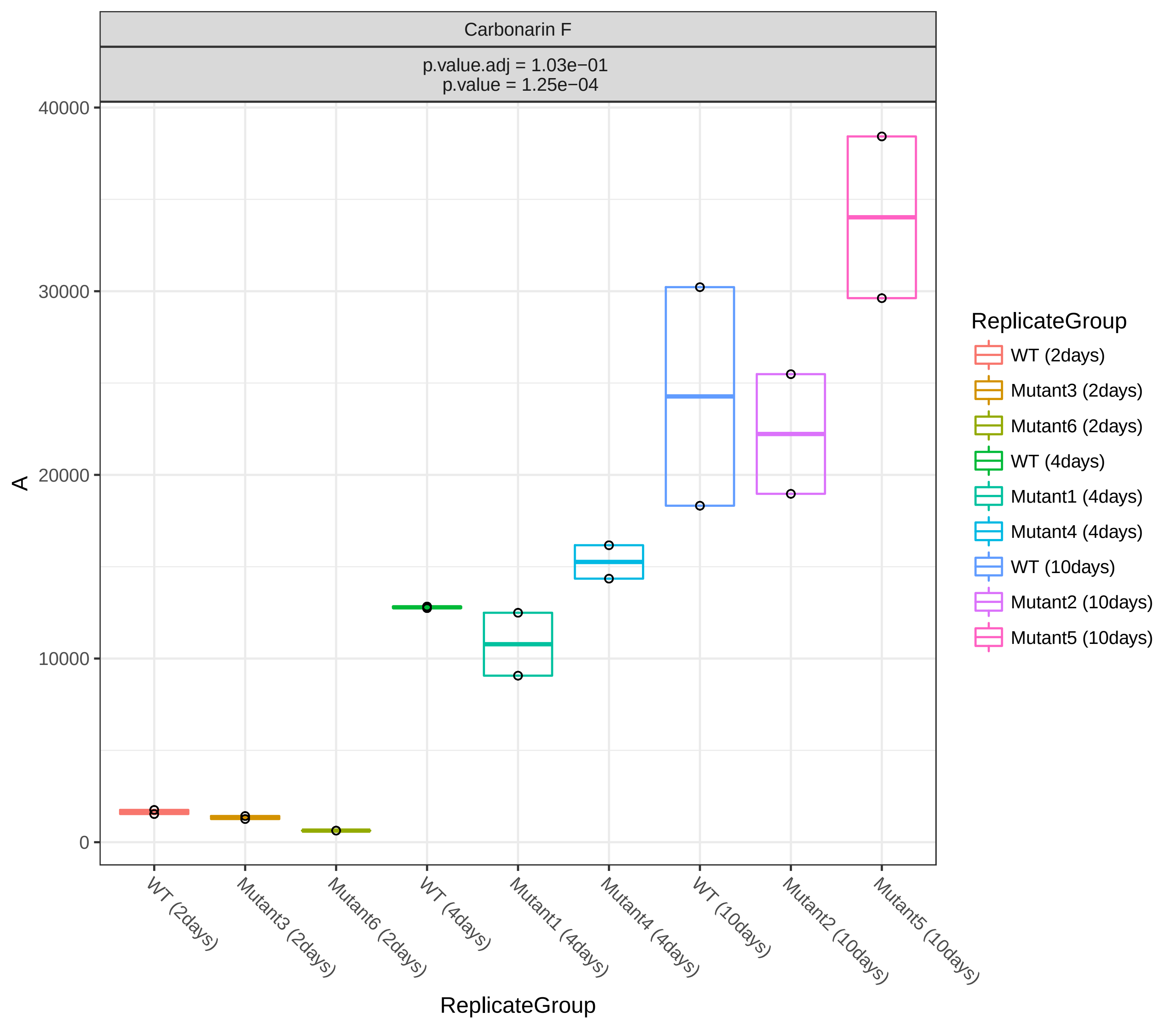
Carbonarin F
p.value.adj = 1.03e−01
p.value = 1.25e−04
A
40000
30000
20000
10000
0
WT (2days)
Mutant3 (2days)
Mutant6 (2days)
WT (4days)
Mutant1 (4days)
Mutant4 (4days)
WT (10days)
Mutant2 (10days)
Mutant5 (10days)
ReplicateGroup
ReplicateGroup
WT (2days)
Mutant3 (2days)
Mutant6 (2days)
WT (4days)
Mutant1 (4days)
Mutant4 (4days)
WT (10days)
Mutant2 (10days)
Mutant5 (10days)

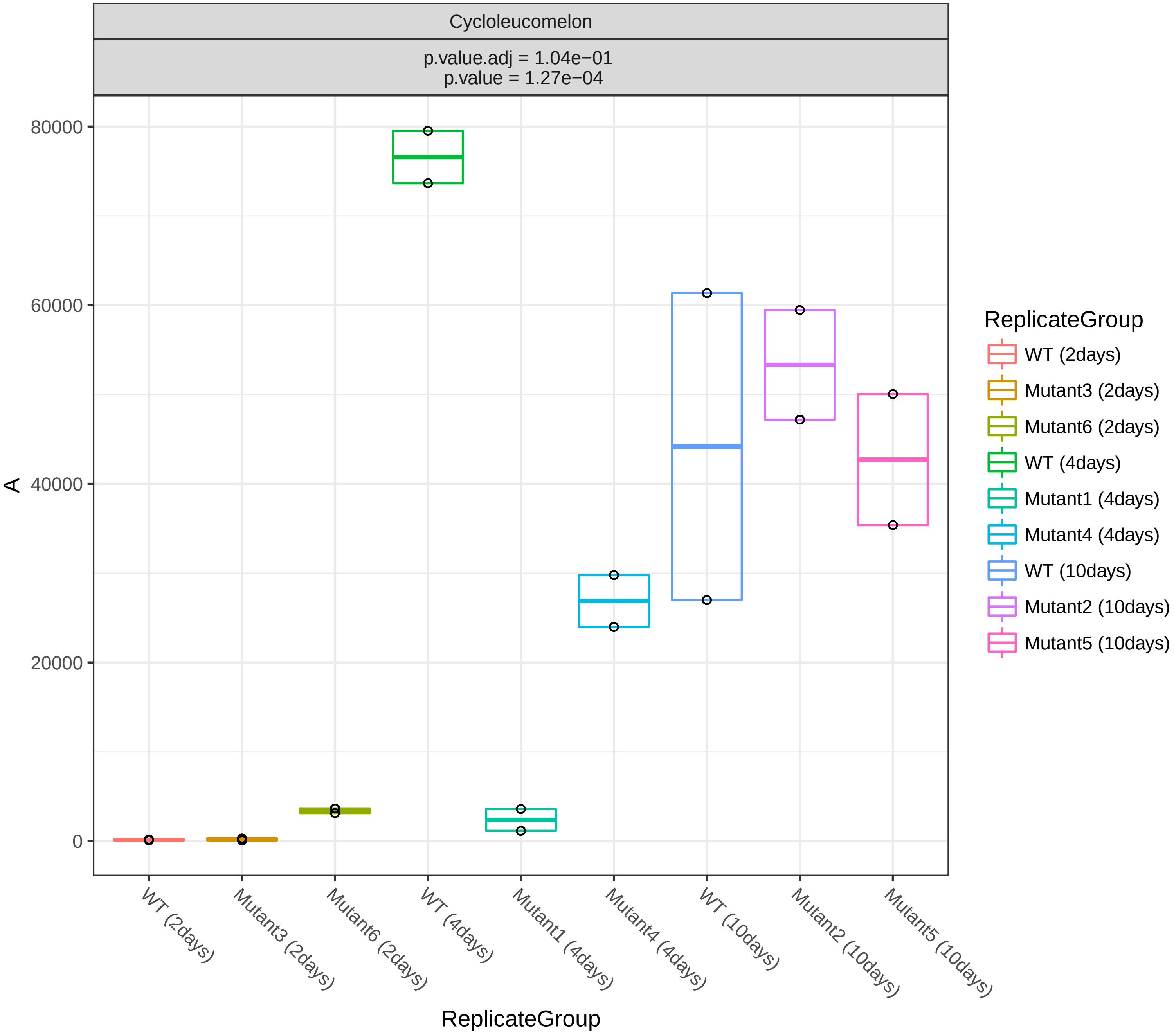
Cycloleucomelon
p.value.adj = 1.04e−01
p.value = 1.27e−04
A
80000
60000
40000
20000
0
ReplicateGroup
WT (2days)
Mutant3 (2days)
Mutant6 (2days)
WT (4days)
Mutant1 (4days)
Mutant4 (4days)
WT (10days)
Mutant2 (10days)
Mutant5 (10days)
ReplicateGroup
WT (2days)
Mutant3 (2days)
Mutant6 (2days)
WT (4days)
Mutant1 (4days)
Mutant4 (4days)
WT (10days)
Mutant2 (10days)
Mutant5 (10days)

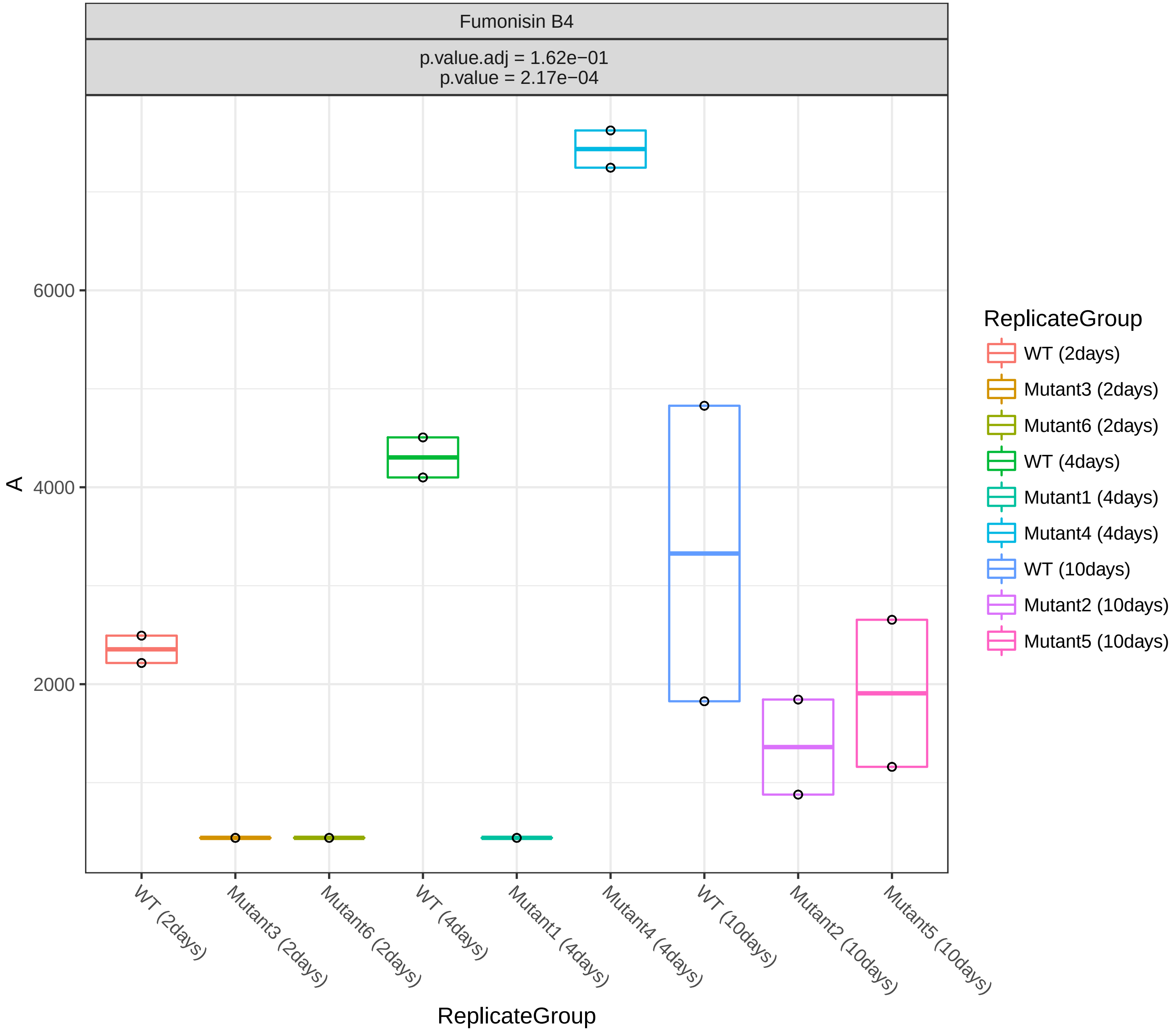
Fumonisin B4
p.value.adj = 1.62e−01
p.value = 2.17e−04
A
6000
4000
2000
ReplicateGroup
WT (2days)
Mutant3 (2days)
Mutant6 (2days)
WT (4days)
Mutant1 (4days)
Mutant4 (4days)
WT (10days)
Mutant2 (10days)
Mutant5 (10days)
ReplicateGroup
WT (2days)
Mutant3 (2days)
Mutant6 (2days)
WT (4days)
Mutant1 (4days)
Mutant4 (4days)
WT (10days)
Mutant2 (10days)
Mutant5 (10days)

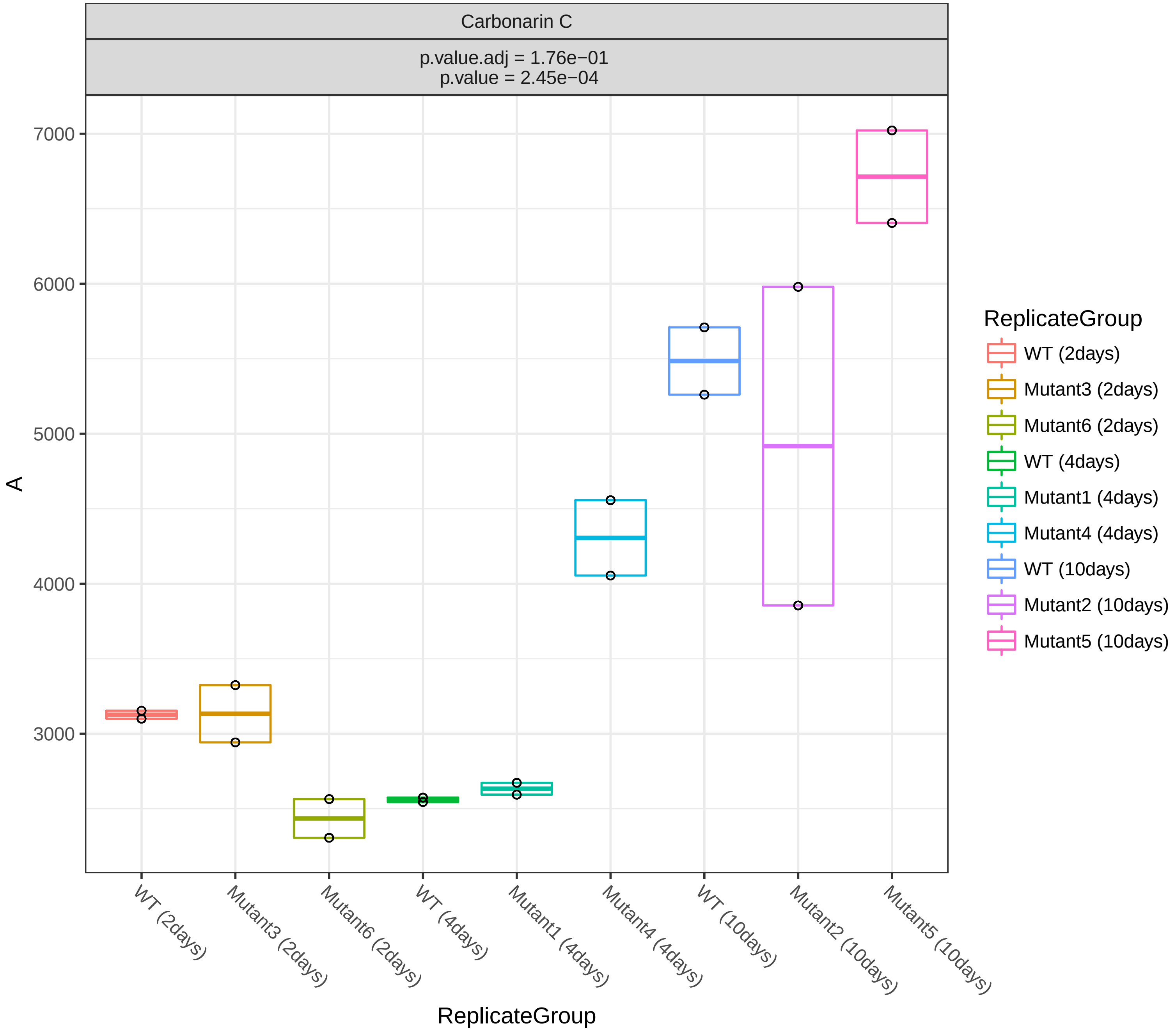

Carbonarin C
p.value.adj = 1.76e−01
p.value = 2.45e−04
7000
6000
5000
4000
3000
A
WT (2days)
Mutant3 (2days)
Mutant6 (2days)
WT (4days)
Mutant1 (4days)
Mutant4 (4days)
WT (10days)
Mutant2 (10days)
Mutant5 (10days)
ReplicateGroup
ReplicateGroup
WT (2days)
Mutant3 (2days)
Mutant6 (2days)
WT (4days)
Mutant1 (4days)
Mutant4 (4days)
WT (10days)
Mutant2 (10days)
Mutant5 (10days)

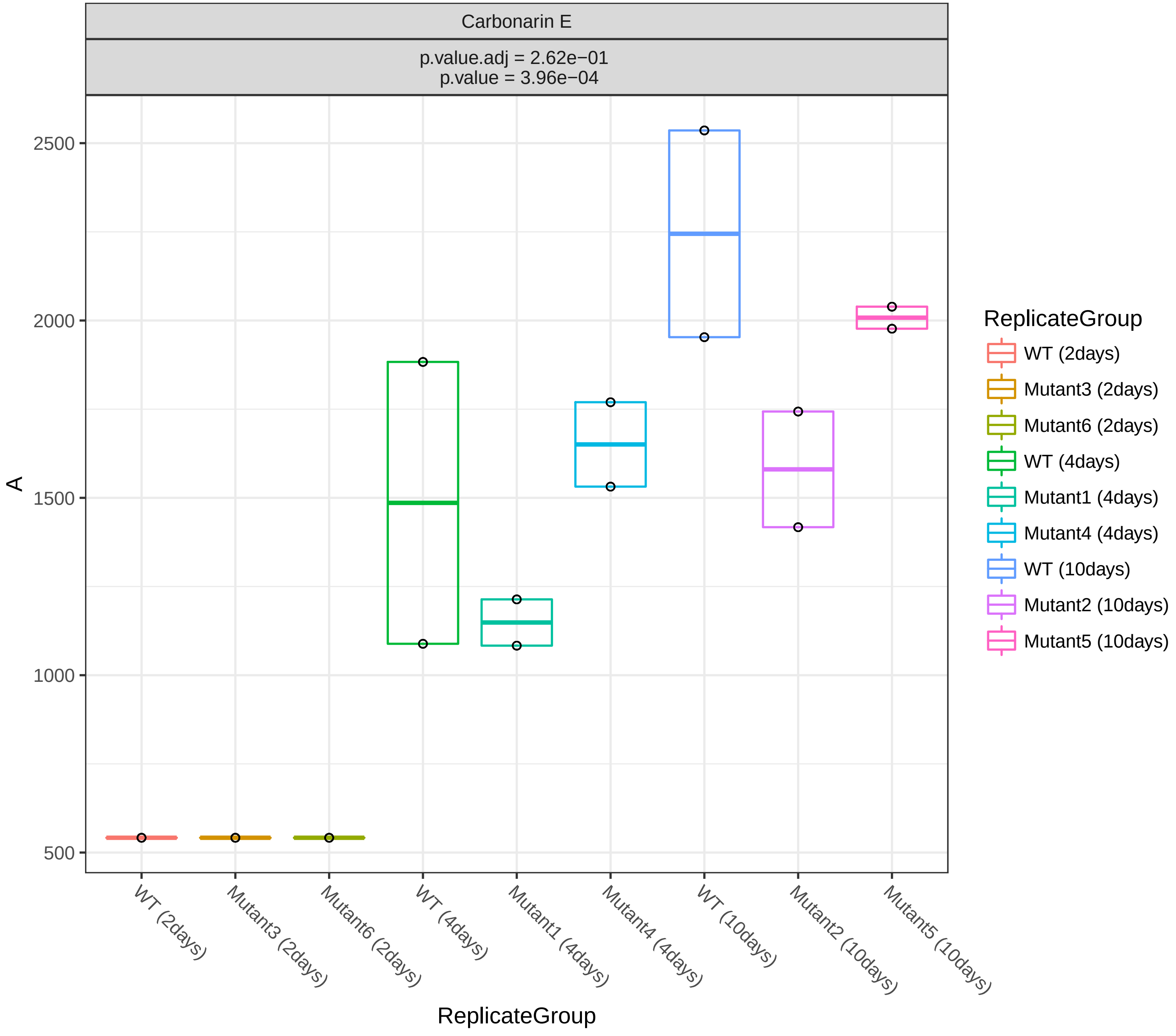
Carbonarin E
p.value.adj = 2.62e−01
p.value = 3.96e−04
2500
2000
1500
1000
500
A
WT (2days)
Mutant3 (2days)
Mutant6 (2days)
WT (4days)
Mutant1 (4days)
Mutant4 (4days)
WT (10days)
Mutant2 (10days)
Mutant5 (10days)
ReplicateGroup
ReplicateGroup
WT (2days)
Mutant3 (2days)
Mutant6 (2days)
WT (4days)
Mutant1 (4days)
Mutant4 (4days)
WT (10days)
Mutant2 (10days)
Mutant5 (10days)

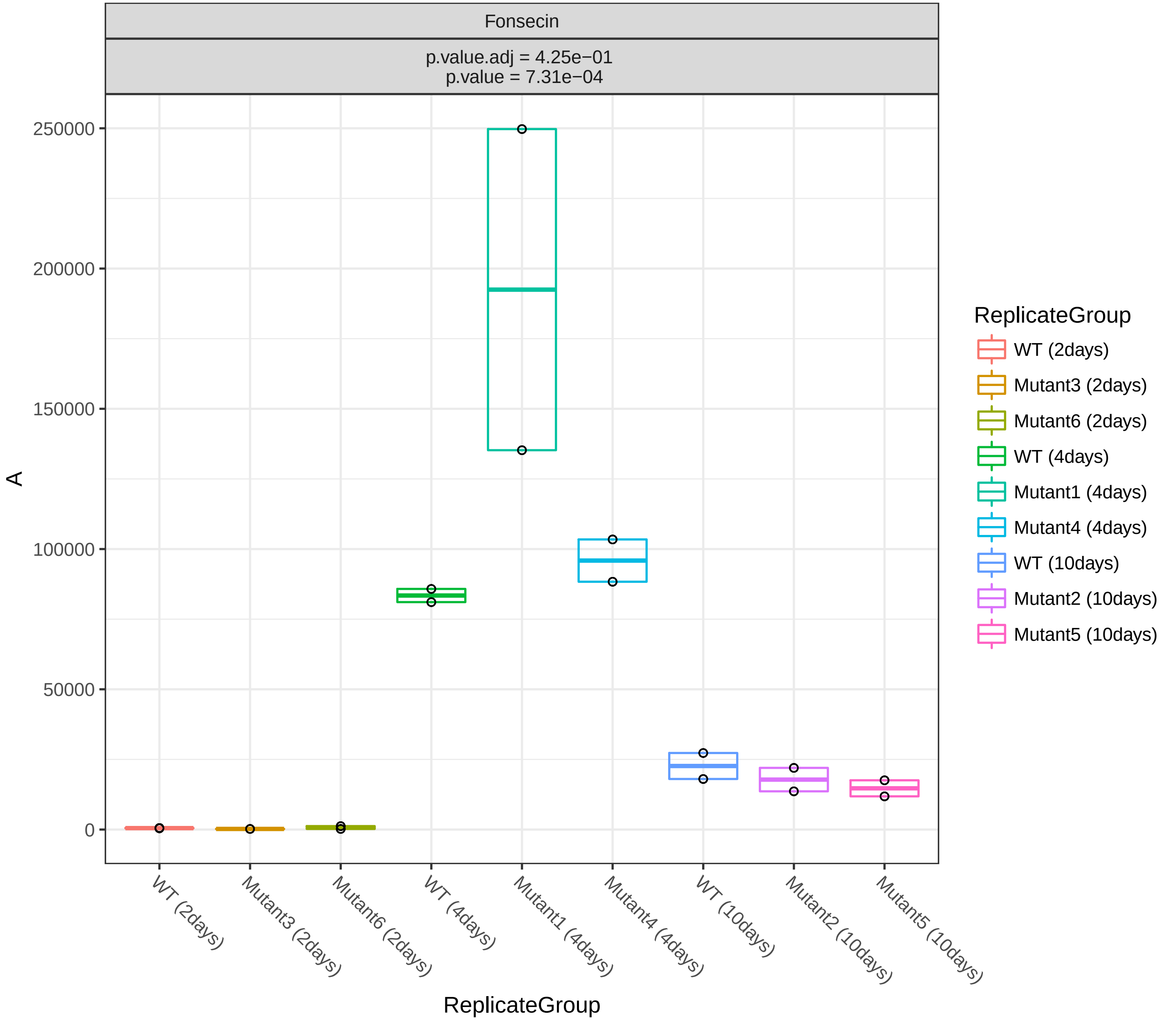
Fonsecin
p.value.adj = 4.25e−01
p.value = 7.31e−04
A
250000
200000
150000
100000
50000
0
WT (2days)
Mutant3 (2days)
Mutant6 (2days)
WT (4days)
Mutant1 (4days)
Mutant4 (4days)
WT (10days)
Mutant2 (10days)
Mutant5 (10days)
ReplicateGroup
ReplicateGroup
WT (2days)
Mutant3 (2days)
Mutant6 (2days)
WT (4days)
Mutant1 (4days)
Mutant4 (4days)
WT (10days)
Mutant2 (10days)
Mutant5 (10days)

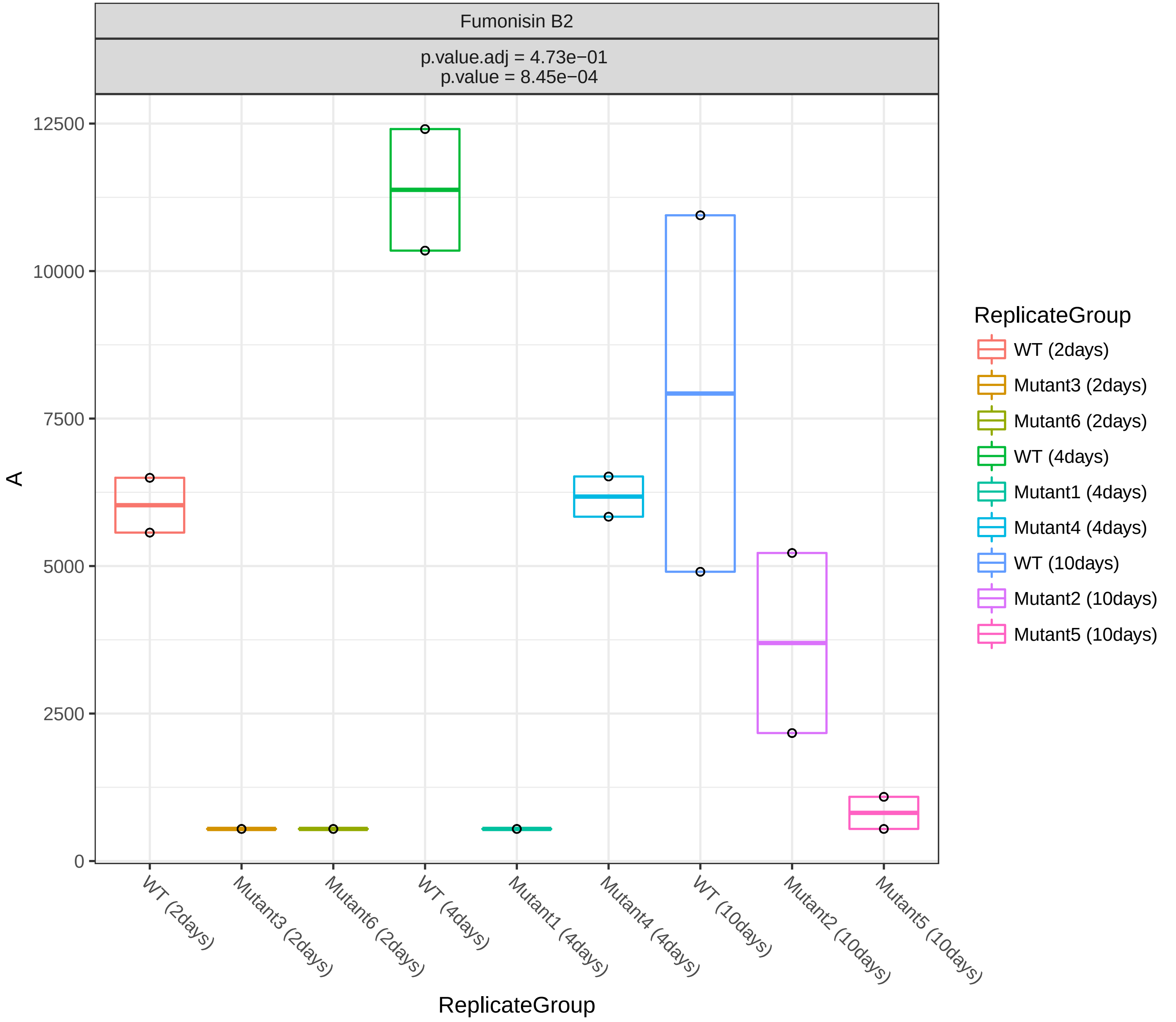
Fumonisin B2
p.value.adj = 4.73e−01
p.value = 8.45e−04
12500
10000
7500
5000
2500
0
A
WT (2days)
Mutant3 (2days)
Mutant6 (2days)
WT (4days)
Mutant1 (4days)
Mutant4 (4days)
WT (10days)
Mutant2 (10days)
Mutant5 (10days)
ReplicateGroup
ReplicateGroup
WT (2days)
Mutant3 (2days)
Mutant6 (2days)
WT (4days)
Mutant1 (4days)
Mutant4 (4days)
WT (10days)
Mutant2 (10days)
Mutant5 (10days)

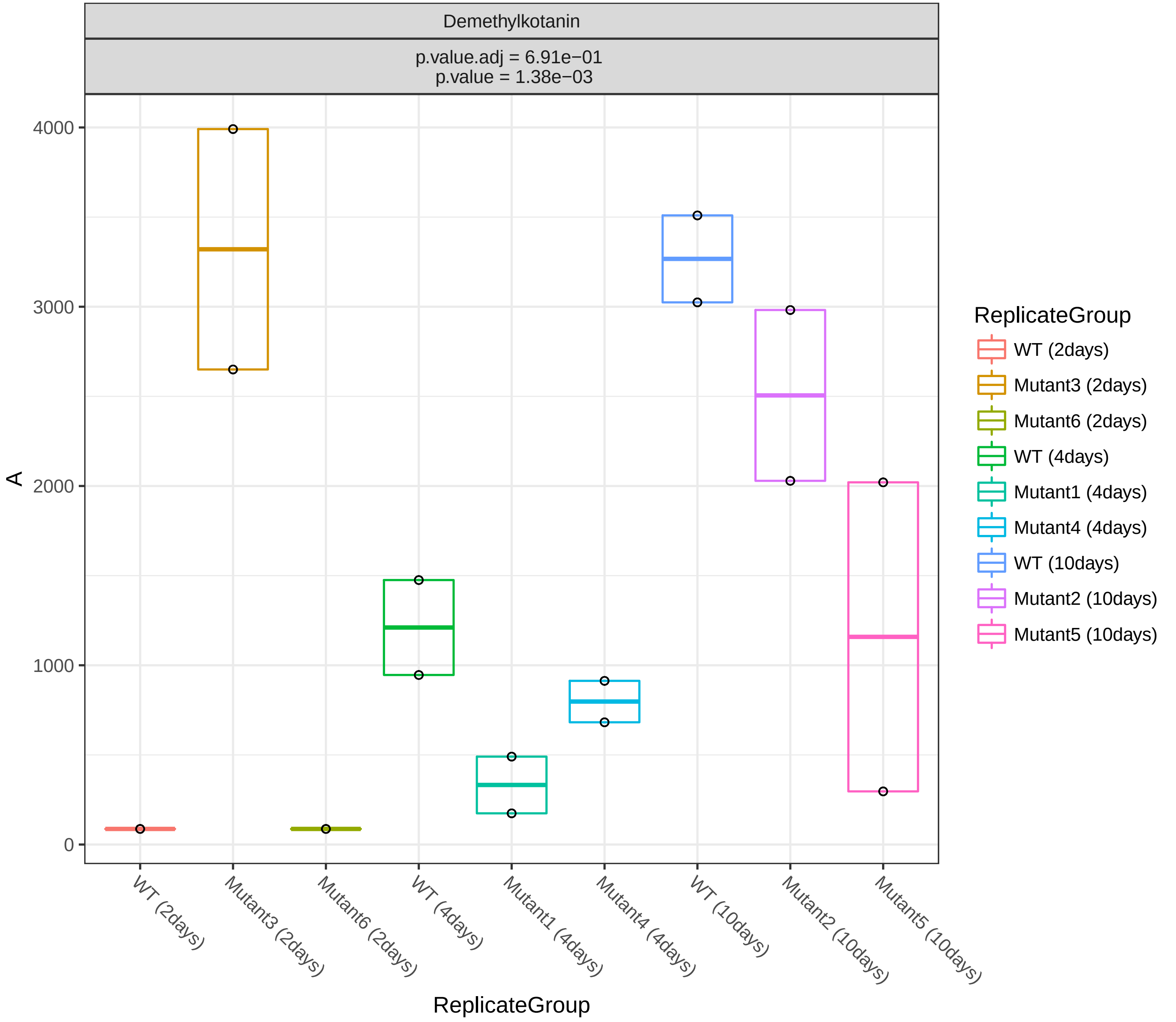
Demethylkotanin
p.value.adj = 6.91e−01
p.value = 1.38e−03
A
4000
3000
2000
1000
0
WT (2days)
Mutant3 (2days)
Mutant6 (2days)
WT (4days)
Mutant1 (4days)
Mutant4 (4days)
WT (10days)
Mutant2 (10days)
Mutant5 (10days)
ReplicateGroup
ReplicateGroup
WT (2days)
Mutant3 (2days)
Mutant6 (2days)
WT (4days)
Mutant1 (4days)
Mutant4 (4days)
WT (10days)
Mutant2 (10days)
Mutant5 (10days)

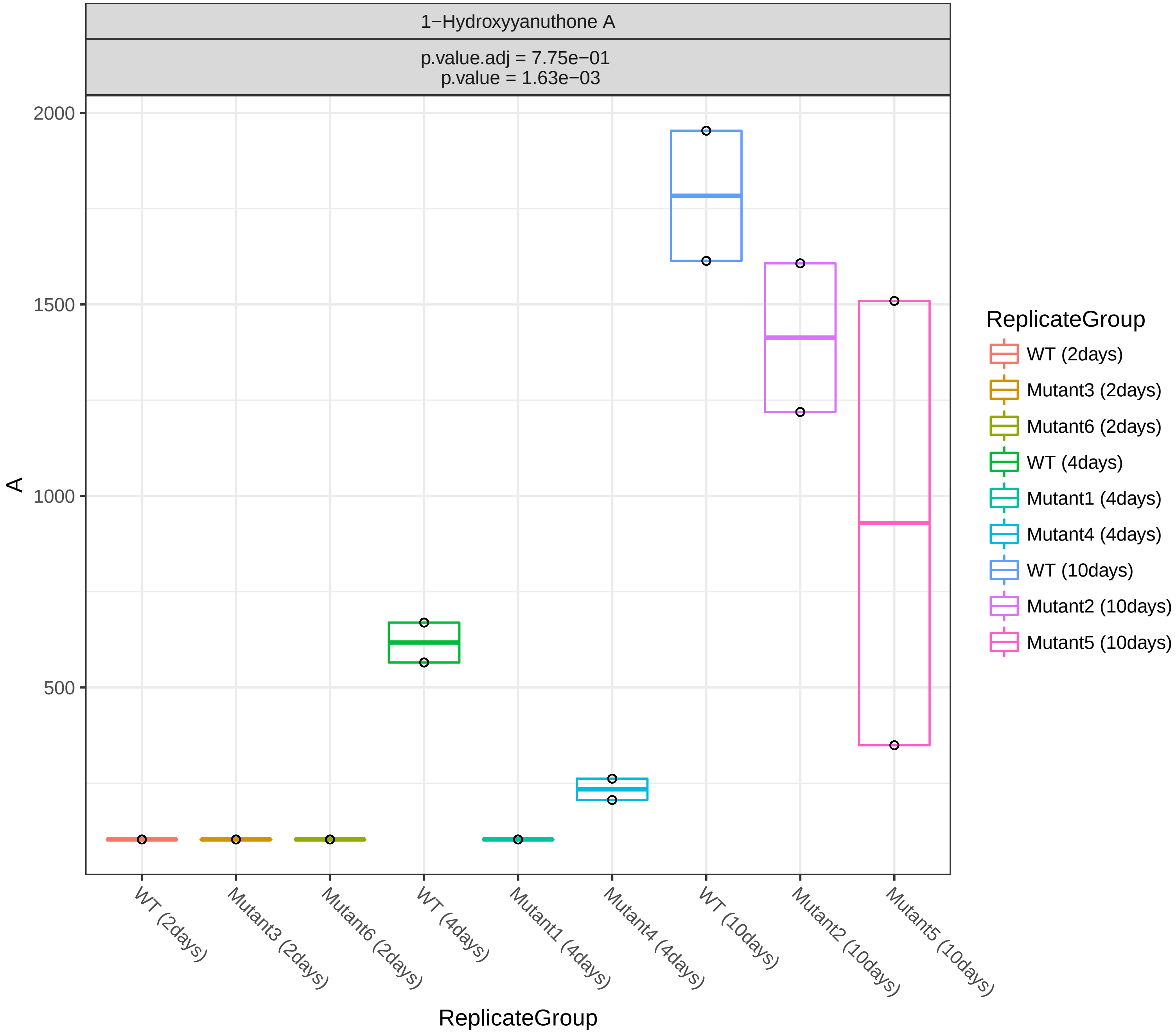
1−Hydroxyyanuthone A
p.value.adj = 7.75e−01
p.value = 1.63e−03
2000
1500
1000
500
A
WT (2days)
Mutant3 (2days)
Mutant6 (2days)
WT (4days)
Mutant1 (4days)
Mutant4 (4days)
WT (10days)
Mutant2 (10days)
Mutant5 (10days)
ReplicateGroup
ReplicateGroup
WT (2days)
Mutant3 (2days)
Mutant6 (2days)
WT (4days)
Mutant1 (4days)
Mutant4 (4days)
WT (10days)
Mutant2 (10days)
Mutant5 (10days)

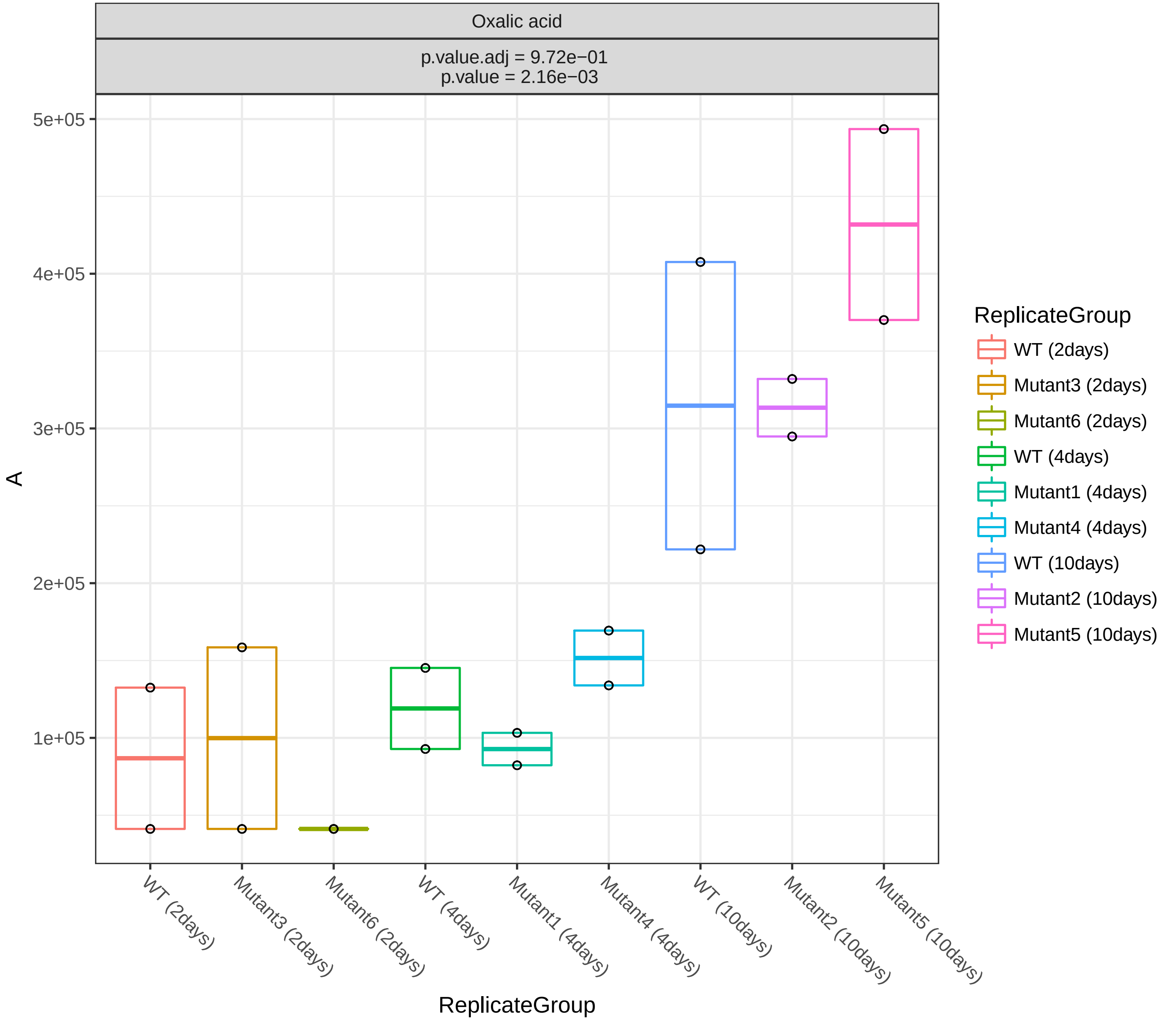
Oxalic acid
p.value.adj = 9.72e−01
p.value = 2.16e−03
5e+05
4e+05
3e+05
2e+05
1e+05
A
WT (2days)
Mutant3 (2days)
Mutant6 (2days)
WT (4days)
Mutant1 (4days)
Mutant4 (4days)
WT (10days)
Mutant2 (10days)
Mutant5 (10days)
ReplicateGroup
ReplicateGroup
WT (2days)
Mutant3 (2days)
Mutant6 (2days)
WT (4days)
Mutant1 (4days)
Mutant4 (4days)
WT (10days)
Mutant2 (10days)
Mutant5 (10days)

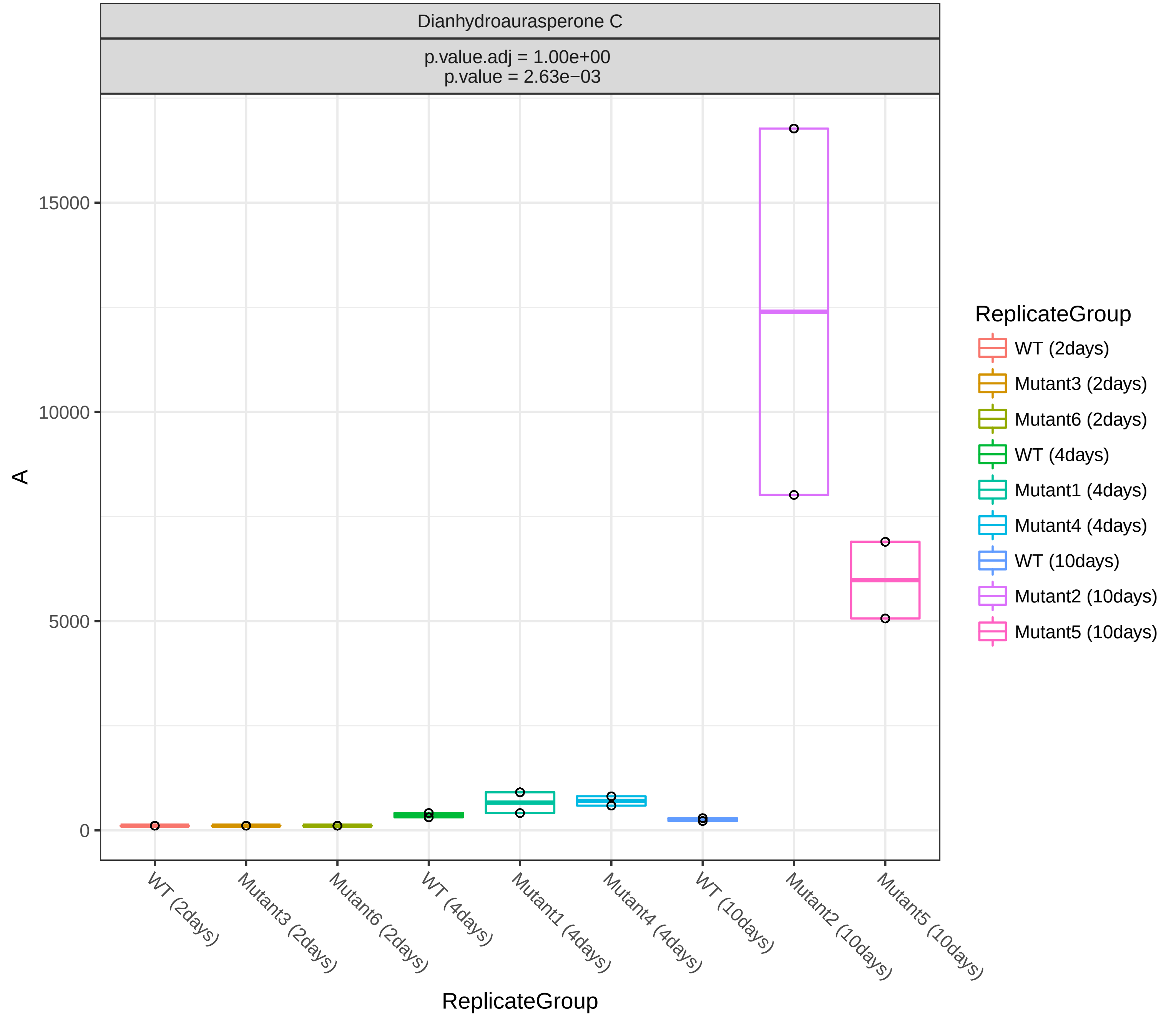
Dianhydroaurasperone C
p.value.adj = 1.00e+00
p.value = 2.63e−03
15000
10000
5000
0
A
WT (2days)
Mutant3 (2days)
Mutant6 (2days)
WT (4days)
Mutant1 (4days)
Mutant4 (4days)
WT (10days)
Mutant2 (10days)
Mutant5 (10days)
ReplicateGroup
ReplicateGroup
WT (2days)
Mutant3 (2days)
Mutant6 (2days)
WT (4days)
Mutant1 (4days)
Mutant4 (4days)
WT (10days)
Mutant2 (10days)
Mutant5 (10days)

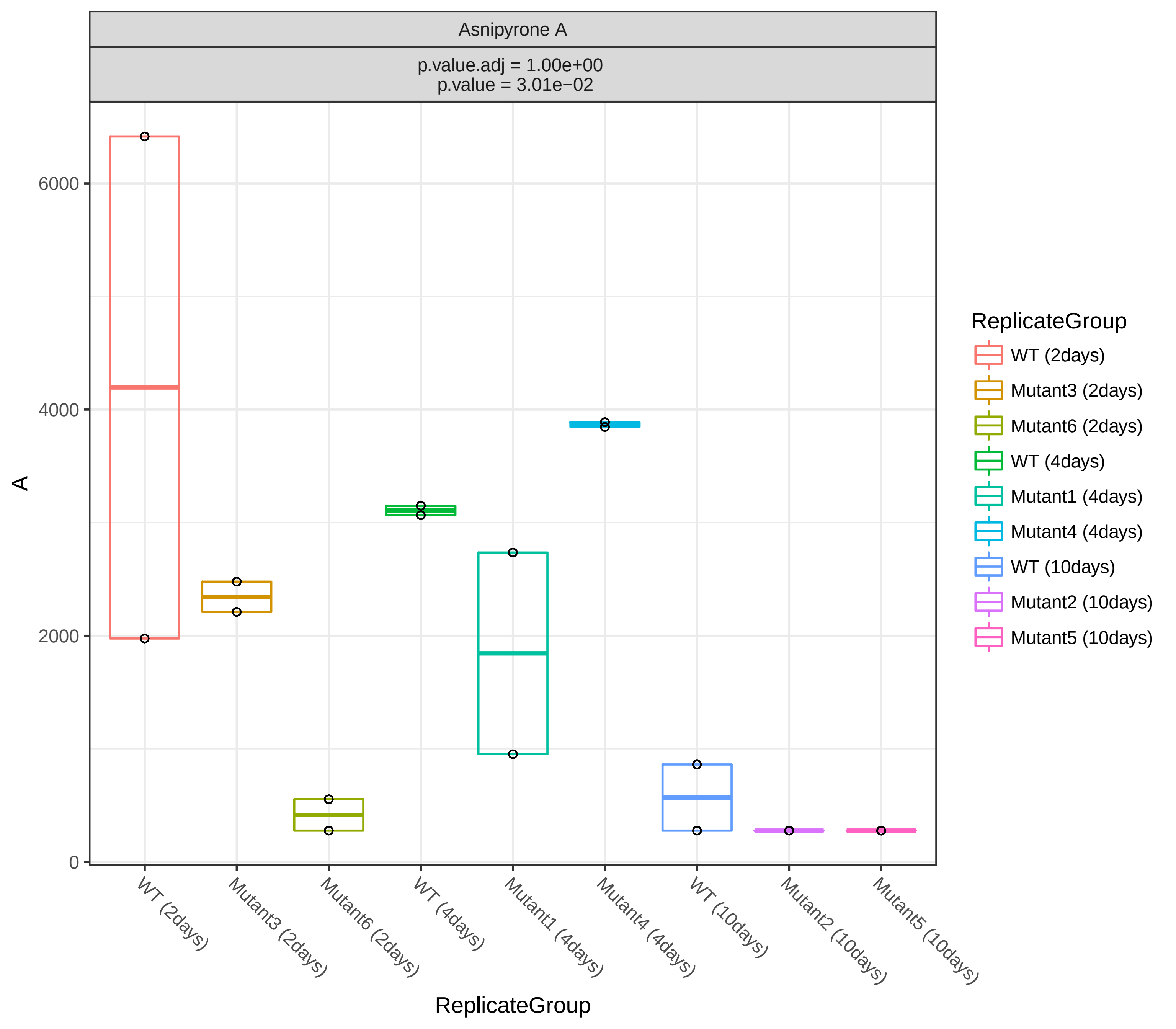
Asnipyrone A
p.value.adj = 1.00e+00
p.value = 3.01e−02
A
0
2000
4000
6000
ReplicateGroup
WT (2days)
Mutant3 (2days)
Mutant6 (2days)
WT (4days)
Mutant1 (4days)
Mutant4 (4days)
WT (10days)
Mutant2 (10days)
Mutant5 (10days)

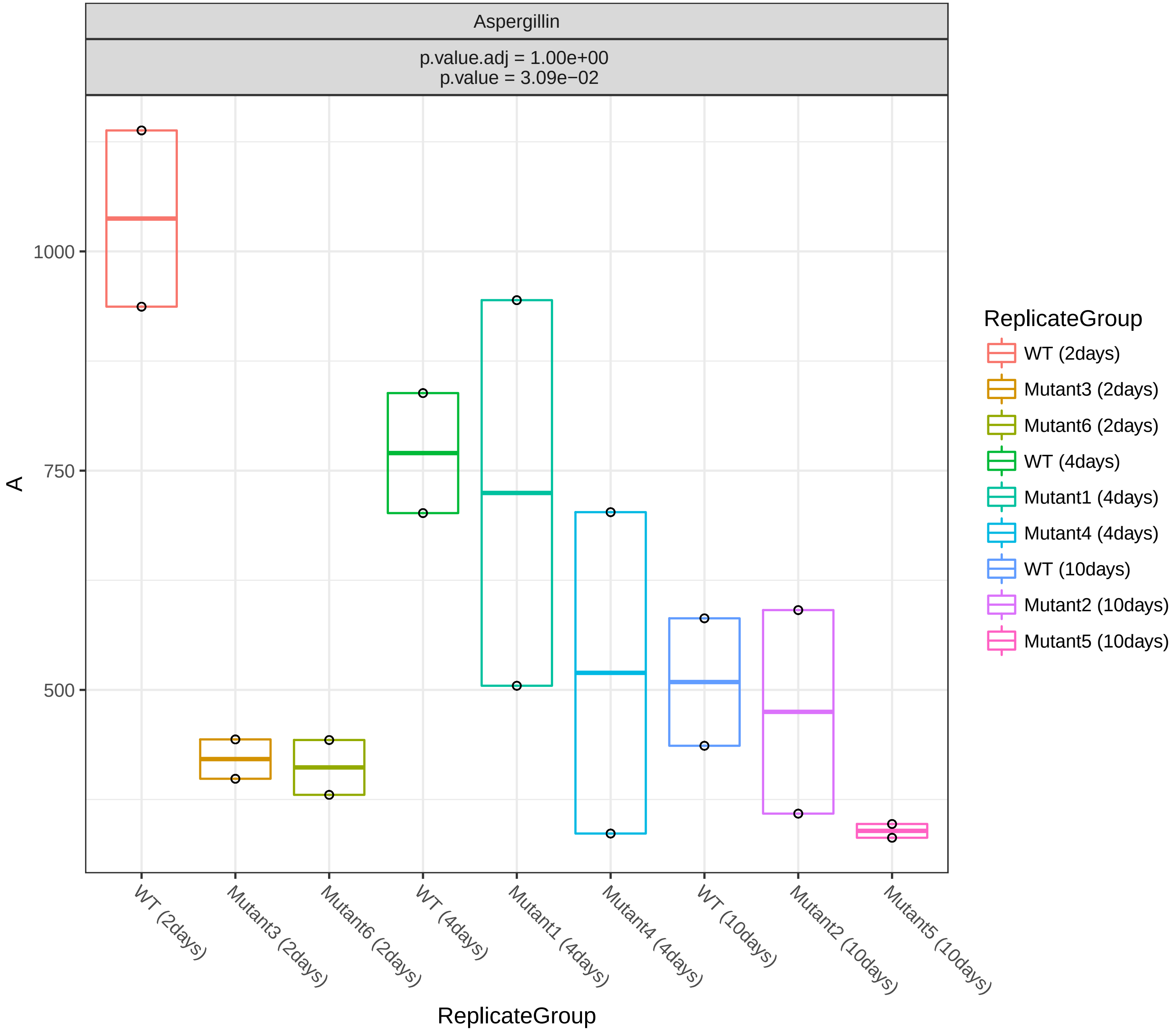

Aspergillin
p.value.adj = 1.00e+00
p.value = 3.09e−02
1000
750
500
A
WT (2days)
Mutant3 (2days)
Mutant6 (2days)
WT (4days)
Mutant1 (4days)
Mutant4 (4days)
WT (10days)
Mutant2 (10days)
Mutant5 (10days)
ReplicateGroup
ReplicateGroup
WT (2days)
Mutant3 (2days)
Mutant6 (2days)
WT (4days)
Mutant1 (4days)
Mutant4 (4days)
WT (10days)
Mutant2 (10days)
Mutant5 (10days)

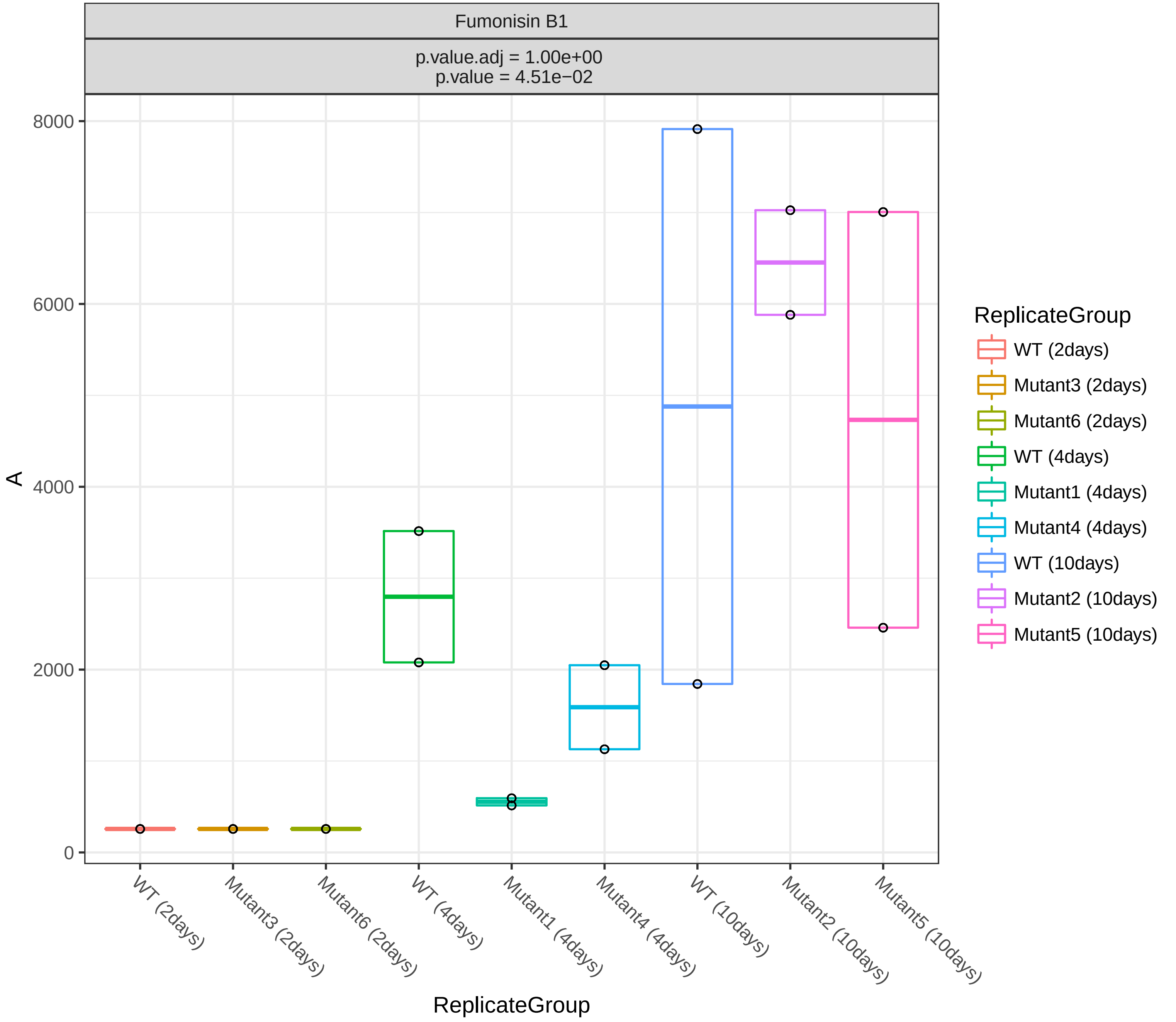
Fumonisin B1
p.value.adj = 1.00e+00
p.value = 4.51e−02
A
8000
6000
4000
2000
0
WT (2days)
Mutant3 (2days)
Mutant6 (2days)
WT (4days)
Mutant1 (4days)
Mutant4 (4days)
WT (10days)
Mutant2 (10days)
Mutant5 (10days)
ReplicateGroup
ReplicateGroup
WT (2days)
Mutant3 (2days)
Mutant6 (2days)
WT (4days)
Mutant1 (4days)
Mutant4 (4days)
WT (10days)
Mutant2 (10days)
Mutant5 (10days)

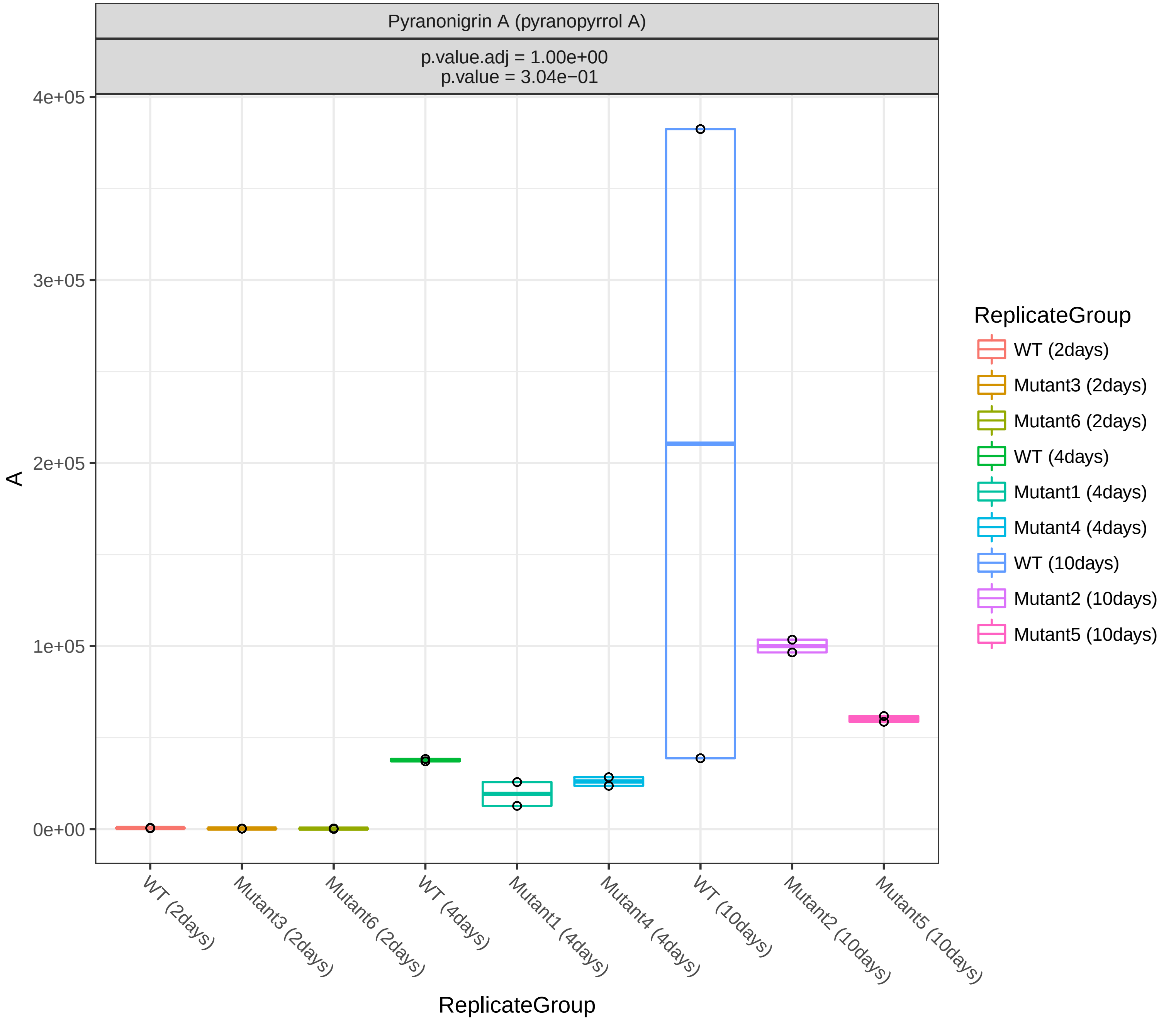

Pyranonigrin A (pyranopyrrol A)
p.value.adj = 1.00e+00
p.value = 3.04e−01
A
4e+05
3e+05
2e+05
1e+05
0e+00
WT (2days)
Mutant3 (2days)
Mutant6 (2days)
WT (4days)
Mutant1 (4days)
Mutant4 (4days)
WT (10days)
Mutant2 (10days)
Mutant5 (10days)
ReplicateGroup
ReplicateGroup
WT (2days)
Mutant3 (2days)
Mutant6 (2days)
WT (4days)
Mutant1 (4days)
Mutant4 (4days)
WT (10days)
Mutant2 (10days)
Mutant5 (10days)
